# Mapping and reprogramming human tissue microenvironments with MintFlow

**DOI:** 10.1101/2025.06.24.661094

**Authors:** Amir Akbarnejad, Lloyd Steele, Daniyal J. Jafree, Sebastian Birk, Marta Rosa Sallese, Koen Rademaker, Adam Boxall, Benjamin Rumney, Catherine Tudor, Minal Patel, Martin Prete, Stanislaw Makarchuk, Colin Y.C. Lee, Jonas Maaskola, Tong Li, Heather Stanley, April Rose Foster, Kenny Roberts, Andrew L. Trinh, Carlo Emanuele Villa, Giuseppe Testa, Satveer Mahil, Arash Mehrjou, Catherine Smith, Sattar Vakili, Menna R. Clatworthy, Omer Ali Bayraktar, Thomas Mitchell, Muzlifah Haniffa, Mohammad Lotfollahi

**Author notes:** Shared first authors (Amir Akbarnejad, Lloyd Steele, Daniyal J. Jafree, Sebastian Birk, Marta Rosa Sallese).

## Abstract

Tissue microenvironments reprogram local cellular states in disease, yet current computational spatial methods remain descriptive and do not simulate tissue perturbation. We present MintFlow, a generative AI algorithm that learns how the tissue microenvironment influences cell states and predicts how tissue perturbations can reprogram them. Applied to three human diseases, MintFlow uncovered distinct pathogenic spatial reprogramming in inflammatory and tumor microenvironments. In atopic dermatitis, MintFlow identified a novel, spatially-imprinted, type 2 (*IL13*^+^*ITGAE*^+^) epidermal T resident memory cell population (type 2 T_RM_), and decoded signaling pathways within the perivascular lymphoid niche. In melanoma, MintFlow identified fibrotic stroma resembling keloid scar tissue. In kidney cancer, MintFlow resolved immunosuppressed CD8^+^ T cell states within tertiary lymphoid structures. Furthermore, MintFlow enabled *in silico* perturbations of disease-relevant cell states and tissue environments. Regulatory T cell modulation in atopic dermatitis was predicted to suppress the pro-inflammatory tissue environment, supporting manipulation of these cells as a therapeutic target. In kidney cancer, in silico T cell replacement recapitulated immune checkpoint blockade, while spatially targeted macrophage depletion reverted immunosuppressed T cell states. The corresponding gene programs correlated with survival in large kidney cancer patient cohorts. Together, these findings position MintFlow as a tool for unbiased disease mechanism prediction and in silico perturbation, accelerating translational hypothesis generation and guiding therapeutic strategies.

## Introduction

Recent advances in single-cell and spatial transcriptomics are reshaping biomedical research by enabling the dissection of tissue architecture and cell identity with unprecedented resolution. These technologies allow researchers to map how specific cell types, transcriptional programs, and cell-cell interactions contribute to health and disease in their native spatial contexts. Such spatially resolved insights are expected to inform next-generation diagnostics, uncover context-specific therapeutic targets, and support the design of precision therapies including immunotherapies and regenerative strategies^1^. Understanding how cellular behavior is shaped within intact tissues is especially critical in complex and multifactorial diseases, where pathological outcomes often arise not only from changes within individual cells, but also from disrupted interactions between cells and their surrounding microenvironment. For instance, spatially restricted immunosuppressive signaling networks in tumors can drive immune evasion and resistance to therapy^2,3^, spatial gradients of chemokines can sustain chronic inflammation in skin diseases such as atopic dermatitis^4^, and localized glial–neuronal signaling has been implicated in the propagation of neuroinflammation in Alzheimer’s disease^5^. A key challenge to understanding such diseased tissue ecosystems is to distinguish intrinsic gene programs that define cellular identity from the effects induced by the surrounding microenvironment.

Computational approaches to resolve cell-intrinsic from microenvironment-induced cell states coupled with the ability to perform *in silico* perturbations to model and virtually reprogram tissue ecosystems would mark a paradigm shift in biomedical discovery. These opportunities move beyond existing descriptive analyses of pathological states, towards predicting disease mechanisms and potential targets to revert disease states. Such capabilities would not only aid the identification of cellular drivers of disease, but also support patient stratification, prognostic modeling, and the prioritization of therapeutic targets based on their spatial and contextual effects. Existing computational methods to model the effect of tissue microenvironment either fail to disentangle intrinsic and microenvironment-induced variation^6–9^, lack mechanistic interpretability and require post-hoc interpretation of embeddings^6,10,11^, are not flexible due to linear assumptions^12–14^, do not scale well^10,13,14^, or require prior knowledge about gene-gene interactions^15–17^ (detailed comparison in Supplementary Note 1). Furthermore, existing methods fall short in prioritizing microenvironments and genes to make novel biological hypotheses, and in enabling *in silico* perturbations to test these hypotheses or generate new ones. Crucially, filling the latter gap promises to enable hypothesis-driven, context-aware precision medicine via in silico perturbations and biological hypotheses generation.

To address the unmet need for modeling how the tissue microenvironment shapes cell identity and clinical outcomes, we present MintFlow, a generative AI framework based on flow matching (Fig. 1 and Methods) with theoretically grounded solution (identifiability proof in Methods and Supplementary Note 2) that disentangles intrinsic from microenvironment-induced gene expression in spatial transcriptomics data. Without relying on prior knowledge, predefined domains, or ligand–receptor pairs, MintFlow learns microenvironment-induced gene programs (MGP) at single-cell resolution and enables *in silico* perturbation of the tissue microenvironment, simulating how changes in local cellular context alter gene expression. Applied across spatial datasets from atopic dermatitis, melanoma, and clear cell renal cell carcinoma (ccRCC), MintFlow uncovered spatially organized, disease-specific cell states. In atopic dermatitis, Mintflow identified novel, microenvironment-induced, type 2 epidermal resident memory T (T_RM_) cells. *In silico* perturbation of skin microenvironments revealed cellular therapy with regulatory T cells (Tregs) as a potentially valuable therapeutic strategy in atopic dermatitis. In melanoma, MintFlow identified fibrotic immune-excluding stroma and angiogenic tumor progenitors; in ccRCC, simulated macrophage depletion reprogrammed exhausted T cell states and stratified survival in large cohorts. These findings position MintFlow as a generative AI model for uncovering spatially encoded drivers of disease and virtually modelling microenvironmental interventions, facilitating recreating these features using *in vitro* and *in vivo* models. We envision MintFlow as a step toward AI-driven construction of virtual tissue models, accelerating the discovery of spatial biomarkers, guiding therapeutic design and, ultimately, improving the clinical translation of spatial biology for patient care.

**Fig. 1.**
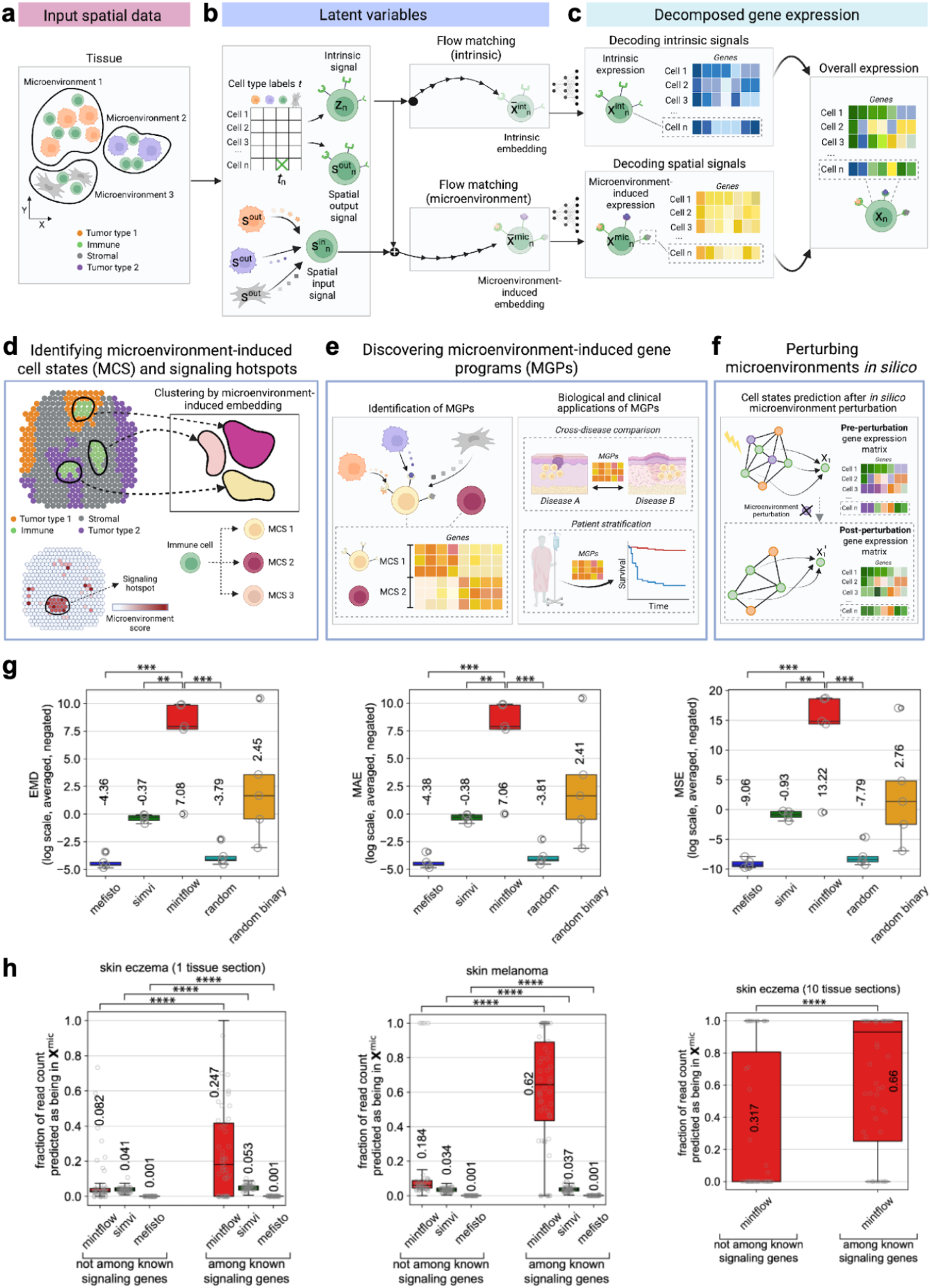
Overview and benchmarking of MintFlow. **a**, Given tissue sections profiled with single-cell resolution spatial transcriptomics, each cell’s microenvironment is determined based on spatial coordinates (Methods). Cell type labels and each cell’s microenvironment cell type composition (MCC) serve as supervision for MintFlow. **b**, MintFlow infers three embedding vectors that respectively encode intrinsic signals (*z*_*n*_), incoming spatial signals from the microenvironment 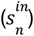, and outgoing spatial signals to the microenvironment 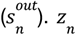 and 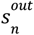 are generated with priors conditioned on cell type labels, while 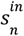 aggregates outgoing spatial signals of the cell’s microenvironment (*s*^*out*^). Using flow matching, *z*_*n*_ and 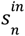 are transformed into intrinsic 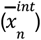 and microenvironment-induced 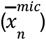 embeddings, which are decoded to generate intrinsic and microenvironment-induced read counts, respectively; together, these reconstruct the original read counts. **d**, Clustering microenvironment-induced embeddings enables identification of fine-grained microenvironment-induced cell states (MCS). A microenvironment score highlights signaling hotspots (Methods). **e**, Users can leverage MintFlow’s inferred MCS and read counts to identify microenvironment-induced gene programs (MGPs), compare between conditions, and stratify patients. **f**, MintFlow enables analysis of the effects on MCS and MGPs after perturbing microenvironments. **g**, MintFlow outperforms alternative methods in read count disentanglement on simulated data. MAE, mean absolute error; EMD, earth mover’s distance; MSE, mean squared error. Metrics are negated such that higher values are better. **h**, Using real data, MintFlow assigns a greater proportion of read counts of known signaling genes to the microenvironment-induced component compared to alternative methods (Methods). Counts < 20 were filtered. Only MintFlow was scalable enough to be applied to 10 tissue sections in skin eczema. Box plot elements are defined as center line, median; box limits, upper and lower quartiles; whiskers, 1.5× interquartile range. Four, three, two, and one asterisks respectively indicate p-values less than or equal to 0.0001, 0.001, 0.01, and 0.05 in a t-test.

## Results

### MintFlow learns microenvironment-induced cell states and predicts tissue perturbations

MintFlow is a deep generative model that learns the effects of spatial context on a cell’s gene expression by explicitly modeling incoming and outgoing cellular communication signals as latent variables. MintFlow learns to generate gene expression data and can disentangle it into components driven by intrinsic variability and spatial context, respectively, enabling identification of cell states that are influenced by their microenvironment. By learning a joint representation of each cell’s intrinsic identity and its tissue context, MintFlow is a powerful tool to improve our understanding of how the local tissue environment reprograms cellular behavior (Fig. 1a–c, Methods).

In addition to disentangling intrinsic and microenvironment-induced effects, MintFlow introduces the concept of tissue perturbation. By simulating changes in the environment, such as cell deletion or replacement, MintFlow predicts how these alterations influence gene expression, offering valuable insights into how microenvironmental shifts impact cells. This capability enables large-scale hypothesis generation and testing, providing new avenues for exploring the dynamic interplay between cells and their surrounding environments.

MintFlow encodes each cell using the gene expression and cell type label of the cell itself, as well as those of cells within the local tissue microenvironment. Precisely, a multi-stage encoder learns a cell’s intrinsic latent features, conditioned on its cell type label, and microenvironment-induced latent features, conditioned on the composition of cells within its tissue microenvironment (i.e. its microenvironment cell type composition (MCC)). These latent features are decoded into corresponding gene expression vectors: 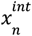 and 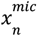, respectively, using flow matching (Methods). The sum of these expression vectors reconstructs the original expression vector.

MintFlow decomposition enables multiple downstream applications. Clustering the microenvironment-induced latent features captures fine-grained microenvironment-induced cell states (MCS; Fig. 1d and Methods). Moreover, it enables the calculation of a “microenvironment score”, quantifying the relative influence of microenvironment signals on each cell to identify spatial hotspots of signaling activity (Fig. 1d and Methods). Microenvironment-induced gene expression can be analyzed with standard single-cell software such as scanpy^18^ to derive differentially expressed genes and identify microenvironment-induced gene programs (MGPs) (Fig. 1e). Additionally, MintFlow enables *in silico* tissue perturbation simulating deletion or replacement of any cell within a tissue microenvironment, and predicts the resultant gene expression following perturbation, enabling large scale hypotheses generation and evaluation (Fig. 1f and Methods).

We benchmarked MintFlow’s capability to predict microenvironment-induced gene expression against alternative methods and random baselines on simulated data. For the latter, we simulated known microenvironment-induced effects to provide ground truth to evaluate model performance (Supplementary Fig. 1 and Methods). MintFlow significantly outperformed alternatives in predicting microenvironment-induced gene expression based on three different metrics (Fig. 1g, Supplementary Figs. 2 and 3, and Methods). We also compared MintFlow’s performance on real data, including single-sample Xenium data from one tissue section of atopic dermatitis and melanoma, respectively, and a multi-sample Xenium dataset of ten atopic dermatitis tissue sections (Fig. 1h and Methods). MintFlow proportions of microenvironment-induced read counts were on average higher in previously reported signaling genes^19^ compared to other genes. This is notable because many of these signaling genes are known to act on neighboring cells, implying that their expression is more likely to be driven by microenvironmental cues rather than intrinsic cellular programs. In contrast, alternative methods either assigned a greater fraction of signaling gene read counts to intrinsic expression or attributed only a minimal proportion to microenvironment-induced activity, thereby underestimating the role of external signals. Together, these findings indicate that MintFlow successfully captures microenvironment-induced gene expression and outperforms alternative methods in distinguishing extrinsic from intrinsic transcriptional influences.

### MintFlow identifies a microenvironment-induced T_RM_ cell state and T cell activation hub in atopic dermatitis

We first applied MintFlow on a dataset of 10 human skin tissue sections from 8 individuals with atopic dermatitis profiled by Xenium 5k, including 8 newly generated sections (Methods). Atopic dermatitis is a T-cell mediated chronic inflammatory skin disease characterized by type 2 inflammation^4^, an immune response associated with anti-helminth (parasite) responses and cytokines such as IL-4 and IL-13. We included both inflamed (lesional) and phenotypically normal (non-lesional) skin samples. After quality control (Methods), we obtained 197 487 cells.

MintFlow identified regions with high microenvironment scores, consistent with high signaling activity, in the superficial dermis of the skin (Fig. 2a). Some of these regions correlated with areas of perivascular inflammatory infiltrate on H&E (Fig. 2a), which is a histopathological feature of disease. To further understand the MintFlow-predicted high signalling regions, we clustered microenvironment-induced gene expression to identify 10 distinct microenvironments in the skin across samples (Fig. 2b, Supplementary Fig. 4a and Methods). One specific cluster (*T_DC)* was prominent in the superficial dermis, where high microenvironment scores were evident (Fig. 2b). The *T_DC* domain was predominantly formed of T cells, with additional antigen-presenting cells (APCs) such as migratory dendritic cells (*MigDCs*) and type 2 conventional dendritic cells (*cDC2*) (Supplementary Fig. 4b). The *T_DC* domain was expanded in inflamed skin compared to non-inflamed skin (Fig. 2c), suggesting a potential pathogenic role.

**Fig. 2.**
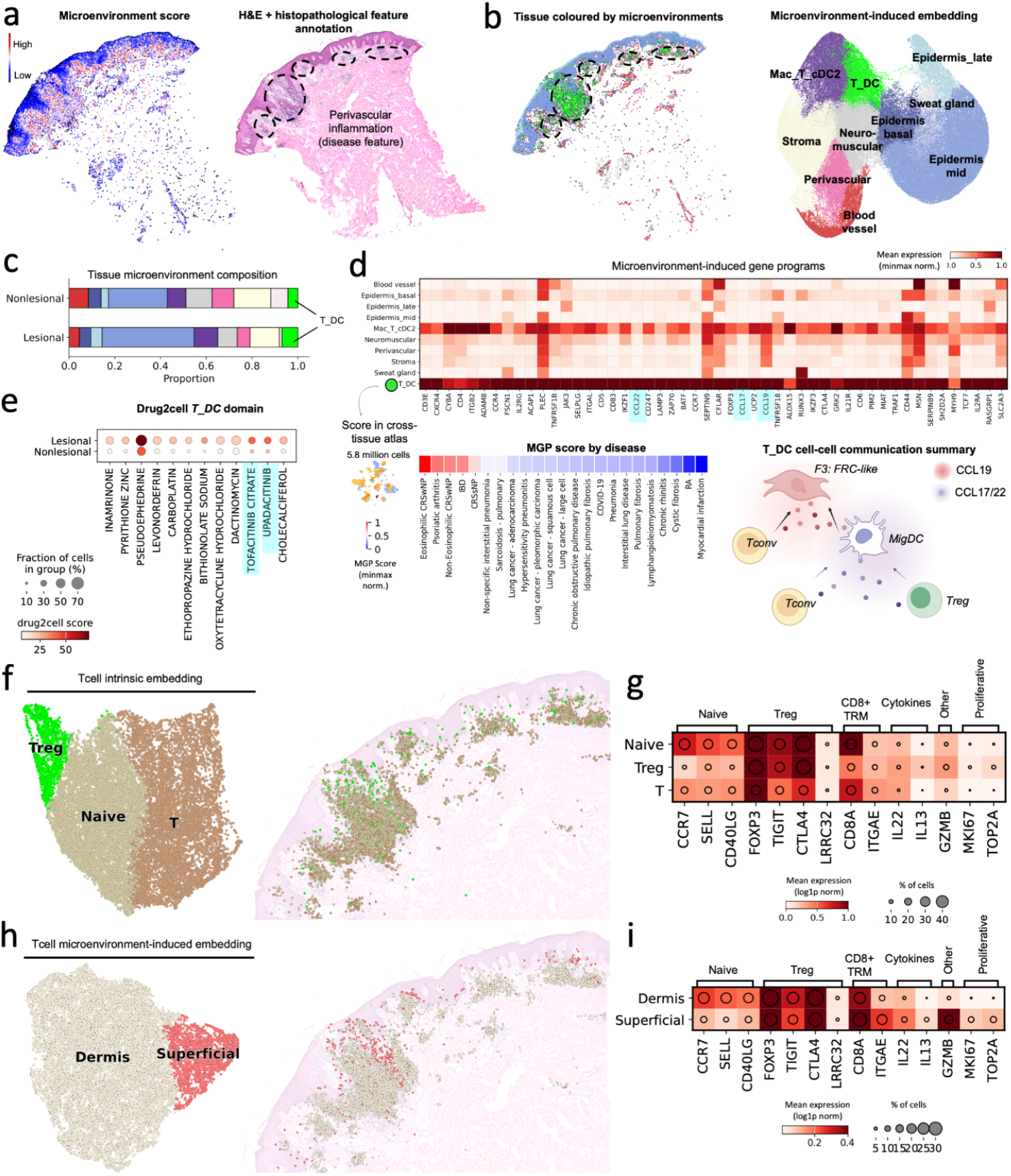
MintFlow identifies a microenvironment-induced T_RM_ cell state and T cell activation hub in atopic dermatitis. **a**, Tissue section colored by microenvironment score and corresponding H&E slide with annotated histopathological features of the same section profiled by Xenium 5k. **b**, Tissue section colored by microenvironment domains and UMAP visualization of the microenvironment-induced embedding. **c**, Tissue composition by microenvironment domain in inflamed and non-inflamed skin. **d**, MGP for *T_DC* domain (top). Gene module scores of the *T_DC* MGP in a cross-tissue atlas (bottom left, Methods). Genes marked in blue are specifically highlighted in the main text. Schematic of active signalling in the *T_DC* MGP (bottom right). **e**, Drug2cell results for genes in the *T_DC* domain for all drugs. **f**, UMAP visualization of intrinsic expression matrix for T cells (left) and location of cells in inflamed tissue (right). **g**, Combined heatmap and dot plot for selected T cell markers for intrinsic clusters. **h**, UMAP visualization of microenvironment-induced expression matrix for T cells (left) and location of cells in inflamed tissue (right). **i**, Combined heatmap and dot plot for selected T cell markers for microenvironment-induced clusters. H&E, Haematoxylin and eosin. MGP, Microenvironment-induced gene program.

We used MintFlow to derive MGPs for the *T_DC* domain. The *T_DC* MGP included genes involved in T cell activation and homing (*ITGB2, SELPLG*)^20^, T cell interactions with APCs (*ICAM3, ITGAL* (encodes LFA-1))^21,22^, type 2 immunity polarization (*IL21R, ALOX15*)^23^, and chemotactic positioning of immune cells (*CCL19, CCL22*) (Fig. 2d). We performed cell-cell communication analysis using the *T_DC* MGP to elucidate active signaling pathways within this region. This analysis highlighted a CCL19/CCR7 axis, in which *F3: FRC-like* and *F2/3: Perivascular* fibroblasts recruit *MigDCs* and *T cells*, as previously reported in inflamed skin^24^. In addition, we identified a *CCL22/CC17-CCR4* axis for *MigDC*-*T* (*Tnaive, Treg, T helper* (*Th*)) interactions (Fig. 2d and Supplementary Fig. 4c). Importantly, *MigDC-T* cell interactions are crucial to generating immune responses^25^. Using pathway analysis we identified that the most enriched process for the *T_DC* MGP was T cell activation (Supplementary Fig. 4d and Methods). These results suggest that the *T_DC* microenvironment represents a T cell activation hub within human skin.

Identification of the *T_DC* MGP enabled us to query potential new drug targets using a computational pipeline (drug2cell)^26^. This analysis correctly identified drugs with known efficacy in atopic dermatitis (upadacitinib (a Janus kinase (JAK) inhibitor), prednisolone (a steroid))^27^, in addition to potentially novel drugs (Fig. 2e and Supplementary Fig. 4e). For example, Amlexanox is used for treating aphthous ulcers and has evidence for a PDE4 inhibitor action^28^, which is an FDA approved topical therapy for atopic dermatitis^29^. Pseudoephedrine is a sympathomimetic drug with vasoconstriction, and possibly anti-inflammatory, actions, which may potentially be beneficial in atopic dermatitis^30^.

Next, we sought to (1) validate the *T_DC* MGP in an external single-cell RNA sequencing (scRNA-seq)^31^ dataset of patients with atopic dermatitis, and (2) investigate its cross-disease relevance using an integrated dataset of 23 different diseases across 6 human tissues (skin, gut, lung, synovium, nasal mucosa, and heart)^24^. For atopic dermatitis scRNA-seq data, the *T_DC* MGP was more highly expressed in inflamed compared to non-inflamed skin in an independent cohort of 4 patients^31^ (Supplementary Fig. 4f and Methods), consistent with a role in T cell activation. For the cross-tissue comparison (Fig. 2d and Methods), the *T_DC* MGP gene module score was highest in eosinophilic chronic rhinosinusitis with nasal polyps (CRSwNP). This is a notable finding, as CRSwNP is a disease of type 2 immunity affecting the nasal mucosa, and dupilumab, a monoclonal antibody targeting IL-4 and IL-13 type 2 immunity signaling, is efficacious in both atopic dermatitis and eosinophilic CRSwNP^32^. We also identified high scores for inflammatory bowel disease (IBD) and psoriatic arthritis, which like atopic dermatitis can be treated with JAK inhibitors. This suggests a role for the *T_DC* domain in T cell activation across tissues, potentially reflecting a similar lymphoid follicular-like niche across human inflammatory diseases^33^.

Given the role of T cells in atopic dermatitis pathology^29^, we next used MintFlow to disentangle T cell gene expression. This enabled us to dissect the T cell subtype distribution into microenvironment-induced and intrinsic characteristics of T cells. For the intrinsic latent space, we identified distinct clusters for naive (*CCR7, SELL*) and regulatory (*FOXP3, TIGIT, CTLA4*) genes (Fig. 2f,g). In addition, clustering by microenvironment-induced gene expression revealed a transcriptionally distinct *CD8A*^+^*ITGAE*^+^*GZMB*^+^ T cell cluster with increased expression of *IL13, IL22*, and *GZMB* that was located in superficial skin (Fig. 2h,i). Transcriptomically and spatially, this T cell phenotype was characteristic of epidermal T_RM_s^34^, which are known to have a spatially-imprinted phenotype across human tissues^35^. *ITGAE* and *GZMB* were identified to be spatially imprinted genes in CD8+ T cells in recent work in mouse intestinal tissue^36^, suggesting that Mintflow correctly identified a microenvironment-induced pathogenic T cell state in atopic dermatitis skin.

### *In silico* spatial perturbations model cellular therapy in atopic dermatitis

We next used MintFlow to perform *in silico* spatial perturbations of the *T_DC* MGP in the inflamed atopic dermatitis skin microenvironment. Specifically, we performed *in silico* depletion/augmentation of Tregs, as Treg modulation is being actively explored as a cellular therapy in immune-mediated diseases^37,38^. In the first perturbation, we removed Tregs from the microenvironment (Treg depletion). In the second perturbation, we replaced all T_conv_ cells (i.e. other T populations) with Tregs (Treg augmentation). We performed each *in silico* perturbation in the same tissue region (Fig. 3a) and assessed the impact on remaining immune cells within the perturbed tissue (T_conv_ cells and MigDCs) using differential gene expression testing and pathway analysis (Methods).

**Fig. 3.**
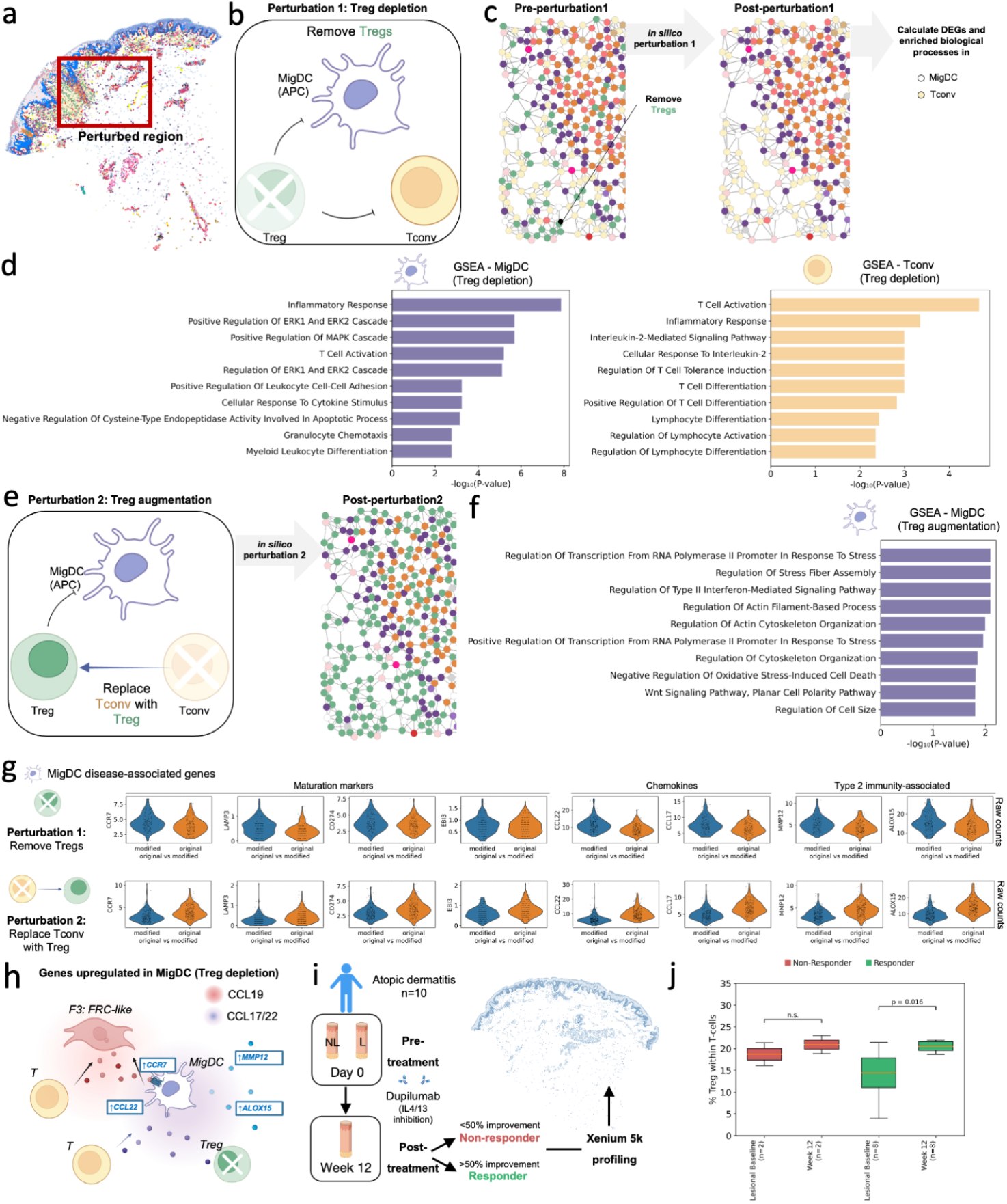
MintFlow *in silico* perturbation highlights therapeutic value of Tregs in atopic dermatitis. **a**, Tissue region used for perturbation. **b**, Schematic representation of perturbation 1 (Treg depletion). **c**, Zoomed-in perturbation region to show changes before and after applying perturbation 1 (Treg depletion). **d**, Gene set enrichment analysis (GSEA) using Gene Ontology (GO) biological processes in MigDCs and T_conv_ under perturbation 1 (Treg depletion). **e**, Schematic representation of perturbation 2 (Treg augmentation). **f**, Gene set enrichment analysis using GO biological processes in MigDCs under perturbation 2 (Treg augmentation). **g**, Violin plots of gene expression for migration and activation markers of MigDCs under perturbation 1 and 2. **h**, Schematic representation of perturbation 1 (Treg depletion), highlighting the roles of upregulated genes in MigDC migration and recruitment of T cells. **i**, Schematic of data collected post-treatment. NL, nonlesional; L, lesional. **j**, Box plot of percentage of Treg cells in tissue section before and after treatment by treatment response by donor. For each response group, we performed a paired, two-sided Wilcoxon signed-rank test comparing lesional baseline vs week 12 Treg%. Sample sizes are indicated on the x-axis.

When Tregs were removed from the microenvironment (Fig. 3b,c), MigDCs upregulated genes involved in inflammatory response (rank 1), regulation of MAPK cascade (rank 3), T cell activation (rank 4), positive regulation of lymphocyte cell-cell adhesion (rank 6), and myeloid-leukocyte differentiation (rank 10) (Fig. 3d). Interestingly, MAPK is reported to be important for DC activation and chemotaxis^25^, suggesting increased MigDC activation. Amongst the top 10 DEGs in MigDCs were the chemokines *CCL17* and *CCL22* (Supplementary Fig. 5a), which we previously identified in the *T_DC* MGP to be linked to attraction of T cells (Fig. 2d), including Tregs (Supplementary Fig. 4c). T_conv_ cells upregulated genes involved in T cell activation (rank 1), inflammatory response (rank 2), and interleukin-2 mediated signaling (rank 3) (Fig. 3d)^29,32^. Overall, the *in silico* perturbation of Treg depletion suggested an increased inflammatory state, and thus likely a worsened atopic dermatitis phenotype. In agreement with this predicted effect, individuals with Treg dysfunction are known to develop severe atopic dermatitis due to hyperinflammatory T cells in IPEX (immune dysregulation, polyendocrinopathy, enteropathy, X-linked) syndrome^39^.

Next, we investigated the effect of Treg augmentation by replacing all T_conv_ cells with Tregs (Fig. 3e). In this setting, inflammatory pathways were not enriched in MigDCs (Fig. 3f and Supplementary Fig. 5b). Instead, the top biological processes related to cytoskeleton modification, changes in cell size, and stress response. Together, this suggests that MintFlow correctly predicted pro-inflammatory effects in Treg depletion, whereas Treg augmentation was not associated with pro-inflammatory pathways.

We next queried whether Mintflow could predict the direction of change in expression of genes linked to active inflammation in MigDCs. In addition to chemokines secreted by MigDCs (*CCL17, CCL22*), we assessed migration/maturation markers (*CCR7, LAMP3, CD274, EBI3*)^40^ and type 2 immunity-related genes expressed by dendritic cells (*MMP12, ALOX15*)^4,41^ (Fig. 3g). In Treg depletion (perturbation 1), these genes were increased, consistent with an active inflammatory state. In Treg augmentation (excess Tregs), these genes were downregulated (Fig. 3g). These results, and the spatial MGP (Fig. 2d), suggest that Tregs modulate skin inflammation in atopic dermatitis and that Treg abundance is associated with an anti-inflammatory (treatment) effect (Fig. 3h).

We next used *in vivo* data from 10 atopic dermatitis patients 12 weeks post-dupilumab treatment to investigate our *in silico* MintFlow predictions (Fig. 3i). We measured changes in Treg abundance by clinical response after treatment (Methods). In responders (>50% improvement from baseline score), Tregs typically increased with treatment from baseline to week 12 (Fig. 3j). In 2 non-responders (<10% improvement), skin Tregs did not show a significant change and were typically elevated in baseline samples compared to responders (Fig. 3j), raising the possibility that differential Treg abundance in inflamed skin influences clinical response to dupilumab treatment. Overall, our findings support the hypothesis that Treg modulation may be a potentially valuable therapeutic strategy in atopic dermatitis, influencing dendritic cell activation, and that differences in Treg profiles could influence treatment response.

### MintFlow identifies inducible keloid-like stroma in cutaneous melanoma

Cell signaling is crucial in mediating interactions between cancer cells, immune cells, and the non-immune stroma in solid tumors^42^. To evaluate MintFlow’s ability to identify microenvironment-induced effects in TME cells, we applied it to a public human cutaneous melanoma dataset profiled using 5000-plex Xenium^43^ (Methods). After quality control, we identified 98,749 cells.

The highest microenvironment scores were found outside of tumor nests (Fig. 4a,b). We identified 8 distinct microenvironment clusters and their MGPs, including two clusters formed predominantly of tumor cells, which we termed *Melanoma1* and *Melanoma2* (Fig. 4c-e and Supplementary Fig. 6a). The cancer-predominant *Melanoma1* and *Melanoma2* microenvironments occupied spatially-distinct regions in the skin, with *Melanoma2* forming a smaller, deeper nodule (Fig. 4f). While known melanoma marker genes (e.g. *MLANA, PRAME*) were present throughout both melanoma regions, the *Melanoma1* MGP uniquely contained genes that were not clearly evident in the *Melanoma2* domain (e.g. *EDNRB, CST1, BAALC, GDF15*) (Fig. 4e,g). Pathway analysis using these MGPs showed enrichment of biological processes of amino acid transport and glycogen catabolism for the *Melanoma1* domain, and cell migration and microtubule polymerization for *Melanoma2* (Supplementary Fig. 6b and Methods). This result suggested functionally distinct regions within the melanoma tumor.

**Fig. 4.**
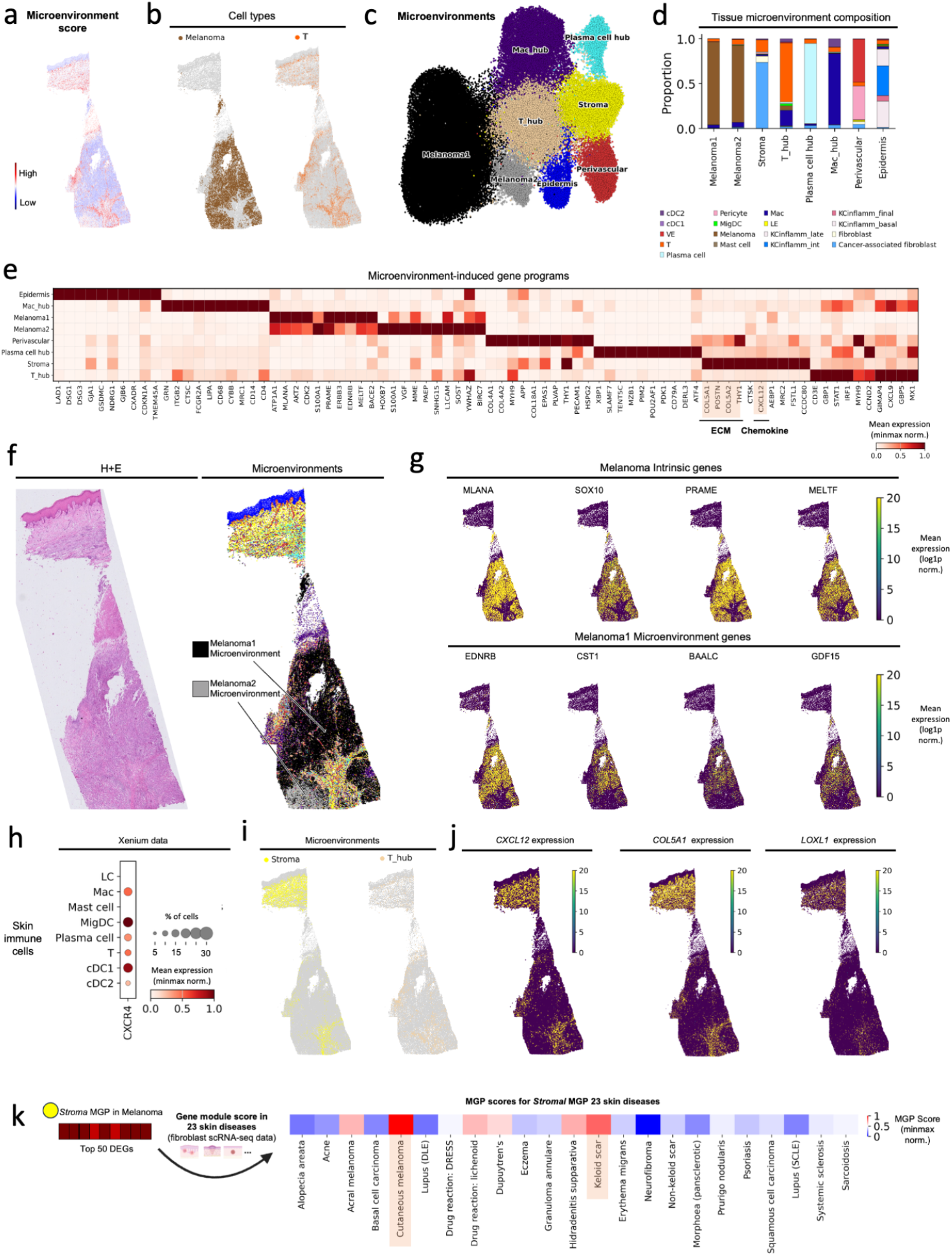
Deciphering tumor microenvironment immune cell segregation in melanoma using MintFlow. **a**, Melanoma tissue section colored by microenvironment score. **b**, Tissue section colored by cell type (melanoma vs other and T cell vs other). **c**, UMAP visualization of the microenvironment-induced embedding for cutaneous melanoma. **d**, Cellular composition of microenvironment domains. **e**, Microenvironment-induced gene programs (MGPs) for each microenvironment domain. **f**, H&E-stained image of the tissue section from cutaneous melanoma and corresponding Xenium section colored by microenvironment domain. **g**, Differentially expressed genes for melanoma cells (intrinsic) (top row) and the Melanoma1 microenvironment domain (bottom row). **h**, Expression of CXCR4, the receptor for the CXCL12 ligand, by skin immune cells (Methods). **i**, Melanoma tissue section colored by microenvironment domains (Stroma and T_hub). **j**, Melanoma tissue section colored by selected genes from the *Stroma* MGP. **k**, Gene module scores for the *Stroma* MGP across fibroblasts from different skin diseases profiled using scRNA-seq data (Methods).

Immunotherapy is widely used in the treatment of advanced melanoma to promote anti-tumor T cell-mediated responses and can achieve sustained remission. Recent work has linked immunotherapy response to the state of cancer-associated fibroblasts (CAFs) in the stroma^44^. We therefore interrogated the *Stroma* microenvironment domain identified by MintFlow, which was formed predominantly of CAFs and T cells (Fig. 4d), to understand how this microenvironment influences T cell activity.

The *Stroma* MGP included genes encoding *CXCL12* and extracellular matrix (ECM) proteins (Fig. 4e). *CXCL12* is a chemoattractant for CXCR4-expressing cells, which include T cells and other immune cells (Fig. 4h). An inducible *CXCL12-CXCR4* axis has been reported to exclude T cells from tumor nests in human cancers^45,46^. Consistently, T cells in melanoma were largely located in the *Stroma* or *T_hub* domains with high CXCL12 expression (Fig. 4b,e,i-j).

In addition to the chemotactic axis, T cells can be segregated away from tumor cells by physical barriers, including collagen fibres^42^. ECM genes in the *Stroma* MGP encoded collagens (*COL4A2, COL5A1, COL5A2, COL11A1*), as well as enzymes that cross-link collagen fibrils (*LOX, LOXL1, LOXL2*) (Fig. 4e,j and Supplementary Fig. 6c). Collagen cross-linking increases the stability of ECM fibres and is associated with worse outcomes in cancer^42^. Overall, our results point towards physical and chemotactic exclusion of T cells from melanoma, leading to a tumor-permissive environment.

To further investigate the nature of the stroma in melanoma, we scored the melanoma *Stroma* MGP across a dataset of fibroblasts from 23 different skin diseases profiled using scRNA-seq^24^ (Methods). We found that the highest expression of the *Stroma* MGP was in cutaneous melanoma, confirming the relevance of the MGP across different technologies (Fig. 4k). Interestingly, we also observed high expression for the melanoma *Stroma* MGP in keloid scarring, which is a fibroproliferative scar with dense stroma that is characterized by increased cross-linking of collagen fibres^47^. Consistent with a microenvironment-induced scar-like stroma, the term “stromagenesis” has been used in cancer studies to describe bi-directional stromal signaling pathways that activate resident fibroblasts to a proliferative state^48^, suggesting this inducible signaling network is captured by Mintflow.

When assessing fibroblasts by intrinsic and spatial profiles, we observed a distinct superficial fibroblast population adjacent to the epidermis (Supplementary Fig. 6d), consistent with *F1: Superficial* fibroblasts that we recently reported (Supplementary Fig. 6e)^24^. Of note, a synergistic interplay between superficial dermal fibroblasts and basal epithelial cells has been reported to reciprocally maintain cellular identity^49^, consistent with a spatially-imprinted profile.

In summary, our results suggest that active microenvironment signaling contributes to an inducible chemotactic and physical barrier against tumor cell infiltration in cutaneous melanoma, including an inducible keloid-like stroma.

### MintFlow identifies a tertiary lymphoid hotspot consisting of distinct T cell states in kidney cancer

To further evaluate the ability of MintFlow to resolve microenvironment-induced cell states within the TME, we applied it to spatial transcriptomic data from a patient with ccRCC, the most common form of adult kidney cancer. Three macroscopically distinct regions of the tumor core, as well as the tumor–normal interface where the cancer abuts non-neoplastic kidney tissue, were profiled using the 5000-plex Xenium platform (Fig. 5a; patient identifiers within Methods^50^). After integration and annotation, the dataset comprised 337,116 cells across 24 transcriptionally distinct cell types. MintFlow was applied to the integrated dataset to generate a latent space representing microenvironment-induced gene expression (Fig. 5b), achieving successful batch integration (Supplementary Fig. 7a) and robust alignment across cell types (Supplementary Fig. 7b). Spatial mapping of predicted microenvironment-induced cell states revealed regional variation in microenvironment score across the tissue (Fig. 5c), potentially reflecting intratumoral heterogeneity within ccRCC. Within the tumor cores, discrete hotspots of elevated microenvironment score were observed, with a prominent structure within one tumor core (Fig. 5c; core 2).

**Fig. 5.**
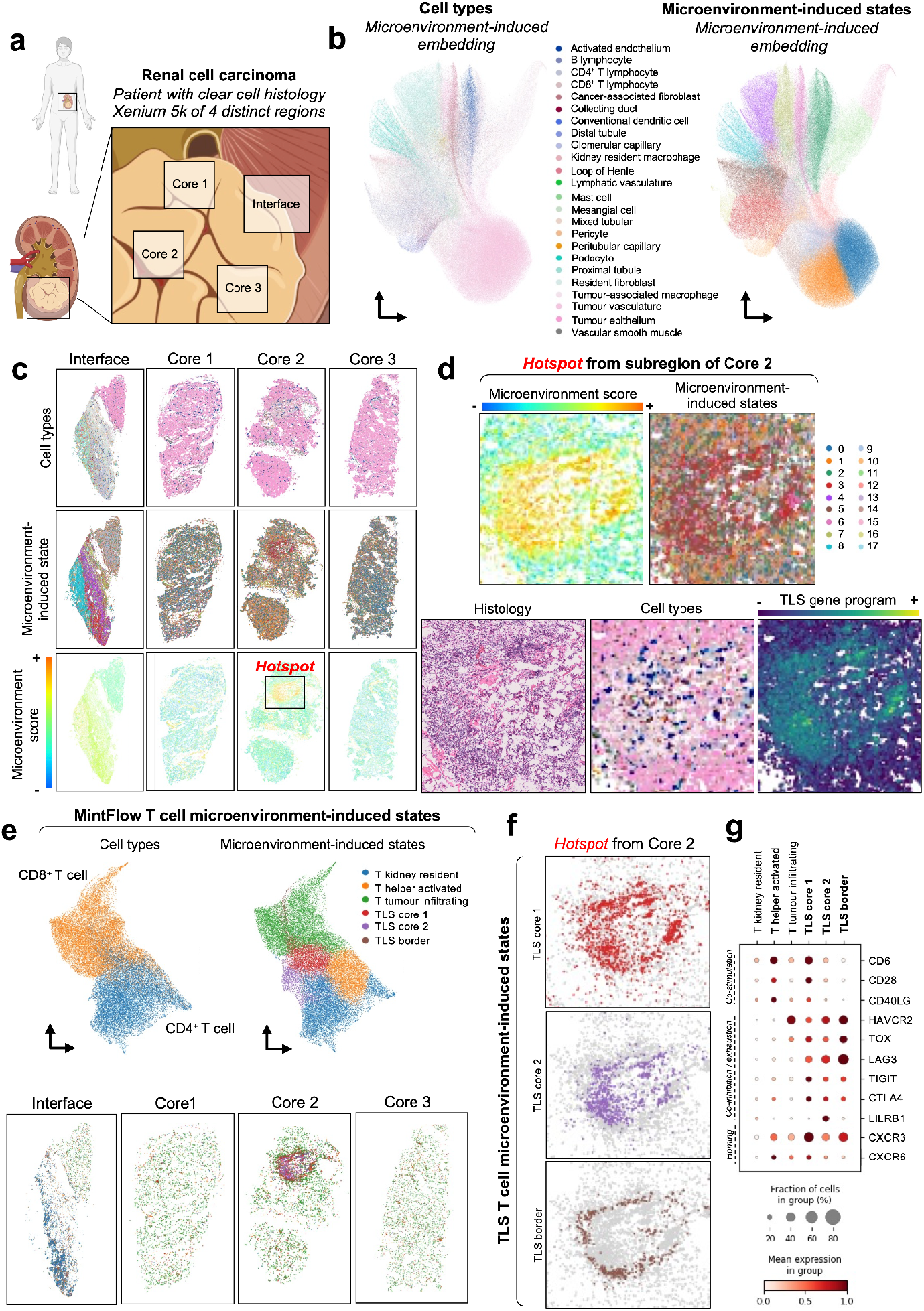
MintFlow analysis of microenvironment-induced cell states in kidney cancer. **a**, Schematic illustrating the spatial transcriptomic profiling strategy in a nephrectomy specimen from a patient with clear cell renal cell carcinoma (ccRCC). Four anatomically distinct tumor regions were sampled: Core 1, Core 2, Core 3, and the tumor–normal interface **b**, UMAP visualizations of MintFlow-inferred microenvironment-induced gene expression profiles, colored by cell type (left) and MintFlow clusters (right). **c**, Spatial mapping of each region showing cell types (top row), MintFlow microenvironment-induced clusters (middle row), and computed microenvironment score (bottom row). A signaling hotspot is highlighted in Core 2. **d**, High-resolution profiling of the hotspot region displaying microenvironment score, MintFlow clusters, histological image, annotated cell types, and a tertiary lymphoid structure (TLS) module score. The TLS score was calculated using the following genes: *CXCL13, CCR7, LTA, CXCR5, CD8A, CD4, CCL19, CD79A, MS4A1*, and *CD3E*. **e**, Subclustering of T cells using MintFlow: UMAPs showing original cell type annotations (left) and MintFlow-derived microenvironment-induced T cell states (right), alongside spatial distribution of these states across the four tissue regions (below). **f**, Magnified view of the TLS region showing spatial localization of MintFlow-inferred T cell states: TLS Core 1, TLS Core 2, and TLS Border T cells. **g**, Dot plot of differentially expressed genes across MintFlow-inferred TLS T cell states. Markers of immune inhibition and exhaustion (e.g., *HAVCR2, LAG3*) are enriched in tumor-infiltrating and TLS-resident T cells, while *CXCR3*, encoding the receptor for CXCL10 and CXCL11, is specifically enriched in TLS T cell subsets.

Histopathological examination of this tumor core revealed a dense lymphocytic aggregate (Fig. 5d), and MintFlow clustering indicated a predominance of T cells within the hotspot. Given its architecture and cellular composition, we hypothesized this structure represented a tertiary lymphoid structure (TLS), ectopic lymphoid aggregates known to arise in chronic inflammation and cancer, and previously associated with clinical outcome in ccRCC^51–53^. To validate this, we computed a TLS gene signature score based on expression of canonical TLS-associated genes (including CXCL13, LTA, and CXCR5)^54^, confirming significant enrichment within the hotspot compared to surrounding tissue (Fig. 5d).

We then sought to leverage MintFlow to further dissect the immune architecture within the TLS beyond the capability of conventional spatial transcriptomics analysis. We subsetted MintFlow-generated embeddings for T cells and performed unsupervised clustering. This analysis revealed six distinct microenvironment-induced T cell states, three of which localized specifically within the TLS (Fig. 5e). These TLS-associated states, which we termed TLS Core 1, TLS Core 2, and TLS Border T cells, occupied spatially distinct domains within the aggregate (Fig. 5f). These microenvironment-induced states were not recoverable either by clustering MintFlow’s intrinsic embeddings (Supplementary Fig. 8a) or by fine-grained reclustering of the original dataset (Supplementary Fig. 8b), highlighting MintFlow’s unique ability to resolve spatially modulated T cell programs.

To define the microenvironment-induced T cell gene programs, we performed differential expression analysis, which revealed that each gene program harbored distinct molecular signatures (Fig. 5g). Consistent with previous observations in ccRCC^55,56^, tumor-infiltrating T cells exhibited enrichment for exhaustion markers such as *HAVCR2, TOX, LAG3*, and were expressed at the highest level by TLS Border T cells. Tumor-infiltrating T cells were also enriched for *CXCR3*, with its cognate ligands, *CXCL9* and *CXCL10*, spatially enriched within the TLS (Supplementary Fig. 9). MintFlow identified similar TLS-associated T cell states across Xenium data of additional ccRCC patients, with MGPs mapping to lymphoid aggregates observed on histology (Supplementary Fig. 10). These findings demonstrate MintFlow’s ability to uncover microenvironment-induced immune architecture not readily detectable by conventional analyses, highlighting T cell states that are enriched within the TLS in ccRCC.

### MintFlow reveals inhibitory macrophage-T cell crosstalk within the TLS of kidney cancer

Having identified distinct microenvironment-induced T cell states within the TLS, we next sought to determine whether these populations could be retrospectively identified in single-cell RNA sequencing (scRNA-seq) data lacking spatial information. We reanalyzed a published scRNA-seq dataset of ccRCC^50^, comprising 147,917 tumor-infiltrating cells from 10 patients, including the donor from whom the Xenium spatial transcriptomics data were derived (Fig. 6a). Using MintFlow-derived gene programs for the TLS Core 1, TLS Core 2, and TLS Border T cells, we projected these signatures onto the T cell compartment of the scRNA-seq dataset (Fig. 6a). Expression of the aggregated TLS T cell gene program was distributed across T cell subclusters without clear demarcation, highlighting the difficulty of identifying spatially organized immune states without explicit spatial resolution. Importantly, the TLS T cell programs were detectable across patients spanning low, intermediate and high Leibovich risk groups (Fig. 6b), supporting the generalizability of these states across clinical stages.

**Fig. 6.**
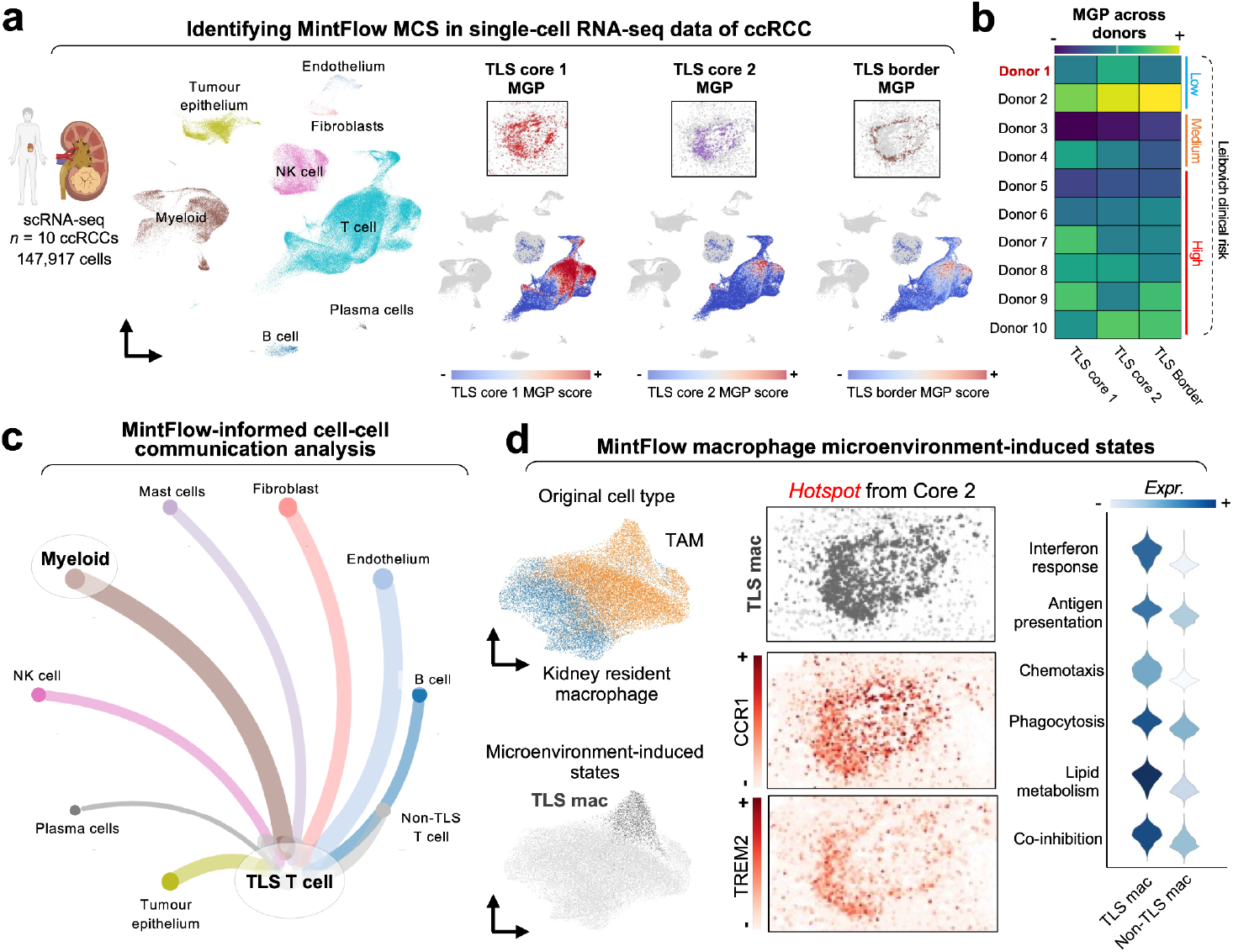
Application of MintFlow to uncover inhibitory macrophage-T cell crosstalk within the TLS of kidney cancer. **a**, Left: UMAP visualization of single-cell RNA sequencing (scRNA-seq) data from 10 patients with clear cell renal cell carcinoma (ccRCC), including one patient whose tumor was also profiled by spatial transcriptomics in this study. Cells are colored by broad cell type annotation. Right: Feature plots showing recovery of the MintFlow-derived TLS T cell microenvironment-induced gene programs within the scRNA-seq dataset. **b**, Heatmap displaying TLS T cell program scores across T cells from each patient in the scRNA-seq cohort, grouped by Leibovich grade: a clinical risk classification for ccRCC. **c**, Ligand–receptor interaction analysis using LIANA applied to the scRNA-seq data. TLS T cells were identified based on MintFlow program scores. The circle plot illustrates predicted interaction strength between TLS and non-TLS T cells and other major cell types. **d**, Subclustering of macrophages using MintFlow microenvironment-induced embeddings from the Xenium dataset. UMAPs show original macrophage annotations (top) and MintFlow-defined TLS macrophages (bottom). Spatial localization of TLS macrophages is shown alongside expression of key markers *TREM2* (middle) and *CCR1* (bottom). Violin plot indicates enrichment of immune regulatory pathways—antigen presentation, interferon response, and T cell co-inhibition, within TLS macrophages compared to all other macrophages.

We next aimed to independently validate the MintFlow-derived TLS Border T cell state in an *in vivo* setting. Mouse models of ccRCC are scarce, and as TLS formation is considered a pan-cancer phenomenon^54^, we analyzed a recently published study in which Kaede transgenic mice were injected subcutaneously with colorectal tumor epithelial cells, followed by treatment with either anti-PD-L1 antibody or isotype control^57^. This model employed photoconversion to distinguish tumor-resident from recently infiltrating cells, allowing spatial inference without direct imaging (Supplementary Fig. 11a). scRNA-seq of sorted CD8^+^ T cells revealed naïve, memory, cycling, and exhausted subsets (Supplementary Fig. 11b). While the TLS Core 1 and 2 gene programs were not enriched within CD8^+^ T cell populations, the TLS Border T cell program was selectively enriched in subsets of memory and exhausted CD8^+^ T cells. Among these, memory cells included both photoconverted and non-photoconverted cells, whereas exhausted cells were almost exclusively photoconverted (Supplementary Fig. 11c), consistent with chronic residency in the tumor microenvironment. Differential gene expression analysis of TLS Border-high exhausted CD8^+^ T cells revealed upregulation of immune activation and lymphoid-organizing genes, including *Il2rb, Stat1*, and *Ltb*, in anti-PD-L1–treated mice compared to controls (Supplementary Fig. 11d). These findings provide evidence that TLS Border T cells identified by MintFlow resemble infiltrating and immunosuppressed T cells observed within an independent model system, and suggest that this spatially constrained population is amenable to reprogramming by immune checkpoint inhibition.

Given the immunosuppressed phenotype observed in TLS T cells, and the detection of a similar exhausted and tumor-resident population in a mouse model that was sensitive to immune checkpoint inhibition, we hypothesized that specific cell–cell interactions within the TLS niche may drive their dysfunctional state. To explore this, we performed ligand–receptor interaction analysis across the previously published scRNA-seq dataset of ccRCC^50^, treating TLS T cells as a discrete population. Among all cell types in the ccRCC microenvironment, TLS T cells exhibited the highest predicted number of interactions with myeloid populations, particularly macrophages (Fig. 6c). Examination of the top 100 ligand–receptor pairs revealed enrichment for major histocompatibility complex (MHC) class I and II interactions, as well as co-inhibitory signals, including molecules such as *LAG3, HAVCR2*, and others known to mediate T cell exhaustion (Supplementary Fig. 12). These findings implicate macrophages as the dominant cell type exerting immune inhibitory activity upon T cells within the TLS.

We next returned to the human RCC spatial Xenium dataset to ascertain the presence and spatial localization of macrophages potentially responsible for inhibitory signaling within the TLS. Subclustering MintFlow’s macrophage latent space for microenvironment-induced gene expression resolved a distinct tumor-associated macrophage subset localized specifically to the TLS and characterized by high expression of *TREM2* and *CCR1* (Fig. 6d). Differential expression analysis of this TLS-associated macrophage subset revealed enrichment for pathways linked to antigen presentation, interferon response, and T cell co-inhibition, consistent with the predicted ligand–receptor interactions. Analysis of other stromal populations, including vasculature (Supplementary Fig. 13a) and fibroblasts (Supplementary Fig. 13b), revealed no distinct microenvironment-induced cell states specifically localized to the TLS, suggesting that macrophage-T cell interactions are the primary mediators of immune suppression within this microenvironment domain.

### *In silico* perturbations predict therapeutic potential of TLS macrophage targeting in kidney cancer

We next leveraged MintFlow’s generative modeling capabilities to perform in silico perturbation experiments in ccRCC. We curated a gene expression signature of T cells exposed to therapy from a previously published scRNA-seq study of ccRCC tumors before and after immune checkpoint blockade (ICB)^58^. Within our Xenium dataset of treatment-naive ccRCC, we identified a phenotypically equivalent T cell subset and virtually replaced the immunosuppressed TLS-resident T cells with these post-ICB-like cells (Fig. 7a,b). To assess the biological validity of this T cell virtual replacement, we assessed the MintFlow-predicted transcriptional changes in neighboring macrophages, utilizing a signature of genes shown to be expressed in ICB-exposed macrophages in ccRCC^58^. Of the genes upregulated within macrophages within ICB-treated scRNA-seq data, twelve genes were included in the Xenium 5K panel and were expressed in >10% of macrophages. Apart from one gene, the remaining eleven were upregulated by macrophages when TLS-resident T cells were replaced by post-ICB-like T cells (Fig. 7b,c and Supplementary Fig. 14a). These genes included *CXCL9* and *CXCL10*, the cognate receptor for which was expressed on TLS T cell subsets (Fig. 5g). Gene set enrichment analysis of the top differentially expressed genes within ICB-like macrophages included terms related to T cell or lymphocyte activation, regulation of phagocytosis and macrophage activation (Supplementary Fig. 14b). These results support the utility of MintFlow for simulating microenvironmental reprogramming with biologically grounded outcomes.

**Fig. 7.**
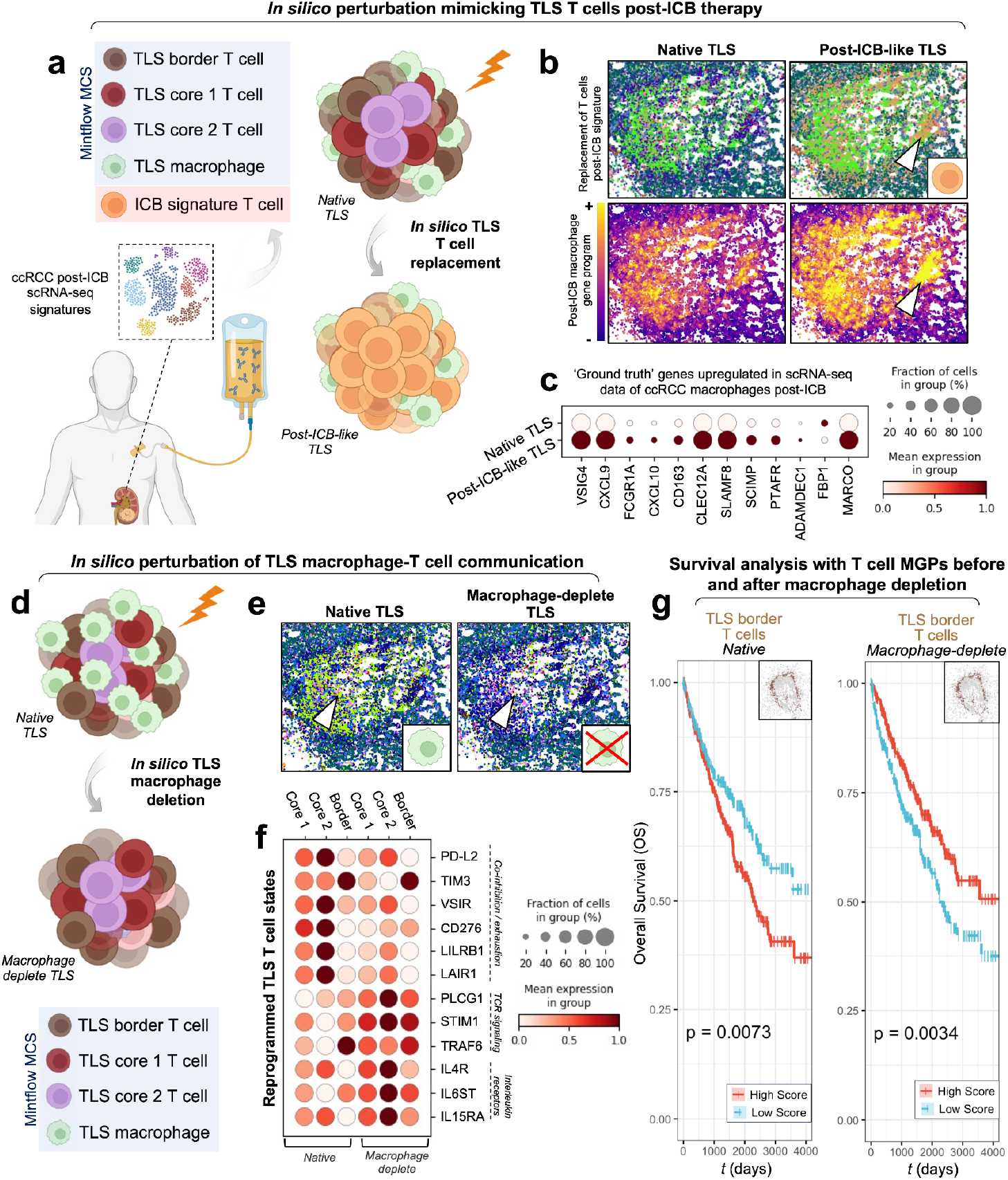
In silico perturbations of the TLS in kidney cancer. **a–c**, Assessing the biological validity of MintFlow’s *in silico* perturbation framework in ccRCC. A gene signature of T cells following immune checkpoint blockade (ICB) was curated from published scRNA-seq data (**a**), and an equivalent T cell subpopulation was identified in our Xenium dataset. Immunosuppressed TLS T cell states, as defined by MintFlow, were replaced with these post-ICB-like T cells (schematic, **a**; spatial visualization, **b**). A signature of macrophage-specific genes known to be upregulated following ICB, considered as ‘ground truth’, were plotted on the tissue, showing upregulation upon perturbation (**b**). Analysis of the individual genes using a dot plot (**c**) showed that 11 out of 12 of the ‘ground truth’ genes were upregulated upon virtual T cell replacement. **d–f**, In silico perturbation of TLS-associated macrophages using MintFlow. Schematic (**d**) and spatial plot (**e**) illustrate virtual macrophage removal. Dot plot (**f**) shows top differentially expressed genes in TLS T cells before and after macrophage deletion. Perturbation reduced expression of co-inhibitory and exhaustion markers (*PD-L2, TIM3, VSIR, CD276, LILRB1, LAIR1*), while increasing expression of TCR signaling molecules (*PLCG1, STIM1, TRAF6*) and interleukin receptors (*IL4R, IL6ST, IL15RA*). **g**, Kaplan–Meier survival analysis of 606 ccRCC patients from the TCGA cohort. Stratification by TLS Border T cell gene expression showed that high expression of the original (pre-perturbation) program was associated with worse survival (p = 0.0073), whereas the reprogrammed gene program predicted a survival benefit (p = 0.0034).

Having provided evidence for the biological validity of our *in silico* perturbation, we then sought to manipulate macrophage-T cell inhibition within the TLS of ccRCC. Specifically, we performed a perturbation in which the TLS-associated macrophages were deleted, predicting how T cell gene expression states would respond in the absence of suppressive myeloid signals (Fig. 7d,e). DEG analysis following macrophage deletion revealed consistent reprogramming across each microenvironment-induced TLS T cell state (Fig. 7f). Molecules associated with T cell co-inhibition and exhaustion, including *PD-L2, TIM3, VSIR, CD276, LILRB1* and *LAIR1*, were downregulated, whereas reprogrammed T cells exhibited increased expression of markers indicating TCR signaling (*PLCG1, STIM1, TRAF6*) and encoding interleukin receptors (*IL4R, IL6ST, IL15RA*). These findings suggest that local disruption of macrophage-mediated signals within the TLS restores an activated T cell phenotype *in silico*.

To assess the clinical significance of MintFlow-predicted reprogramming to a larger and more generalizable cohort of patients, we applied the microenvironment-induced gene programs to bulk RNA sequencing data from 606 ccRCC tumors in The Cancer Genome Atlas (TCGA) cohort. For each TLS T cell state, we computed gene expression scores across patients and stratified individuals into ‘high’ and ‘low’ expression groups. Kaplan–Meier survival analysis using available overall survival data revealed that neither the TLS Core 1, TLS Core 2 nor TLS macrophage gene programs were significantly associated with patient outcome (Supplementary Fig. 15). However, high expression of the TLS Border T cell program, representing the peripheral, suppressive T cell state within the TLS, was significantly associated with worse overall survival (p = 0.0073) (Fig. 7g). When we applied the MintFlow-inferred gene program for TLS Border T cells after macrophage removal *in silico*, high expression of the reprogrammed state was associated with a survival advantage (p = 0.0034). These findings suggest that targeted modulation of macrophage–T cell interactions within tertiary lymphoid structures could therapeutically reprogram local immunity and potentially improve patient outcomes in ccRCC.

## Discussion

Spatial transcriptomics offers unprecedented insight into tissue organization and how it is altered in disease. However, current computational approaches remain largely descriptive and do not resolve how the microenvironment actively shapes cellular gene expression, limiting their utility for mechanistic discovery, patient stratification and therapeutic prediction. Here, we introduce MintFlow, a deep generative AI model that learns how spatial context reprograms cell states. MintFlow identifies microenvironment-induced cell states and gene programs that underpin human pathology. Importantly, MintFlow’s *in silico* perturbation enables users to simulate how removal or replacement of specific cell types reconfigures local transcriptional states. Across multiple diseases, MintFlow not only recapitulates established spatial features but also uncovers previously unrecognized, clinically relevant cell states and programs dictated by the microenvironment in inflammatory and malignant diseased contexts. These findings demonstrate MintFlow’s potential as a predictive framework for spatial biology, with broad applications for virtual tissue reprogramming to guide biomarker discovery and drug repurposing.

A central strength of MintFlow lies in its ability to recover microenvironment-induced transcriptional programs that are often obscured in traditional analyses. Applied across three distinct diseases, MintFlow uncovered both disease-specific and generalizable spatial cell states. In atopic dermatitis, it identified a T cell–dendritic cell activation hub for type 2 immunity, conserved across other type 2–inflammatory diseases such as eosinophilic chronic rhinosinusitis CRSwNP. Mintflow also identified superficial *CD8*^+^*IL13*^+^*ITGAE*^+^ resident memory–like T cells as a microenvironment-induced state, consistent with recent murine findings^36^. While resident memory T cells have been well described in psoriasis and vitiligo^59,60^, they have not previously been implicated in atopic dermatitis^34^. Conversely, in melanoma, MintFlow resolved a fibroblast-rich, collagen-crosslinked stromal program expressing *CXCL12*, associated with T cell exclusion and resembling fibrotic keloid tissue. Finally, in ccRCC, MintFlow prioritized TLS as a hotspot of signaling activity, composed of three spatially distinct T cell states. This included an immunosuppressed border population not detectable by conventional clustering, distinguished by a gene program associated with poor survival in a large and heterogeneous cohort of patients.

A defining feature of MintFlow is its capacity for *in silico* perturbation of the tissue, leveraging generative AI to predict how modifying the microenvironment alters gene expression. In atopic dermatitis, MintFlow simulated the depletion or augmentation of Treg cells, recapitulating known biology: Treg depletion amplified inflammatory signaling, consistent with FOXP3-mutant IPEX syndrome, while Treg augmentation did not show elevated inflammatory pathways. Treg cellular therapy is being actively explored in other immune-mediated diseases such as IBD, with proof-of-concept that Treg therapies reverse disease activity^39^. In ccRCC, MintFlow identified macrophage–T cell interactions within TLS as key mediators of local immune suppression. Simulated deletion of TLS-associated macrophages led to downregulation of co-inhibition and exhaustion markers in TLS T cells, and instead, reprogrammed them toward more activated states. When *in silico* reprogrammed T cell states were projected onto TCGA ccRCC bulk tumor profiles, they became associated with a survival advantage, demonstrating that MintFlow goes beyond spatially resolving cell–cell interactions but also links these to clinically meaningful outcomes. These findings position MintFlow as a powerful framework for uncovering spatial drivers of cellular dysfunction and simulating therapeutic rewiring of diseased tissue microenvironments.

Many of the perturbations explored by MintFlow, such as cell type-specific depletion within intact tissue architecture, are challenging or unfeasible to perform at scale using *in vitro* models or animal systems, highlighting the unique value of virtual experimentation in guiding mechanistic and translational hypotheses. Of known candidates identified by MintFlow, several have already shown utility in translational settings. For example, of the TLS T cell genes downregulated by *in silico* macrophage deletion in ccRCC, *LAIR1* is upregulated in RCC and its expression correlates with poor survival^61^, *TIM3* expression is associated with response to immunotherapy^62^ and LILRB1 inhibitors are undergoing early phase clinical trials in kidney cancer patients^63^. We also tested the sensitivity of MintFlow tissue perturbations to the number of cells altered in the tissue and measured the effect size change in response to the perturbations. We observed that as a larger subpopulation of target cells are perturbed, the expression of the considered genes also changed incrementally (Supplementary Fig. 16). We further performed an assessment of biological validity of our perturbation in ccRCC, demonstrating that replacement of T cells with a subset resembling those treated with ICB predicted altered gene expression within macrophages in line with genes known to be upregulated in ccRCC macrophages upon ICB treatment^58^.

Although MintFlow introduces a novel approach for modeling microenvironment-induced cell variation and simulating *in silico* perturbations, several limitations remain. One includes sensitivity to low read counts (Supplementary Fig. 2 and 3) and the need for consistent gene panels across tissue sections. While the flow matching process is computationally intensive during training, our implementation incorporates a neighbor sampling strategy to enhance scalability, enabling training the model on multiple tissue sections encompassing millions of cells (Methods). MintFlow is susceptible to segmentation errors produced by current technologies, but we anticipate segmentation errors to be reduced by future advances in technology and novel segmentation methodologies. Future work could expand the modeling framework beyond multiple tissue sections to support training on multi-tissue spatial atlases, enabling transfer learning across datasets and analysis of microenvironmental influences at atlas scale. Additionally, while we have released MintFlow as an open source Python package, training MintFlow can be challenging and requires dataset-specific fine-tuning of hyperparameters due to its complex modeling architecture. To support adoption, we provide detailed documentation and guidance at https://mintflow.readthedocs.io.

We envision MintFlow as a transformative tool for spatial transcriptomics analysis in both basic research and clinical applications. Its ability to disentangle spatial drivers of gene expression, simulate cellular perturbations, and predict outcomes, positions it as a platform for therapeutic hypothesis generation, drug response modeling and patient stratification. By integrating MintFlow into spatial workflows, researchers and clinicians can gain mechanistic insight into disease architecture and virtually test interventions. This framework has the potential to revolutionize how we analyze tissue microenvironments, accelerating the discovery of novel biomarkers and therapies to ultimately improve the translation of biological discovery to patient outcomes.

## Data Availability

Public Xenium FFPE skin sections were downloaded from 10x Genomics (https://www.10xgenomics.com/datasets/xenium-prime-ffpe-human-skin). Simulated data is available for download as described at https://github.com/Lotfollahi-lab/mintflow-reproducibility. All other data will be made available soon.

## Code Availability

MintFlow is available as a Python package, maintained at https://github.com/Lotfollahi-lab/mintflow. All code to reproduce experiments, including benchmarking and analyses, is available at https://github.com/Lotfollahi-lab/mintflow-reproducibility. We provide comprehensive documentation encompassing tutorials and a user guide at https://mintflow.readthedocs.io/.

## Acknowledgments

We acknowledge core funding from Wellcome (WT220540/Z/20/A). M.H. is funded by Wellcome (WT107931/Z/15/Z) and the CIFAR MacMillan Multiscale Human program. M.L and M.H acknowledge support and funding from Open Targets. S.B. is supported by the Helmholtz Association under the joint research school “Munich School for Data Science—MUDS”. D.J.J. is supported by a Wellcome Trust Accelerator Award (314710/Z/24/Z), the Foulkes Foundation and the Specialised Foundation Programme at the East of England NHS Deanery. L.S. is supported by a Wellcome Clinical Research Training Fellowship and Trinity Internal Graduate Studentship. S.B. is grateful to Paula, Pebble and Pixel Villa Fulton for their inspirational support.

## Author Contributions

M.L. and M.H. conceived the project and designed the experiments. A.A. designed and implemented the model with contributions from S.B., A.M. and theoretical feedback and contributions from S.V. L.S., D.J.J., M.R.S., K.R. and M.R.C. performed data analysis. A.B., B.R., C.T., M. Patel, M. Prete, S. Makarchuk, S. Mahil, T.L., H.S., A.R.F., K. Roberts, A.L.T., and C.S. contributed to sample collection, performing experiments, data generation and processing. C.E.V. and G.T. supported M.R.S. T.M. supported D.J.J., contributed to data generation and provided kidney samples. O.A.B., M.H. and M.L. supervised the work. S.B., A.A., D.J.J., M.R.S., O.A.B., M.H., M.L. and L.S. wrote the paper.

## Methods

### Model overview and assumptions

Our proposed generative model (Supplementary Fig. 17) explicitly models intrinsic as well as spatial signals between a cell and its microenvironment, generating intrinsic and microenvironment-induced embeddings for each cell. A cell’s intrinsic embedding is conditioned on its cell type label, while its microenvironment-induced embedding is conditioned on its microenvironment cell type composition (MCC). To define cells’ microenvironments, a k-Nearest Neighbors graph (k=5) is created where nodes represent cells and edges connect neighboring cells. A cell’s set of neighbors in the graph is considered its microenvironment, and its MCC is computed as the proportion of each cell type within its microenvironment. Cells’ embeddings and their relation are defined as follows. The variable 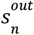, a priori conditioned on cell type, is the signal that the *n*-th cell communicates to neighboring cells in the microenvironment. In the *n*-the cell, the 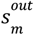 signals from all neighbor cells *m* in the microenvironment are received and aggregated in 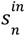 via simple neighborhood averaging: 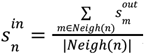,where *Neigh*(*n*) is the set of neighbors of the *n*-th cell, and |*Neigh(n*)| is its node degree. As 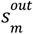 is a priori conditioned on the cell type of cell *m*, the aggregation indirectly conditions 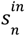 on the microenvironment cell type composition of cell n. Intuitively,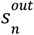 is expected to contain information about the signal that the *n*-th cell secrets into its microenvironment, while 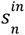 contains information about the signal that the *n*-th cell receives from its microenvironment. The variable *z*_*n*_, apriori conditioned on cell type, captures the intrinsic variation of the *n*-th cell, which is not influenced by its microenvironment. Intuitively, *z*_*n*_ comprises internal signals such as the expression of receptors, cell cycle regulators, and housekeeping genes. We assume the expression in each cell has two components:

1. The part which is not triggered by a signal from outside: In the *n*-th cell this part of expression is encoded in 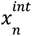. It is only affected by the internal cell state *z*_*n*_ (Supplementary Fig. 17)
2. The part which is triggered by a signal from outside: In the *n*-th cell this part of expression is encoded in 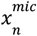. It is affected by the signal that the *n*-th cell receives from its neighbors, encoded in variable 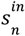, and internal properties that are relevant for processing these external signals such as the receptor expression which is encoded in the variable *z* _*n*_(Supplementary Fig. 17).

Since training flow-based generative models on high-dimensional data is impractical^64^, we perform flow matching in latent space, using a neural ordinary differential equation (neural ODE)^65^ decoder to obtain 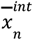 and 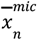, which are lower-dimensional representations of the intrinsic 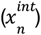 and microenvironment-induced 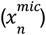 components of expression. We then employ decoders that take in 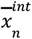 and 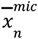 and generate the intrinsic and microenvironment-induced gene expression vectors 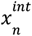 and 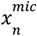, respectively. We denote the count readout in the *n*-th cell by *x*_*n*_. We have that 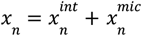, where 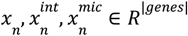 and |*genes*| is the number of genes. The readouts *x*_*n*_ are observed while the vectors 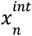 and 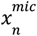 are latent. Note that we could have directly decoded *x*_*n*_ from 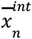 and 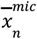, completely bypassing and discarding 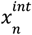 and 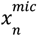. However, the inclusion of 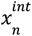 and 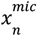 provides an explicit decomposition of the readout matrix as *X* = *X*^*int*^ + *X*^*mic*^, where *X*^*int*^ and *X*^*mic*^ are cell by gene matrices and can directly be used for biological analysis. In other words, this design choice adds interpretability to our model thereby removing the need for archetypal or linear probe analysis on the low-dimensional embeddings, which is used by other methods^10^.

### Notes on identifiability and objective terms

Our proposed model is a deep generative model to generate expression data 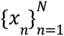. Such a model with non-linear decoders is known to be unidentifiable in general^66^. However, a line of research guarantees identifiability if latent factors are conditioned on some side information^67–70^.

Inspired by this, we condition the latent embeddings *z*_*n*_ and 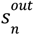 of the *n*-th cell on its cell type annotation, denoted by the one-hot encoded vector *t*_*n*_(Supplementary Fig. 17). In particular, for 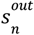 we define a conditional normal distribution whose mean is a “linear” function of *t*_*n*_ as follows:

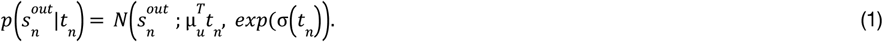

Therefore, for the *n*-th cell according to Eq. (1) the prior on 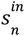 is a normal distribution with mean 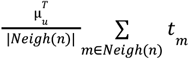. Note that since *t*_*m*_ are one-hot encoded vectors, the term 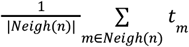 contains the frequency of different cell types among neighbors of the *n*-th cell. We refer to this vector as MCC (microenvironment cell type composition) and to the cell type one-hot encoded vector as CT (cell type).

With this setup, we mimic the iVAE method^67^ that guarantees identifiability by conditioning the latent embedding on some label *u* (Supplementary Fig. 18a). In our case, we condition *z* and *s*^*in*^ on CT and MCC (Supplementary Fig. 18b). The only iVAE^67^ assumption that we violate at this point is that in our case the gene expression vectors *x* could be interleaved and dependent among neighboring cells while in iVAE^67^ the *x* vectors are assumed independent.

Consequently, in our proposed method the vector [*z, s*^*in*^] is expected to be identifiable up to permutation of its dimensions. Notably the *dim*(*z*) + *dim*(*s*) dimensions of the vector [*z, s*^*in*^] can freely permute, hindering the separation of intrinsic and microenvironment-induced embeddings. To identify *z* and *s*^*in*^, we therefore introduce two additional objectives:

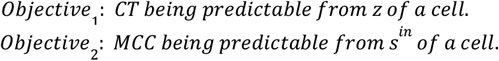

Intuitively, among the *dim*(*z*) + *dim*(*s*) dimensions of the vector [*z, s*^*in*^], *dim*(*z*) (resp. *dim*(*s*)) dimensions that are related to CT (resp. MCC) are assigned to *z* (resp.*s*^*in*^). Importantly, although *Ojbective*_1_ and *Objective*_2_ encourage the identification of *z* and *s*^*in*^, the dimensions of those vectors can still freely and benignly permute. In a cell type homogeneous region the CT and MCC vectors become equal, impairing their ability to identify the intrinsic/microenvironment-induced parts of the vector [*z, s*^*in*^]. However, the existence of non-homogenous regions in other tissue regions make the intrinsic/microenvironment-induced dimensions identifiable.

Let *F*(·,·) be the decoder for the observed gene expression vectors, i.e. *F*(*z, s*^*in*^) = *x*. For the sake of argument, let us also assume that no dimensionality reduction happens, i.e. the Variables 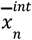 and 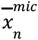 would be omitted (Supplementary Fig. 18c). Although so far we discussed why the vectors *z* and *s*^*in*^ are expected to be identifiable up to permutation, this doesn’t fulfil the objective of the proposed method, because recall that besides identifying the function *F*(·,·), our goal is to decompose the gene expression vector *x* to *x*^*int*^ and *x*^*mic*^. Let’s assume that the function *F*(·,·) is known and we want to decompose it as

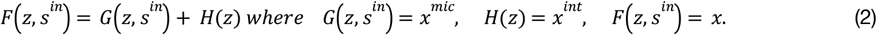

One unsatisfactory solution to the decomposition of Eq. 2 is *H*(*z*) being zero for all *z*, and *F*(·,·) = *G*(·,·) for all *z* and *s*^*in*^. In other words, this unsatisfactory solution effectively ignores the path 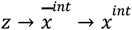 (Supplementary Fig. 17a) and, for all cells, puts all read counts *x* in *x*^*mic*^. Conversely, the decoder *G*(*z, s*^*in*^) may ignore *s*^*in*^ by putting all the dataset variation in *z*, leading to another unsatisfactory solution. Empirically, we found the former bad solution quite likely to be found by the method. To avoid these cases we introduce two additional objectives:

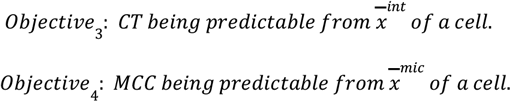

Putting aside unwanted solutions, without any additional assumptions the decomposition of Eq. 2 is not unique. Because for any arbitrary *H*(*z*), *G* (*z, s*^*in*^) could be simply defined as *F*(*z, s*^*in*^) − *H*(*z*) and Eq. 2 holds. To make the decomposition of Eq. 2 unique, we impose conditions related to optimal transport.

#### Proposition 1.

Consider the decomposition of Eq. 2 where *F*(·,·) is known and *G*(·,·) and *H*(·) are to be found. Let λ be an arbitrary positive scalar and *c*(·,·) be the Euclidean distance. If *G*(·,·) and *H*(·) are constrained to optimize the cost functional of Eq. 3, then in the decomposition of

Eq. 2 *G*(·,·) and *H*(·) are unique. Put differently, the optimization functional problem of Eq. 3 - when *H*(·) and *G*(·,·) are constrained to sum up to a known function *F*(·,·) according to Eq. 2 - has a unique solution.

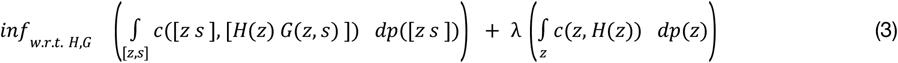

Notice that in Proposition 1 each term in Eq. 3 looks similar to Monge’s formulation of optimal transport but without the boundary condition on the push forward function, and instead with the summation constraint of Eq. 2 on *H*(·) and *G*(·,·). Proof and a more formal statement of Proposition 1 is provided in Supplementary Note 2.

The connection between flow matching with mini-batches and the global optimal transport plan is not completely understood^71^, and our method likewise is not guaranteed to find the optimal solution of Eq. 3. Nevertheless by introducing the flow matching objective we expect the marginal vector field to remain relatively simple^72^ and remain somewhat close to optimal for the objective of Eq. 3. To this end, we add the following objectives:

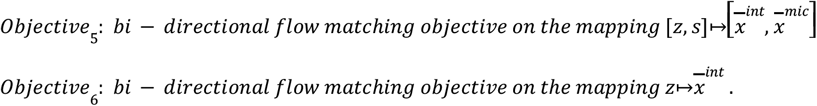

Note that one can consider *Objective*_5_ and *Objective*_6_ as independent and separate objectives in addition to the usual evidence lower-bound (ELBO) terms, but we design the variational family such that they appear in the ELBO (Supplementary Fig. 19 and Supplementary Note 5). The above discussion sums up the rationale behind the objective terms and describes why the proposed model is expected to be identifiable. However, to further encourage separation of intrinsic and spatial signals, we consider two additional objectives:

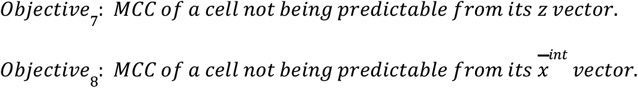

The above objectives are induced by the discretized version of MCC, meaning from *z* or 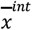 of a cell a predictor cannot tell if it resides next to any specific cell type. This is in line with the basic intuition that the internal state of a cell does not contain information about the type of neighboring cells. *Objective*_7_ and *Objective*_8_ are implemented by Wasserstein distance distribution matching^73,74^. More precisely, for each cell type population, there is a cell type-specific discriminator (or dual function in the sense of Wasserstein distance estimation) that takes in *z* or 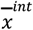 of a cell in the population and predicts if it resides in the neighborhood of any cell of that cell type assigned to the cell type-specific discriminator. In other words, if there are |*cell type*|-many cell types, we consider |*cell type*| × |*cell type*|-many discriminators or dual functions where the fist |*cell type*| multiplicand is pertinent to the number of populations per cell type and the second |*cell type*| multiplicand corresponds to each MCC vector dimension. To reduce the amount of computation, we consider |*cell type*| MLP backbones each having |*cell type*|-many output heads. We implemented Wasserstein distribution matching similar to Gulrajani *et al*^*74*^.

Finally, as a standard modeling assumption^75^, we encourage the intrinsic embeddings and expression components of cells with the same cell type label to be close. This is done via a mean squared loss for each pair of cells in the subgraph returned by PyTorch Geometric’s neighbor loader and for their intrinsic embeddings and expression components:

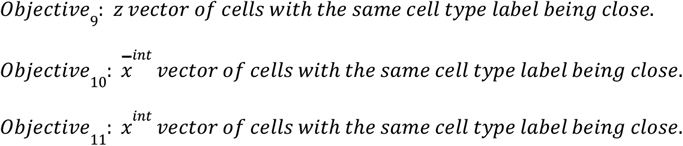

### Scalability

We now describe how our proposed MintFlow method achieves scalability (the details of the encoder/decoder formulation and architectures are provided in Supplementary Notes 3 and 4). The main hurdle towards scalability is that representations are interleaved via the spatial neighborhood graph; In the generative model (Supplementary Fig. 17a) it happens because the *s*^*our*^ vectors are averaged among neighbours, and on the inference side (Supplementary Fig. 17b) it happens because a graph neural network (GNN) decomposes the gene expression vector *x* ^*n*^ of the *n*-th cell given those of its neighbors, to arrive at 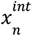 and 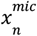. To solve this issue we adopted inductive graph learning which enables learning GNNs with subsampled subgraphs rather than processing the entire graph as a whole^76^. Given a GNN with *k*-hop receptive field, a selected subgraph has two sets of nodes: (1) central nodes for whom all nodes in the *k*-hop neighborhood (testing phase) or a subset of neighbors in 1-, 2-, 3-, …, and *k*-hop are present in the subgraph. (2) non-central nodes whose role is to provide to the GNN the context around the central nodes, so it can compute reliable values for them. In other words, GNN’s output for the central nodes is valid and, e.g., a graph node classification loss can be computed from them. But the GNN’s output for non-central nodes is not necessarily valid. This poses a challenge for MintFlow, because the vectors *x*^*int*^ and *x*^*mic*^ need to be computed for all cells in the subgraph (Supplementary Fig. 20a), including the central and non-central ones, from which the other vectors *z,s* ^*in*^, and *s*^*out*^ are computed for all cells. Doing so is necessary, because in the decoding phase all *s*^*out*^ vectors are needed, which in turn necessitates having *x*^*int*^ and *x*^*mic*^ for all cells. To alleviate this problem we resort to the probabilistic nature of the model; in the variational distribution we put a lower-bound on the predicted variance (i.e. uncertainty) for the *x*^*int*^ and *x*^*mic*^ vectors of non-central nodes to emphasize that those vectors are only some rough estimates by the encoder. Furthermore, we use the customized sampler described next.

We used PyTorch Geometric’s neighbor loader^76^ both for training and testing. The default sampler randomly selects some nodes (i.e. cells) as the central nodes and adds the *k*-hop neighbors to the subgraph. By doing so, non-central nodes (i.e. cells) may dominate the subgraph (Supplementary Fig. 20a). To tackle this issue we use a customized sampler that selects random crops from a tissue section (the blue squares in Supplementary Fig. 20b) as being the central nodes. Consequently, central nodes’ neighbors are likely to be among central nodes, and non-central nodes are expected to reside on the boundaries of the selected crops (the red regions in Supplementary Fig. 20b). Thus, central nodes (i.e. cells) are expected to dominate the selected subgraph. Recall that - unlike the regular case of, e.g., training a GNN node classifier - we need to use encoder predictions for both central and non-central nodes, hence in our use case the customized sampler is highly preferable. For efficient and scalable evaluation, we used the layered approach^77^ in the testing phase.

### Handling batch effect

MintFlow can be applied to a set of tissue sections with known or unknown technical or biological batch effects between them. To handle batch effects, the decoder modules for 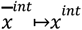 and 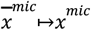 as well as the first encoder module for *x*↦[*x*^*int*^, *x*^*mic*^] take in a one-hot encoded batch identifier. Furthermore, we implemented Wasserstein distribution matching similar to Gulrajani *et al*.^74^ via two batch discriminators (or Wasserstein distance estimators) placed on top of 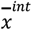 and 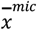. Intuitively, in the model of Supplementary Fig. 17a, the variables 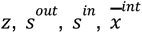 and 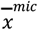 are encouraged to be devoid of batch effects while *x*^*int*^ and *x*^*mic*^ are batch- and tissue section-dependant. For this reason, when running MintFlow on more than one tissue section with multiple batch identifiers, we use 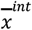 and 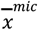 for different downstream analyses like cell subpopulation identification. However, when MintFlow is run on a single tissue section one can use *x*^*int*^ and *x*^*mic*^, e.g., after dimensionality reduction with PCA. There are two independently-tunable coefficients for batch mixing loss terms of 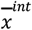 and 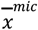 whereby one can unequally encourage the batch mixing objectives on these embeddings. For example, one may want to encourage batch mixing mostly on 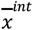 rather than 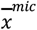, and this can be done by setting the aforementioned two loss term coefficients to proper values.

### Annealing the reconstruction loss

In our proposed model there are two latent vectors *x*^*int*^ and *x*^*mic*^ with as many dimensions as the number of genes (Supplementary Fig. 17a), and this causes complications, because two encoder/decoder architectures are needed: one for 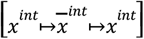 and one for 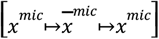. In particular, the predicted *x*^*int*^ and *x*^*mic*^ vectors keep changing during training and the corresponding encoder/decoder have to adapt their parameters on the fly. Conversely, during training the encoder/decoder reconstruction losses discourage the variational distribution to freely modify the assignment of read counts to *x*^*int*^ and *x*^*mic*^, because doing so increases the reconstruction loss of the encoder/decoder. To alleviate this issue, we anneal the reconstruction loss, starting from a small loss coefficient and increasing it linearly to a larger number as the training continues, and finally continuing the training for a number of epochs with the largest coefficient for reconstruction loss.

### Computing the microenvironment score

We use the identified *x*^*int*^ and *x*^*mic*^ vectors to compute a per-cell microenvironment score, which for the *n*-th cell is computed as: 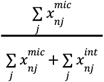, where the *j* index enumerates genes.

Intuitively, a higher microenvironment score means that a larger proportion of the cell’s total count is predicted to be induced by the microenvironment. Therefore, as shown in Fig. 1d, colocalized cells with high microenvironment scores are candidate signaling hotspots.

### *In silico* tissue perturbation

After training, the generative model can generate expression data merely given cell type labels and the spatial neighborhood graph of cells (Supplementary Fig. 17a). Notably, the generative model generates microenvironment-induced and intrinsic components of expression as well as total expression generated as the sum of the two components. This enables *in silico* perturbation of a tissue region, generating expression data for the pre- and post-perturbed tissue region, and finally observing how the generated expression vectors and their microenvironment-induced and intrinsic components change between pre- and post-perturbed tissue. Our *in silico* perturbation allows to arbitrarily change cell type labels and the spatial neighborhood graph. Nonetheless, in our experiments we picked a tissue crop from real data and either altered cell type labels or discarded subpopulations of cells of specific types. Afterwards, the neighborhood graph is recreated for the post-perturbed tissue based on spatial locations of cells.

### Datasets

We used MintFlow to analyze spatial transcriptomics data from four datasets: a simulated dataset, newly generated atopic dermatitis, and ccRCC datasets and publicly available data of cutaneous melanoma (Supplementary Note 6 for detailed description). All spatial data were profiled using the 10x Genomics Xenium in situ 5k-plex platform and, where specified, analyzed with matched single-cell or single-nucleus RNA-seq datasets for validation and annotation. All human samples were obtained with appropriate ethics approvals and informed consent.

### Data generation and acquisition

Cutaneous melanoma data was previously published (Data Availability section). All other data was newly generated.

#### Simulated data

Simulated spatial data were generated using the NicheCompass simulation framework^7^, and comprised 10,000 cells and 2,000 genes across eight spatial regions. In the experiments presented in Supplementary Fig. 3, we excluded regions with MCCs consisting of only one cell type, because this is a harder setting where the MCC and CT labels are the same in some tissue regions and thus the separation of intrinsic and microenvironment-induced effects is challenging (Methods, *Objective*_1_ and *Objective*_2_).

#### Atopic dermatitis

Five lesional and five non-lesional skin biopsies were obtained from adult patients with atopic dermatitis (REC EC00/128). Sections (10 µm) were prepared and profiled via Xenium-5k, with protein staining for segmentation and subsequent H&E imaging. For investigating perturbation predictions, we collected week 12 samples from the same cohort post-treatment with dupilumab (anti-IL4R monoclonal antibody). Treatment response was based on Eczema Area and Severity Index (EASI) scores, which is a clinically validated tool. Clinical responses were binned as non-responder (<10% improvement) and responder (>50% improvement) - with no intermediate treatment response values in the cohort.

#### Clear cell renal carcinoma

One nephrectomy case (PD43284; REC 03/018) was profiled across four regions (three tumor cores, one tumor–normal interface) and used for MintFlow training. Two other nephrectomy cases (PD45814 and PD45816) were used for validation.

Tissue was processed via the Xenium pipeline as above.

### Preprocessing and cell type annotation

Raw counts were filtered to exclude poor quality cells. Gene expression matrices were normalized before log transformation. Dimensionality reduction using PCA and kNN graph construction, and clustering using the Leiden algorithm, were performed for each dataset. For annotation, we used a dataset-specific approach. For atopic dermatitis and ccRCC, we used scVI embeddings and Leiden clustering for annotation, guided by DEGs, spatial context, and validation via CellTypist and ScanVI^31,24^. Melanoma annotations followed 10x metadata and DEGs for each cell type.

### MintFlow training and outputs

MintFlow was trained on each dataset using the parameters specified in Supplementary Table 1. Utilizing *X*^*mic*^, datasets were clustered to resolve microenvironment-induced gene programs (MGPs), with marker gene identification using scanpy.tl.score_genes. These MGPs were projected onto external scRNA-seq datasets, including a cross-tissue human atlas^24^ and a ccRCC atlas^50^.

### Cell-cell communication, drug repurposing analysis and metabolic profiling

For atopic dermatitis, CellPhoneDB v5^78^ was used to analyze chemokine interactions in the T_DC domain. We used our atopic dermatitis scRNA-seq data to obtain higher resolution cell annotations for immune cells^31^. Drug2cell^26^ was used to rank candidate therapies targeting MGPs. Gene set enrichment scores for GO biological processes were calculated using GSEApy. For scoring the *Stroma* melanoma gene program, we used scanpy.tl.score_genes applied to an integrated dataset of skin fibroblasts^24^. For ccRCC, cell-cell communication was imputed using LIANA+^79^ in “rank aggregate” mode, incorporating multiple ligand–receptor prediction algorithms, including CellPhoneDB, NATMI and CellChat.

### Assessing the biological validity of in silico perturbation

To assess the biological validity of *in silico* TLS perturbations in ccRCC, we leveraged publicly available scRNA-seq data of ccRCC, including samples treated with ICB^58^. This study provided gene expression signatures for T cells and macrophages exposed to ICB therapy. We therefore identified T cells within our ccRCC Xenium data by identifying the top 5% of T cells enriched for the signature genes derived from the scRNA-seq data^58^. We then leveraged MintFlow’s perturbation module to replace all T cells within the TLS region with post-ICB-like T cells, and calculated the predicted effect upon gene expression of neighboring macrophages. To assess the validity of the perturbation, we used genes from the original scRNA-seq study known to be upregulated by ccRCC macrophages after ICB therapy^58^, and calculated a module score and examined the individual changes in gene expression before and after perturbation with dot plots and violin plots. Gene set enrichment scores for GO biological processes enriched in perturbed macrophages were calculated using GSEA.

### Survival analysis

For ccRCC, MintFlow-derived T cell MGPs were applied to TCGA bulk RNA-seq data (*n* = 606 patients), Kaplan–Meier survival curves were generated after stratifying patients by median SGP score using R packages survival and survminer.

### Benchmarking

To benchmark SIMVI^10^, we used the code from https://github.com/KlugerLab/SIMVI (retrieved on April 8th, 2025). We set the hyperparameters as recommended in the MERFISH MTG data tutorial and used embedding size 10. We used the implementation of MEFISTO^13^ in MOFA python package^80^ and with default parameters. As done in Dong *et al*.^*10*^ we computed the median of scale factors computed by MEFISTO^13^, picked the factors whose corresponding scale is higher than the median value, and considered the decoded cell by gene matrix as microenvironment-induced expression vector. The default embedding size of the proposed MintFlow is 100. Whenever comparing MintFlow to other baselines, we reduced the embedding size to 10 since some baselines have huge memory and computation requirements, and are not runnable with embedding size equal to 100. Nonetheless, for some datasets running baselines was not feasible even when their hyperparameters were carefully tuned. In those cases we ran the proposed MintFlow with default hyperparameters. For MintFlow, we used slightly different settings in different analyses. For example, in the benchmarking of Fig. 1h (left and middle), we reduced the size of embeddings to 10, and in the ablation study of Supplementary Fig. 21 we reduced the number of epochs, thereby making it possible to run several experiments in a reasonable time. Our experiments’ configuration files for MintFlow CLI (command-line interface) are available online: https://github.com/Lotfollahi-lab/mintflow-reproducibility.

### Metrics

For simulated data, the ground truth intrinsic and microenvironment-induced expression matrices (i.e. ground truth *x*^*int*^ and *x*^*mic*^) are available. Therefore we used common metrics to measure how close MintFlow’s predicted *x*^*mic*^ was to the ground truth: MAE (mean absolute error), MSE (mean squared error), and EMD (earth mover’s distance). We computed the MAE and MSE using numpy, and EMD using scipy.stats.wasserstein_distance.

For real spatial transcriptomics data, no ground truth for the microenvironment-induced component of expression is available, thus we cannot use the MAE, MSE, and EMD metrics. Therefore we acquired a list of genes expected to be related to cell-cell signaling. Specifically, we processed and combined many databases gathered at https://github.com/LewisLabUCSD/Ligand-Receptor-Pairs^19^. In our list we included each gene of interacting complexes, whenever complexes were reported in each database. We consider any gene that appears in any database as a “signaling gene” and otherwise a “non-signaling gene”. Intuitively, if a read count (*i*.*e*. one element of the observed cell by gene matrix) corresponds to a signaling gene, then MintFlow is expected to assign a higher proportion of it to *x*^*mic*^ relative to *x*^*int*^. This criterion is used in Fig. 1h.

### Ablation study on simulated data

To experimentally validate Proposition 1 and show the effect of the flow matching objective we conducted experiments on simulated data. We varied:

- The importance coefficient of the flow matching objectives (*i*.*e*.*Objective* _5_ and *Objective*_6_).
- The scale factor for the encoders of *z* and *s*^*out*^. Note that when the scale becomes large, in the limit it resembles how bi-directional flow matching^71^ would generate samples of [*z, s*^*in*^] from the based distribution.

The former and latter were varied between {0· 01, 0· 1, 0· 2, 0· 5, 1· 0} and {0· 01, 0· 1, 0· 2, 0· 5, 1· 0}, respectively, and the metrics MSE, MAE, and EMD were computed for 3 random runs in each experiment (Supplementary Fig. 21). The first row shows the average of each metric over 3 different runs. The second row shows the box plots for each anti-diagonal of the heatmaps. Note that, unlike in the heatmaps, all 3 runs are included in the box plots. For example “anti-diag 1” corresponds to row 1 and column 1 in the heatmaps (*i*.*e*. one element of the matrix), but since there are 3 runs the box of “anti-diag 1” in the box plot contains 3 circles. In Supplementary Fig. 21 the third and fourth rows show the boxplots for each row and each column. According to rows 1 and 2, as both the encoder noise for [*z, s*^*in*^] and the coefficient of flow matching loss increase (*i*.*e*. the lower right section of the heatmaps), the MSE and MAE metrics (cols 1 and 2) and Wasserstein metrics for *z* and [*z, s*^*in*^] (cols 3 and 4) reduce. Recall that the Wasserstein metrics correspond to the terms in Eq. (3) in Proposition 1, hence these figures show that reducing those terms helps in identifying the latent intrinsic and microenvironment-induced expression components, as anticipated by Proposition 1.

### Incremental application of atopic dermatitis in silico perturbations

We repeated the atopic dermatitis perturbations (Fig. 3b,e) but instead of perturbing all cells of interest at once, we incrementally deleted or changed cells of specific types and inspected the gene expression of cells whose microenvironment cell type composition (MCC) had changed from the initial unperturbed tissue. For the *in silico* perturbation of Treg depletion we recreated the neighborhood graph after each incremental deletion of Treg cells. For reporting top DEGs, we selected genes from the top 2000 genes for the cell type from a skin single cell atlas ^31^.

## References

1. Rood, J. E., Maartens, A., Hupalowska, A., Teichmann, S. A. & Regev, A. Impact of the Human Cell Atlas on medicine. Nat. Med. 28, 2486–2496 (2022).

2. Binnewies, M. et al. Understanding the tumor immune microenvironment (TIME) for effective therapy. Nature Medicine 24, 541–550 (2018).

3. Ji, A. L. et al. Multimodal Analysis of Composition and Spatial Architecture in Human Squamous Cell Carcinoma. Cell 182, 497–514.e22 (2020).

4. Fiskin, E. et al Multi-modal skin atlas identifies a multicellular immune-stromal community associated with altered cornification and specific T cell expansion in atopic dermatitis. bioRxiv (2023) doi:10.1101/2023.10.29.563503.

5. Habib, N. et al. Disease-associated astrocytes in Alzheimer’s disease and aging. Nat Neurosci 23, 701–706 (2020).

6. Fischer, D. S., Schaar, A. C. & Theis, F. J. Modeling intercellular communication in tissues using spatial graphs of cells. Nat. Biotechnol. 41, 332–336 (2023).

7. Birk, S. et al. Quantitative characterization of cell niches in spatially resolved omics data. Nat. Genet. (2025) doi:10.1038/s41588-025-02120-6.

8. Singhal, V. et al. BANKSY unifies cell typing and tissue domain segmentation for scalable spatial omics data analysis. Nat. Genet. (2024) doi:10.1038/s41588-024-01664-3.

9. Varrone, M., Tavernari, D., Santamaria-Martínez, A., Walsh, L. A. & Ciriello, G. CellCharter reveals spatial cell niches associated with tissue remodeling and cell plasticity. Nat. Genet. 56, 74–84 (2024).

10. Dong, M., Su, D. G., Kluger, H., Fan, R. & Kluger, Y. SIMVI disentangles intrinsic and spatial-induced cellular states in spatial omics data. Nat. Commun. 16, 2990 (2025).

11. Levy, N., Ingelfinger, F., Ergen-Behr, C. & Nadler, B. Nichevi: A probabilistic framework to embed cellular interaction in spatial transcriptomics. (2024).

12. Jerby-Arnon, L. & Regev, A. DIALOGUE maps multicellular programs in tissue from single-cell or spatial transcriptomics data. Nature Biotechnology 40, 1467–1477 (2022).

13. Velten, B. et al. Identifying temporal and spatial patterns of variation from multimodal data using MEFISTO. Nat. Methods 19, 179–186 (2022).

14. Townes, F. W. & Engelhardt, B. E. Nonnegative spatial factorization applied to spatial genomics. Nat. Methods 20, 229–238 (2023).

15. Chen, S., Regev, A., Condon, A. & Ding, J. CellUntangler: separating distinct biological signals in single-cell data with deep generative models. bioRxiv (2025) doi:10.1101/2025.01.10.632490.

16. Efremova, M., Vento-Tormo, M., Teichmann, S. A. & Vento-Tormo, R. CellPhoneDB: inferring cell-cell communication from combined expression of multi-subunit ligand-receptor complexes. Nat. Protoc. 15, 1484–1506 (2020).

17. Jin, S. et al. Inference and analysis of cell-cell communication using CellChat. Nat. Commun. 12, 1088 (2021).

18. Wolf, F. A., Angerer, P. & Theis, F. J. SCANPY: large-scale single-cell gene expression data analysis. Genome Biol. 19, 15 (2018).

19. Armingol, E., Officer, A., Harismendy, O. & Lewis, N. E. Deciphering cell-cell interactions and communication from gene expression. Nat. Rev. Genet. 22, 71–88 (2021).

20. Morrison, V. L. et al. The β2 integrin-kindlin-3 interaction is essential for T-cell homing but dispensable for T-cell activation in vivo. Blood 122, 1428–1436 (2013).

21. Montoya, M. C. et al. Role of ICAM-3 in the initial interaction of T lymphocytes and APCs. Nat. Immunol. 3, 159–168 (2002).

22. Gérard, A., Cope, A. P., Kemper, C., Alon, R. & Köchl, R. LFA-1 in T cell priming, differentiation, and effector functions. Trends Immunol. 42, 706–722 (2021).

23. Jin, H. et al. IL-21R is essential for epicutaneous sensitization and allergic skin inflammation in humans and mice. J. Clin. Invest. 119, 47–60 (2009).

24. Steele, L. et al. A single cell and spatial genomics atlas of human skin fibroblasts in health and disease. bioRxiv 2024.12.23.629194 (2024) doi:10.1101/2024.12.23.629194.

25. Liu, J., Zhang, X., Cheng, Y. & Cao, X. Dendritic cell migration in inflammation and immunity. Cell. Mol. Immunol. 18, 2461–2471 (2021).

26. Kanemaru, K. et al. Spatially resolved multiomics of human cardiac niches. Nature 619, 801–810 (2023).

27. Bachert, C. et al. Biologics for chronic rhinosinusitis with nasal polyps. J. Allergy Clin. Immunol. 145, 725–739 (2020).

28. Han, Y. et al. Amlexanox exerts anti-inflammatory actions by targeting phosphodiesterase 4B in lipopolysaccharide-activated macrophages. Biochim. Biophys. Acta Mol. Cell Res. 1867, 118766 (2020).

29. Weidinger, S., Beck, L. A., Bieber, T., Kabashima, K. & Irvine, A. D. Atopic dermatitis. Nat. Rev. Dis. Primers 4, 1 (2018).

30. Chen, X., Lin, J., Liang, Q., Chen, X. & Wu, Z. Pseudoephedrine alleviates atopic dermatitis-like inflammatory responses in vivo and in vitro. Life Sci. 258, 118139 (2020).

31. Reynolds, G. et al. Developmental cell programs are co-opted in inflammatory skin disease. Science 371, eaba6500 (2021).

32. Kato, A. & Kita, H. The immunology of asthma and chronic rhinosinusitis. Nat. Rev. Immunol. 1–19 (2025).

33. Pitzalis, C., Jones, G. W., Bombardieri, M. & Jones, S. A. Ectopic lymphoid-like structures in infection, cancer and autoimmunity. Nat. Rev. Immunol. 14, 447–462 (2014).

34. Migayron, L., Merhi, R., Seneschal, J. & Boniface, K. Resident memory T cells in nonlesional skin and healed lesions of patients with chronic inflammatory diseases: Appearances can be deceptive. J. Allergy Clin. Immunol. 153, 606–614 (2024).

35. Obers, A. et al. Retinoic acid and TGF-β orchestrate organ-specific programs of tissue residency. Immunity 57, 2615–2633.e10 (2024).

36. Reina-Campos, M. et al. Tissue-resident memory CD8 T cell diversity is spatiotemporally imprinted. Nature 1–10 (2025).

37. Osaki, M. & Sakaguchi, S. Soluble CTLA-4 regulates immune homeostasis and promotes resolution of inflammation by suppressing type 1 but allowing type 2 immunity. Immunity 58, 889–908.e13 (2025).

38. Bluestone, J. A. et al. Regulatory T cell therapies to treat autoimmune diseases and transplant rejection. Nat. Immunol. 26, 819–824 (2025).

39. Ho, P. et al. Harnessing regulatory T cells to establish immune tolerance. Sci. Transl. Med. 16, eadm8859 (2024).

40. Moon, C. Y. et al. Dendritic cell maturation in cancer. Nat. Rev. Cancer 25, 225–248 (2025).

41. Wang, W. et al. Single-cell profiling identifies mechanisms of inflammatory heterogeneity in chronic rhinosinusitis. Nat. Immunol. 23, 1484–1494 (2022).

42. Arpinati, L., Carradori, G. & Scherz-Shouval, R. CAF-induced physical constraints controlling T cell state and localization in solid tumours. Nat. Rev. Cancer 24, 676–693 (2024).

43. Preview Data: FFPE Human Skin Primary Dermal Melanoma with 5K Human Pan Tissue and Pathways Panel. 10x Genomics https://www.10xgenomics.com/datasets/xenium-prime-ffpe-human-skin.

44. Gao, Y. et al. Cross-tissue human fibroblast atlas reveals myofibroblast subtypes with distinct roles in immune modulation. Cancer Cell 0, (2024).

45. Sangaletti, S. et al. Mesenchymal transition of high-grade breast carcinomas depends on extracellular matrix control of myeloid suppressor cell activity. Cell Rep. 17, 233–248 (2016).

46. Murdamoothoo, D. et al. Tenascin-C immobilizes infiltrating T lymphocytes through CXCL12 promoting breast cancer progression. EMBO Mol. Med. 13, e13270 (2021).

47. Yamauchi, M., Barker, T. H., Gibbons, D. L. & Kurie, J. M. The fibrotic tumor stroma. J. Clin. Invest. 128, 16–25 (2018).

48. Chhabra, Y. & Weeraratna, A. T. Fibroblasts in cancer: Unity in heterogeneity. Cell 186, 1580–1609 (2023).

49. Lichtenberger, B. M., Mastrogiannaki, M. & Watt, F. M. Epidermal β-catenin activation remodels the dermis via paracrine signalling to distinct fibroblast lineages. Nat. Commun. 7, 10537 (2016).

50. Li, R. et al. Mapping single-cell transcriptomes in the intra-tumoral and associated territories of kidney cancer. Cancer Cell 40, 1583–1599.e10 (2022).

51. Masuda, T. et al. Unique characteristics of tertiary lymphoid structures in kidney clear cell carcinoma: prognostic outcome and comparison with bladder cancer. J. Immunother. Cancer 10, e003883 (2022).

52. Xu, W. et al. Heterogeneity in tertiary lymphoid structures predicts distinct prognosis and immune microenvironment characterizations of clear cell renal cell carcinoma. J. Immunother. Cancer 11, e006667 (2023).

53. Meylan, M. et al. Tertiary lymphoid structures generate and propagate anti-tumor antibody-producing plasma cells in renal cell cancer. Immunity 55, 527–541.e5 (2022).

54. Teillaud, J.-L., Houel, A., Panouillot, M., Riffard, C. & Dieu-Nosjean, M.-C. Tertiary lymphoid structures in anticancer immunity. Nat. Rev. Cancer 24, 629–646 (2024).

55. Pichler, R. et al. A chemokine network of T cell exhaustion and metabolic reprogramming in renal cell carcinoma. Front. Immunol. 14, 1095195 (2023).

56. Braun, D. A. Progressive immune dysfunction with advancing disease stage in renal cell carcinoma Cancer Cell. 39, 632–648.

57. Lee, C. Y. et al. In vivo labelling resolves distinct temporal, spatial, and functional properties of tumour macrophages, and identifies subset-specific effects of PD-L1 blockade. Cancer Immunol. Res. (2025) doi:10.1158/2326-6066.CIR-24-1233.

58. Bi, K. et al. Tumor and immune reprogramming during immunotherapy in advanced renal cell carcinoma. Cancer Cell 39, 649–661.e5 (2021).

59. Cheuk, S. et al. CD49a expression defines tissue-resident CD8+ T cells poised for cytotoxic function in human skin. Immunity 46, 287–300 (2017).

60. Matos, T. R. et al. Clinically resolved psoriatic lesions contain psoriasis-specific IL-17-producing αβ T cell clones. J. Clin. Invest. 127, 4031–4041 (2017).

61. Jingushi, K. et al. Leukocyte-associated immunoglobulin-like receptor 1 promotes tumorigenesis in RCC. Oncol. Rep. 41, 1293–1303 (2019).

62. Kato, R. et al. TIM3 expression on tumor cells predicts response to anti-PD-1 therapy for renal cancer. Transl. Oncol. 14, 100918 (2021).

63. ClinicalTrials.gov. https://clinicaltrials.gov/study/NCT06546553?cond=Kidney%20Cancer&term=LILRB1&rank=1.

64. Rombach, R., Blattmann, A., Lorenz, D., Esser, P. & Ommer, B. High-resolution image synthesis with latent diffusion models. arXiv [cs.CV] (2021).

65. Chen, R. T. Q., Rubanova, Y., Bettencourt, J. & Duvenaud, D. Neural ordinary differential equations. arXiv [cs.LG] (2018).

66. Hyvärinen, A. & Pajunen, P. Nonlinear independent component analysis: Existence and uniqueness results. Neural Netw. 12, 429–439 (1999).

67. Khemakhem, I., Kingma, D., Monti, R. & Hyvarinen, A. Variational Autoencoders and Nonlinear ICA: A Unifying Framework. in Proceedings of the Twenty Third International Conference on Artificial Intelligence and Statistics (eds. Chiappa, S. & Calandra, R.) vol. 108 2207–2217 (PMLR, 26--28 Aug 2020).

68. Lachapelle, S. et al. Disentanglement via Mechanism Sparsity Regularization: A New Principle for Nonlinear ICA. in Proceedings of the First Conference on Causal Learning and Reasoning (eds. Schölkopf, B., Uhler, C. & Zhang, K.) vol. 177 428–484 (PMLR, 11--13 Apr 2022).

69. Moran, G. E., Sridhar, D., Wang, Y. & Blei, D. M. Identifiable deep generative models via sparse decoding. arXiv [stat.ML] (2021).

70. Lopez, R. et al. Learning Causal Representations of Single Cells via Sparse Mechanism Shift Modeling. in Proceedings of the Second Conference on Causal Learning and Reasoning (eds. van der Schaar, M., Zhang, C. & Janzing, D.) vol. 213 662–691 (PMLR, 11--14 Apr 2023).

71. Tong, A. et al. Improving and generalizing flow-based generative models with minibatch optimal transport. arXiv [cs.LG] (2023).

72. Lipman, Y., Chen, R. T. Q., Ben-Hamu, H., Nickel, M. & Le, M. Flow Matching for generative modeling. Int Conf Learn Represent abs/2210.02747, (2022).

73. Arjovsky, M., Chintala, S. & Bottou, L. Wasserstein Generative Adversarial Networks. ICML 70, 214–223 (2017).

74. Gulrajani, I., Ahmed, F., Arjovsky, M., Dumoulin, V. & Courville, A. C. Improved Training of Wasserstein GANs. Neural Inf Process Syst 30, 5767–5777 (2017).

75. Kleshchevnikov, V. et al. Cell2location maps fine-grained cell types in spatial transcriptomics. Nat. Biotechnol. 40, 661–671 (2022).

76. Hamilton, W. L., Ying, R. & Leskovec, J. Inductive representation learning on large graphs. arXiv [cs.SI] (2017).

77. Yin, P. et al. DGI: An easy and efficient framework for GNN model evaluation. in Proceedings of the 29th ACM SIGKDD Conference on Knowledge Discovery and Data Mining 5439–5450 (ACM, New York, NY, USA, 2023).

78. Vento-Tormo, R. et al. Single-cell reconstruction of the early maternal-fetal interface in humans. Nature 563, 347–353 (2018).

79. Dimitrov, D. et al. LIANA+ provides an all-in-one framework for cell-cell communication inference. Nat. Cell Biol. 26, 1613–1622 (2024).

80. Argelaguet, R. et al. MOFA+: a statistical framework for comprehensive integration of multi-modal single-cell data. Genome Biol. 21, 111 (2020).

