## Supplementary Information for "Mapping and reprogramming human tissue microenvironments with MintFlow"

### Supplementary Figures

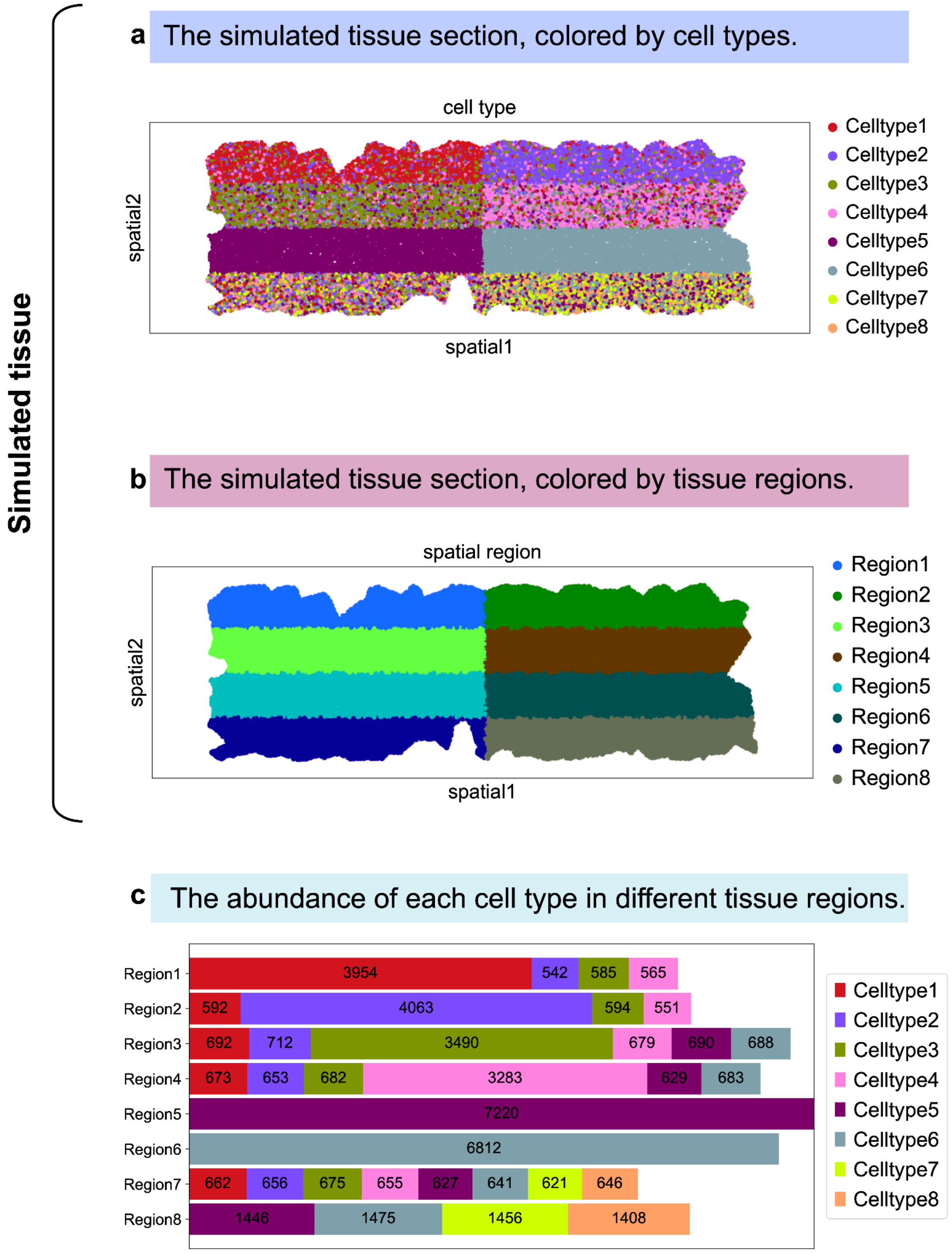

**Supplementary Fig. 1 | Simulated data. a**, The simulated tissue section colored by cell types. **b**, The simulated tissue section colored by regions. There are eight distinct regions and each cell belongs to one of them. **c**, The abundance of each cell type in each tissue region.

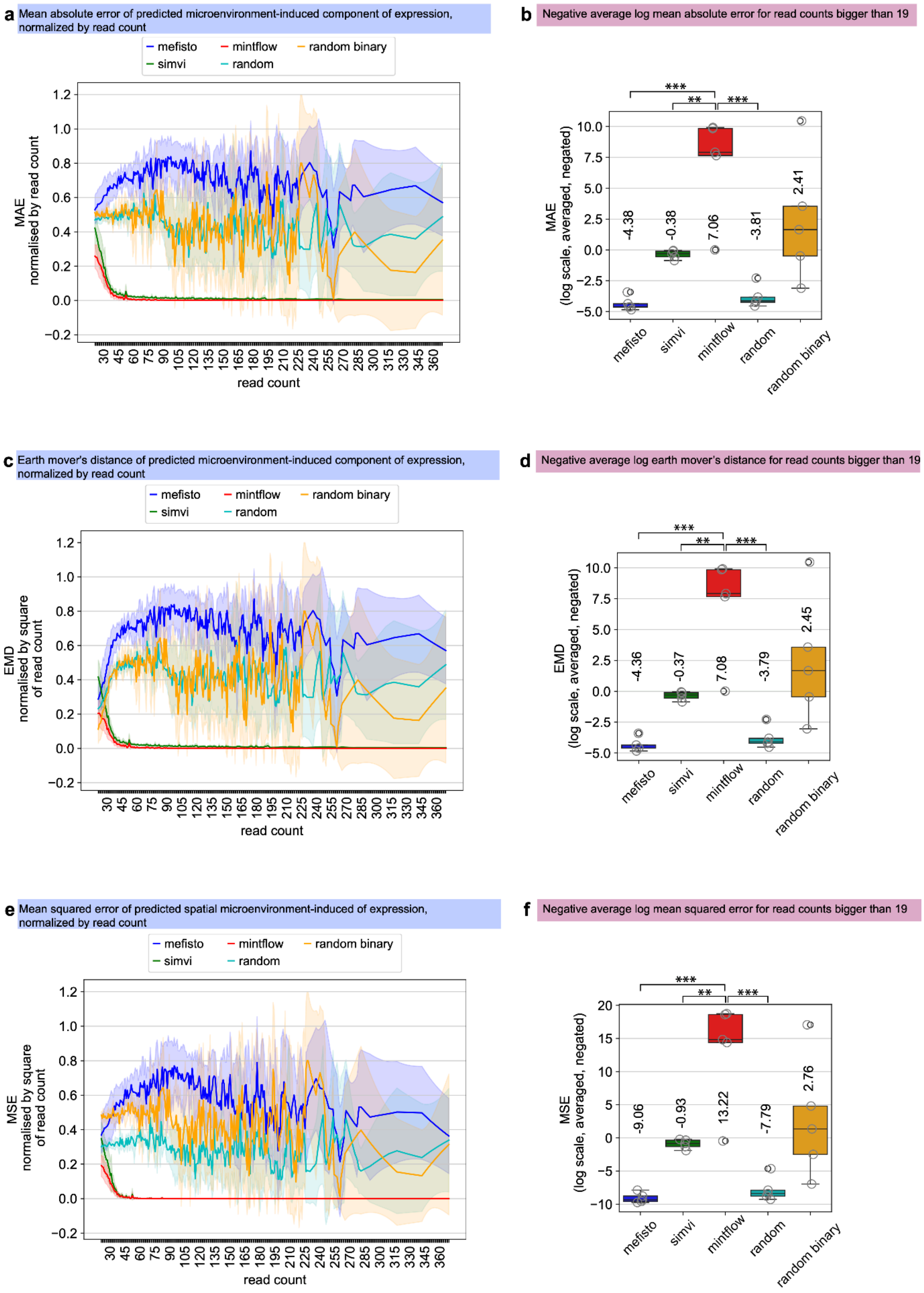
a
Mean absolute error of predicted microenvironment-induced component of expression, normalized by read count
mefisto
mintflow
random binary
simvi
random
MAE
normalised by read count
read count
b
Negative average log mean absolute error for read counts bigger than 19
MAE
(log scale, averaged, negated)
***
**
***
-4.38
-0.38
7.06
-3.81
2.41
mefisto
simvi
mintflow
random
random binary
c
Earth mover's distance of predicted microenvironment-induced component of expression, normalized by read count
mefisto
mintflow
random binary
simvi
random
EMD
normalised by square
of read count
read count
d
Negative average log earth mover's distance for read counts bigger than 19
EMD
(log scale, averaged, negated)
***
**
***
-4.36
-0.37
7.08
-3.79
2.45
mefisto
simvi
mintflow
random
random binary
e
Mean squared error of predicted spatial microenvironment-induced of expression, normalized by read count
mefisto
mintflow
random binary
simvi
random
MSE
normalised by square
of read count
read count
f
Negative average log mean squared error for read counts bigger than 19
MSE
(log scale, averaged, negated)
***
**
***
-9.06
-0.93
13.22
-7.79
2.76
mefisto
simvi
mintflow
random
random binary

**Supplementary Fig. 2 | Method evaluation on simulated data.** Evaluation of different methods in predicting the microenvironment-induced component of gene expression. **a**, **c**, **e**, Mean absolute error (MAE), earth mover's distance (EMD), and mean squared error (MSE) for different read count values. **b**, **d**, **f**, The corresponding metrics for read counts > 19, log-transformed, averaged, and negated. Note that since for simulated data the ground truth intrinsic and microenvironment-induced components of expression are available, we were able to compute these metrics. Boxplot elements are defined as center line, median; box limits, upper and lower quartiles; whiskers, 1.5× interquartile range.

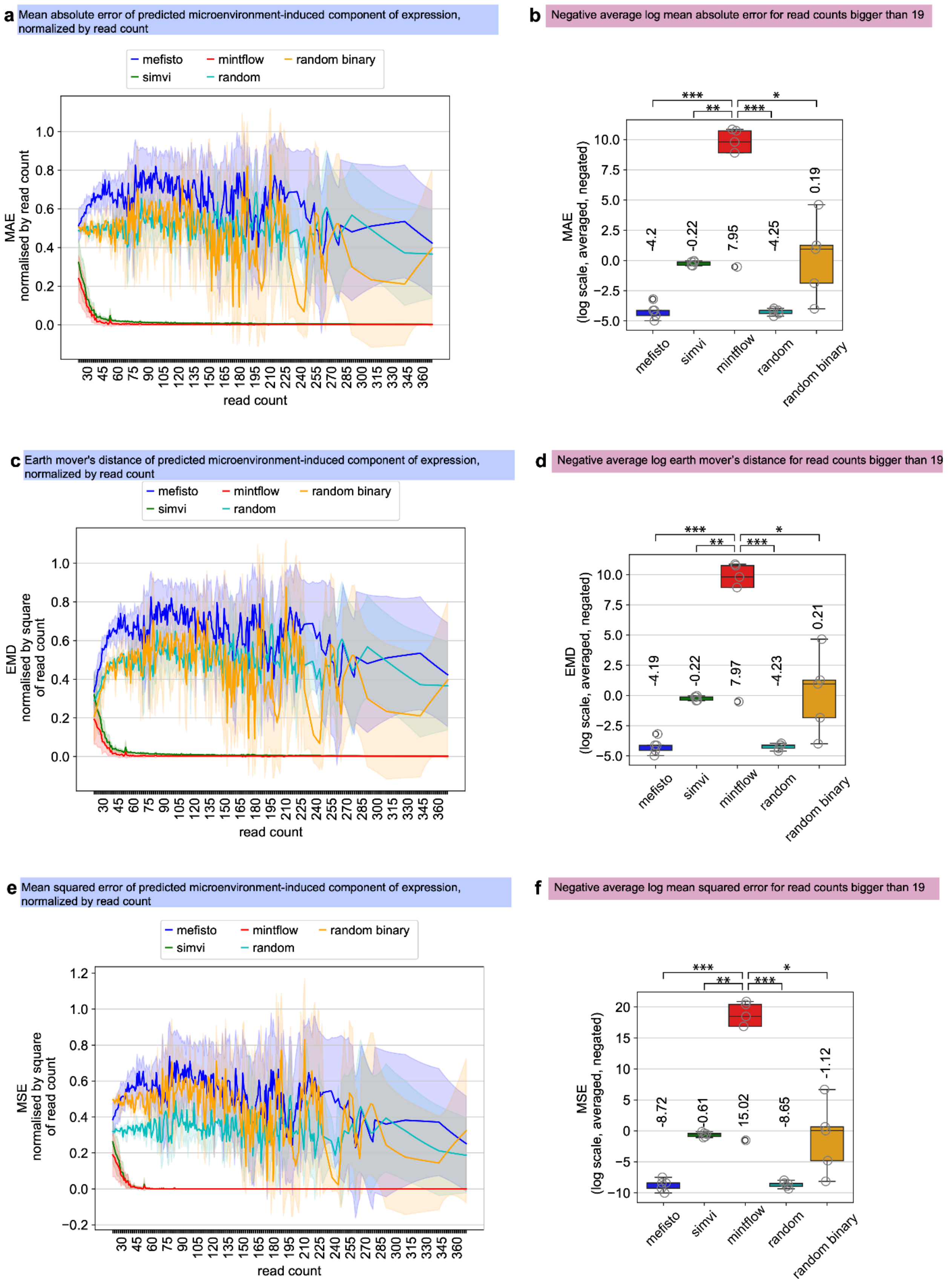
a
Mean absolute error of predicted microenvironment-induced component of expression, normalized by read count
mefisto
mintflow
random binary
simvi
random
MAE
normalised by read count
read count
b
Negative average log mean absolute error for read counts bigger than 19
MAE
(log scale, averaged, negated)
-4.2
-0.22
7.95
-4.25
0.19
mefisto
simvi
mintflow
random
random binary
c
Earth mover's distance of predicted microenvironment-induced component of expression, normalized by read count
mefisto
mintflow
random binary
simvi
random
EMD
normalised by square
of read count
read count
d
Negative average log earth mover's distance for read counts bigger than 19
EMD
(log scale, averaged, negated)
-4.19
-0.22
7.97
-4.23
0.21
mefisto
simvi
mintflow
random
random binary
e
Mean squared error of predicted microenvironment-induced component of expression, normalized by read count
mefisto
mintflow
random binary
simvi
random
MSE
normalised by square
of read count
read count
f
Negative average log mean squared error for read counts bigger than 19
MSE
(log scale, averaged, negated)
-8.72
-0.61
15.02
-8.65
-1.12
mefisto
simvi
mintflow
random
random binary

**Supplementary Fig. 3 | Method evaluation on simulated data (after discarding two cell type-homogeneous regions).** Evaluation of different methods in predicting the spatial component of expression. **a**, **c**, **e**, Mean absolute error (MAE), earth mover's distance (EMD), and mean squared error (MSE) for different read count values. **b**, **d**, **f**, The corresponding metrics for read counts > 19, log-transformed, averaged, and negated. Note that since for simulated data the ground truth intrinsic and microenvironment-induced components of expression are available, we were able to compute these metrics. This evaluation was done after two homogenous tissue regions, i.e. regions with cells of the same type, were discarded. Boxplot elements are defined as center line, median; box limits, upper and lower quartiles; whiskers, 1.5× interquartile range.

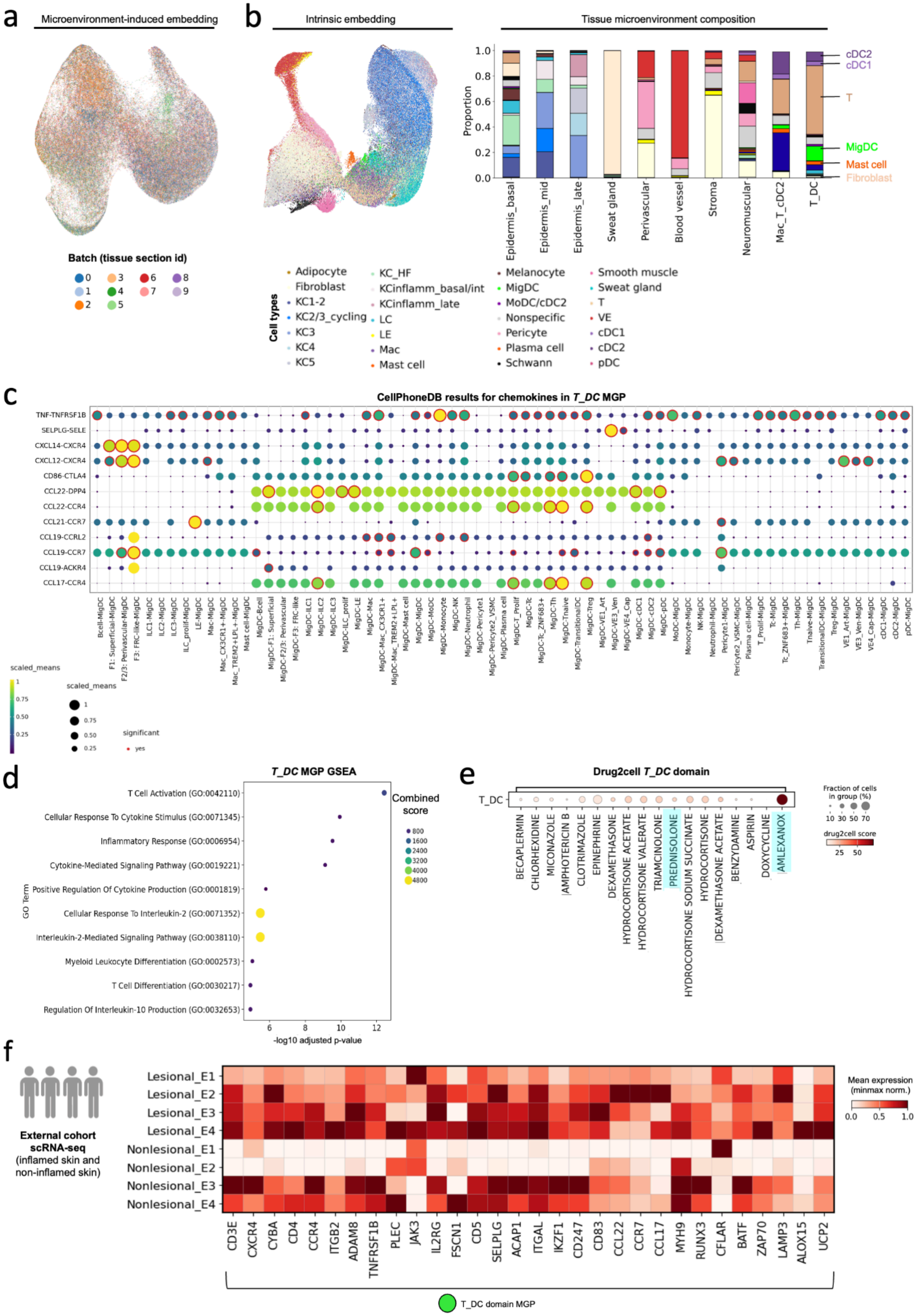
a
Microenvironment-induced embedding
Batch (tissue section id)
0 1 2 3 4 5 6 7 8 9
b
Intrinsic embedding
Tissue microenvironment composition
Proportion
1.0 0.8 0.6 0.4 0.2 0.0
Epidermis_basal
Epidermis_mid
Epidermis_late
Sweat gland
Perivascular
Blood vessel
Stroma
Neuromuscular
Mac_T_cDC2
T_DC
cDC2
cDC1
T
MigDC
Mast cell
Fibroblast
Cell types
Adipocyte
Fibroblast
KC1-2
KC2/3_cycling
KC3
KC4
KC5
KC_HF
KCinflamm_basal/int
KCinflamm_late
LC
LE
Mac
Mast cell
Melanocyte
MigDC
MoDC/cDC2
Nonspecific
Pericyte
Plasma cell
Schwann
Smooth muscle
Sweat gland
T
VE
cDC1
cDC2
pDC
c
CellPhoneDB results for chemokines in T_DC MGP
TNF-TNFRSF1B
SELPLG-SELE
CXCL14-CXCR4
CXCL12-CXCR4
CD86-CTLA4
CCL22-DPP4
CCL22-CCR4
CCL21-CCR7
CCL19-CCRL2
CCL19-CCR7
CCL19-ACKR4
CCL17-CCR4
scaled_means
significant
yes
d
T_DC MGP GSEA
T Cell Activation (GO:0042110)
Cellular Response To Cytokine Stimulus (GO:0071345)
Inflammatory Response (GO:0006954)
Cytokine-Mediated Signaling Pathway (GO:0019221)
Positive Regulation Of Cytokine Production (GO:0001819)
Cellular Response To Interleukin-2 (GO:0071352)
Interleukin-2-Mediated Signaling Pathway (GO:0038110)
Myeloid Leukocyte Differentiation (GO:0002573)
T Cell Differentiation (GO:0030217)
Regulation Of Interleukin-10 Production (GO:0032653)
GO Term
-log10 adjusted p-value
Combined score
800 1600 2400 3200 4000 4800
e
Drug2cell T_DC domain
T_DC
BECAPLERMIN
CHLORHEXIDINE
MICONAZOLE
AMPHOTERICIN B
CLOTRIMAZOLE
EPINEPHRINE
DEXAMETHASONE
HYDROCORTISONE ACETATE
HYDROCORTISONE VALERATE
TRIAMCINOLONE
PREDNISOLONE
HYDROCORTISONE SODIUM SUCCINATE
HYDROCORTISONE
DEXAMETHASONE ACETATE
BENZYDAMINE
ASPIRIN
DOXYCYCLINE
AMLEXANOX
Fraction of cells in group (%)
10 30 50 70
drug2cell score
25 50
f
External cohort scRNA-seq (inflamed skin and non-inflamed skin)
Lesional_E1
Lesional_E2
Lesional_E3
Lesional_E4
Nonlesional_E1
Nonlesional_E2
Nonlesional_E3
Nonlesional_E4
CD3E CXCR4 CYBA CD4 CCR4 ITGB2 ADAM8 TNFRSF1B PLEC JAK3 IL2RG FSCN1 CD5 SELPLG ACAP1 ITGAL IKZF1 CD247 CD83 CCL22 CCR7 CCL17 MYH9 RUNX3 CFLAR BATF ZAP70 LAMP3 ALOX15 UCP2
Mean expression (minmax norm.)
0.0 0.5 1.0
T_DC domain MGP

**Supplementary Fig. 4 | Additional atopic dermatitis analysis**. **a**, UMAP visualization of microenvironment-induced embedding colored by batch/tissue section. **b**, UMAP visualization of intrinsic embedding colored by cell type. Cellular composition of microenvironment domains. **c**, Cell-cell communication analysis results for genes in the *T_DC* MGP using atopic dermatitis scRNA-seq data (further details in Methods). **d**, Gene set enrichment analysis (GSEA) results for the *T_DC* MGP. **e**, Drug2cell results for genes in the *T_DC* domain for topical therapies. **f**, Expression of *T_DC* MGP in an external validation cohort of 4 patients using atopic dermatitis scRNA-seq data. MGP, Microenvironment-induced gene program.

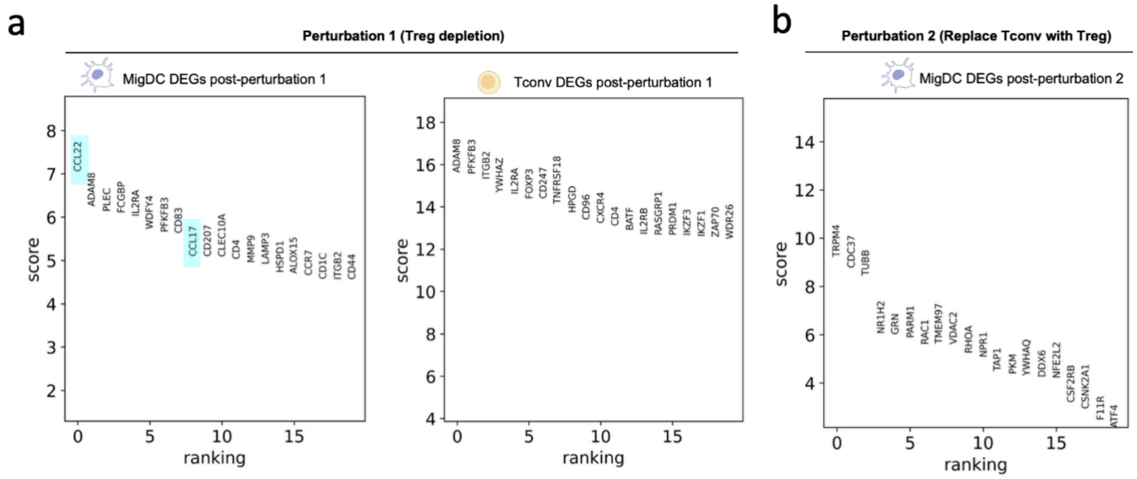

**Supplementary Fig. 5 | Additional perturbation analysis**. **a**, Top differentially expressed genes for MigDCs and Tconv cells in perturbation 1 (Treg depletion). **b**, Top differentially expressed genes for MigDCs in perturbation 2 (Treg augmentation).

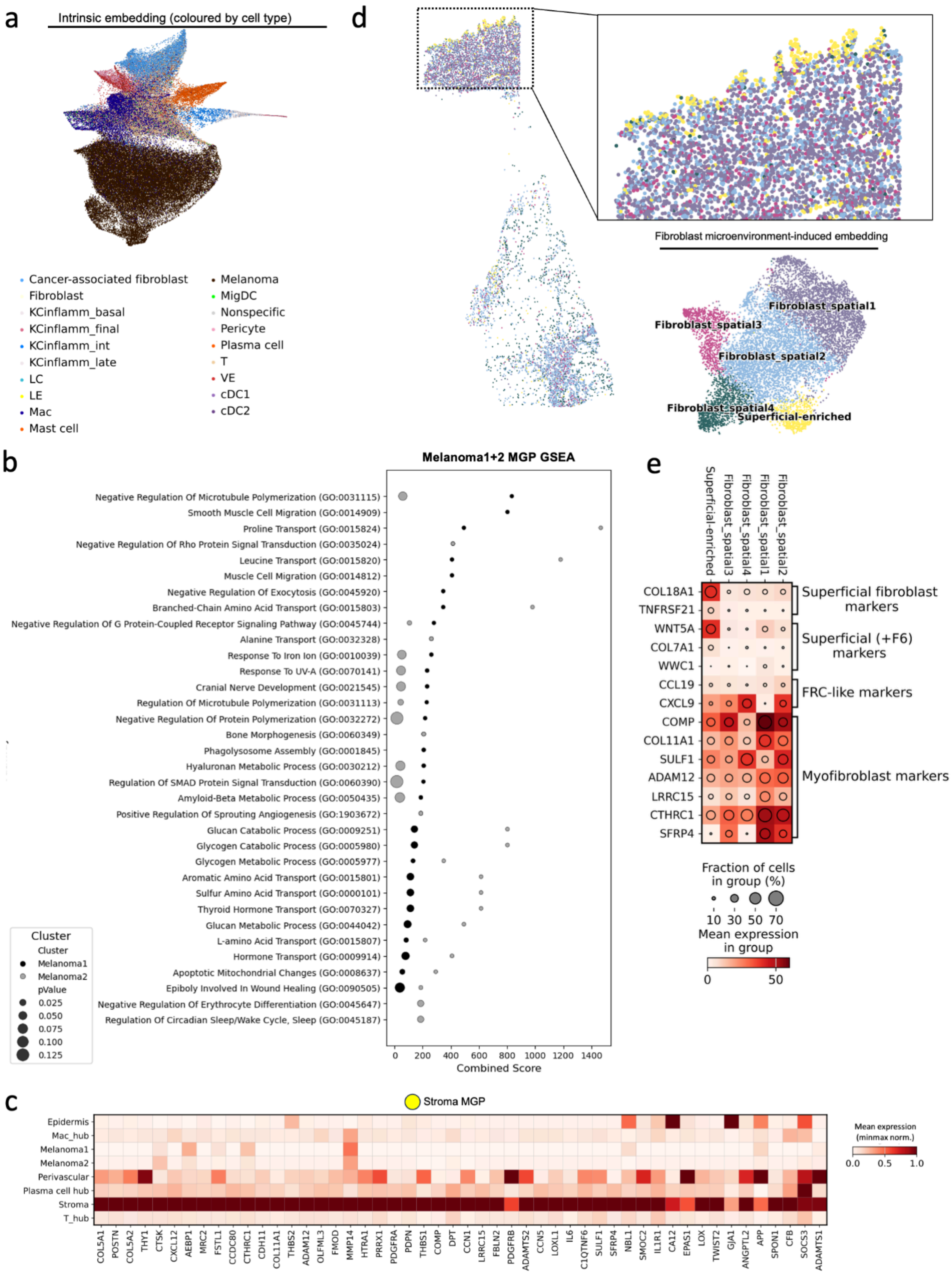
a
Intrinsic embedding (coloured by cell type)
Cancer-associated fibroblast
Fibroblast
KCinflamm_basal
KCinflamm_final
KCinflamm_int
KCinflamm_late
LC
LE
Mac
Mast cell
Melanoma
MigDC
Nonspecific
Pericyte
Plasma cell
T
VE
cDC1
cDC2
d
Fibroblast microenvironment-induced embedding
Fibroblast_spatial1
Fibroblast_spatial3
Fibroblast_spatial2
Fibroblast_spatial4
Superficial-enriched
b
Melanoma1+2 MGP GSEA
Negative Regulation Of Microtubule Polymerization (GO:0031115)
Smooth Muscle Cell Migration (GO:0014909)
Proline Transport (GO:0015824)
Negative Regulation Of Rho Protein Signal Transduction (GO:0035024)
Leucine Transport (GO:0015820)
Muscle Cell Migration (GO:0014812)
Negative Regulation Of Exocytosis (GO:0045920)
Branched-Chain Amino Acid Transport (GO:0015803)
Negative Regulation Of G Protein-Coupled Receptor Signaling Pathway (GO:0045744)
Alanine Transport (GO:0032328)
Response To Iron Ion (GO:0010039)
Response To UV-A (GO:0070141)
Cranial Nerve Development (GO:0021545)
Regulation Of Microtubule Polymerization (GO:0031113)
Negative Regulation Of Protein Polymerization (GO:0032272)
Bone Morphogenesis (GO:0060349)
Phagolysosome Assembly (GO:0001845)
Hyaluronan Metabolic Process (GO:0030212)
Regulation Of SMAD Protein Signal Transduction (GO:0060390)
Amyloid-Beta Metabolic Process (GO:0050435)
Positive Regulation Of Sprouting Angiogenesis (GO:1903672)
Glucan Catabolic Process (GO:0009251)
Glycogen Catabolic Process (GO:0005980)
Glycogen Metabolic Process (GO:0005977)
Aromatic Amino Acid Transport (GO:0015801)
Sulfur Amino Acid Transport (GO:0000101)
Thyroid Hormone Transport (GO:0070327)
Glucan Metabolic Process (GO:0044042)
L-amino Acid Transport (GO:0015807)
Hormone Transport (GO:0009914)
Apoptotic Mitochondrial Changes (GO:0008637)
Epiboly Involved In Wound Healing (GO:0090505)
Negative Regulation Of Erythrocyte Differentiation (GO:0045647)
Regulation Of Circadian Sleep/Wake Cycle, Sleep (GO:0045187)
Cluster
Cluster
Melanoma1
Melanoma2
pValue
0.025
0.050
0.075
0.100
0.125
0 200 400 600 800 1000 1200 1400
Combined Score
e
Superficial-enriched
Fibroblast_spatial3
Fibroblast_spatial4
Fibroblast_spatial1
Fibroblast_spatial2
COL18A1
TNFRSF21
WNT5A
COL7A1
WWC1
CCL19
CXCL9
COMP
COL11A1
SULF1
ADAM12
LRRC15
CTHRC1
SFRP4
Superficial fibroblast markers
Superficial (+F6) markers
FRC-like markers
Myofibroblast markers
Fraction of cells in group (%)
10 30 50 70
Mean expression in group
0 50
c
Stroma MGP
Epidermis
Mac_hub
Melanoma1
Melanoma2
Perivascular
Plasma cell hub
Stroma
T_hub
COL5A1 POSTN COL5A2 THY1 CTSK CXCL12 AEBP1 MRC2 FSTL1 CCDC80 CTHRC1 CDH11 COL11A1 THBS2 ADAM12 OLFML3 FMOD MMP14 HTRA1 PRRX1 PDGFRA PDPN THBS1 COMP DPT CCN1 LRRC15 FBLN2 PDGFRB ADAMTS2 CCN5 LOXL1 IL6 C1QTNF6 SULF1 SFRP4 NBL1 SMOC2 IL1R1 CA12 EPAS1 LOX TWIST2 GJA1 ANGPTL2 APP SPON1 CFB SOCS3 ADAMTS1
Mean expression (minmax norm.)
0.0 0.5 1.0

**Supplementary Fig. 6 | Additional melanoma analysis**. **a**, UMAP visualization of intrinsic embedding colored by cell type. **b**, Enriched Gene Ontology pathways from gene set enrichment analysis (GSEA) for the *Melanoma1* and *Melanoma2* microenvironment domains. **c**, Top 50 differential expressed genes in the *Stroma* MGP. **d**, Tissue colored by microenvironment-induced embedding clusters (left), and UMAP visualization of microenvironment-induced embedding (bottom right). **e**, Expression of fibroblast marker genes across different microenvironment-induced clusters, with enrichment of superficial markers. MGP, Microenvironment-induced Gene Program.

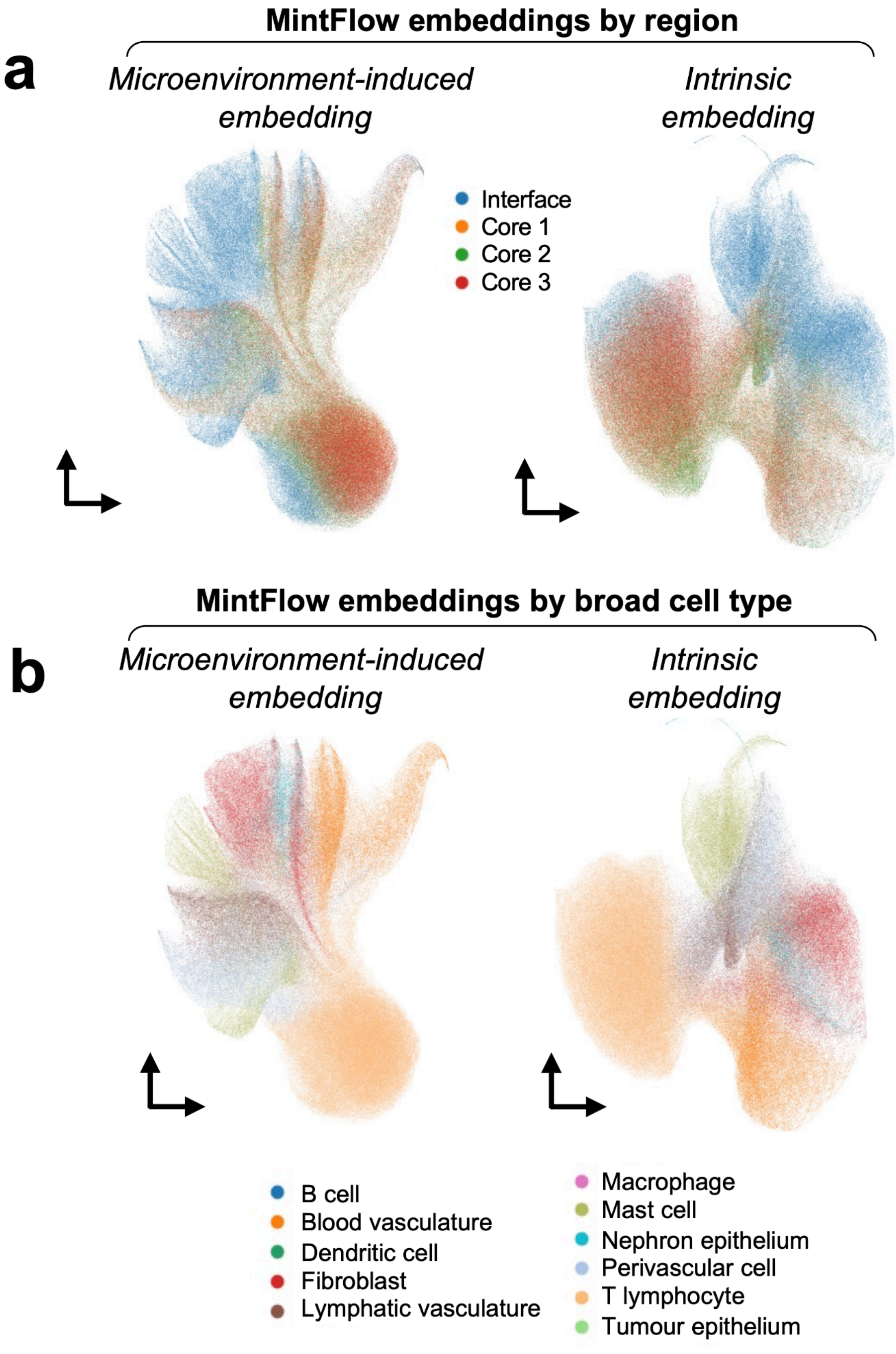
MintFlow embeddings by region
a
Microenvironment-induced embedding
Intrinsic embedding
Interface
Core 1
Core 2
Core 3
MintFlow embeddings by broad cell type
b
Microenvironment-induced embedding
Intrinsic embedding
B cell
Blood vasculature
Dendritic cell
Fibroblast
Lymphatic vasculature
Macrophage
Mast cell
Nephron epithelium
Perivascular cell
T lymphocyte
Tumour epithelium

**Supplementary Fig. 7 | MintFlow integration of regions and cell types in kidney cancer. a**, MintFlow embeddings grouped by region from the renal cell carcinoma (RCC) nephrectomy. The left UMAP is based on MintFlow microenvironment-induced embeddings and the right UMAP based on intrinsic embeddings. **b**, MintFlow embeddings grouped by broadly classified cell types within the RCC dataset. The left UMAP is based on MintFlow microenvironment-induced embeddings and the right UMAP based on intrinsic embeddings.

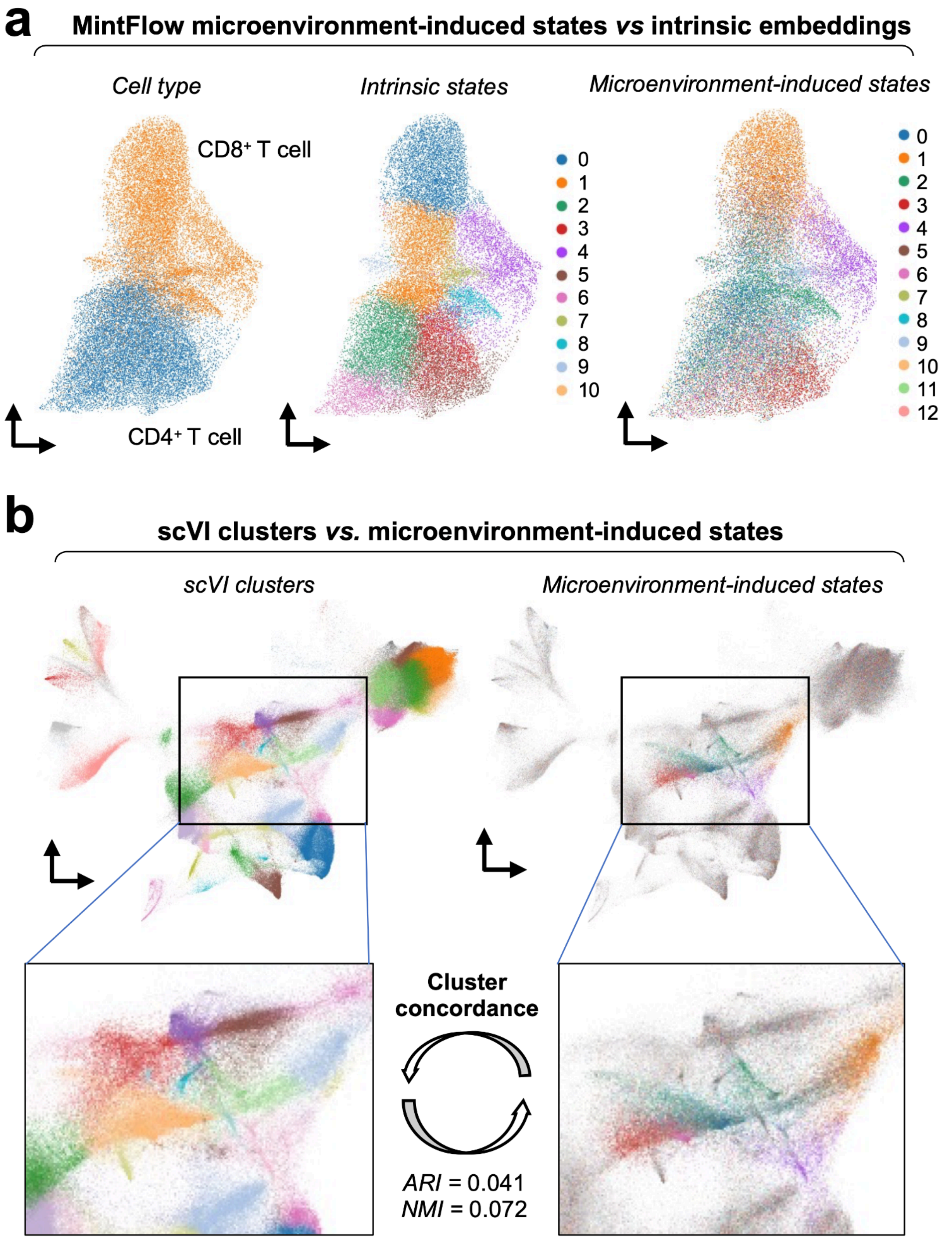

**Supplementary Fig. 8 | MintFlow microenvironment-induced cell states in kidney cancer**

**are different from intrinsic states and original clusters. a**, UMAPs showing intrinsic embeddings for T cells in renal cell carcinoma (RCC) generated by MintFlow. The left UMAP is grouped by original cell type, the middle by intrinsic T cell states and the right by microenvironment-induced T cell states. **b**, Analysis of original clustering for the RCC dataset (left UMAP), with the right UMAP showing the microenvironment-induced T cell states superimposed. There is low concordance of original clusters and microenvironment-induced T cell states (ARI; Adjusted Rand Index = 0.041, Normalized Mutual Information = 0.072).

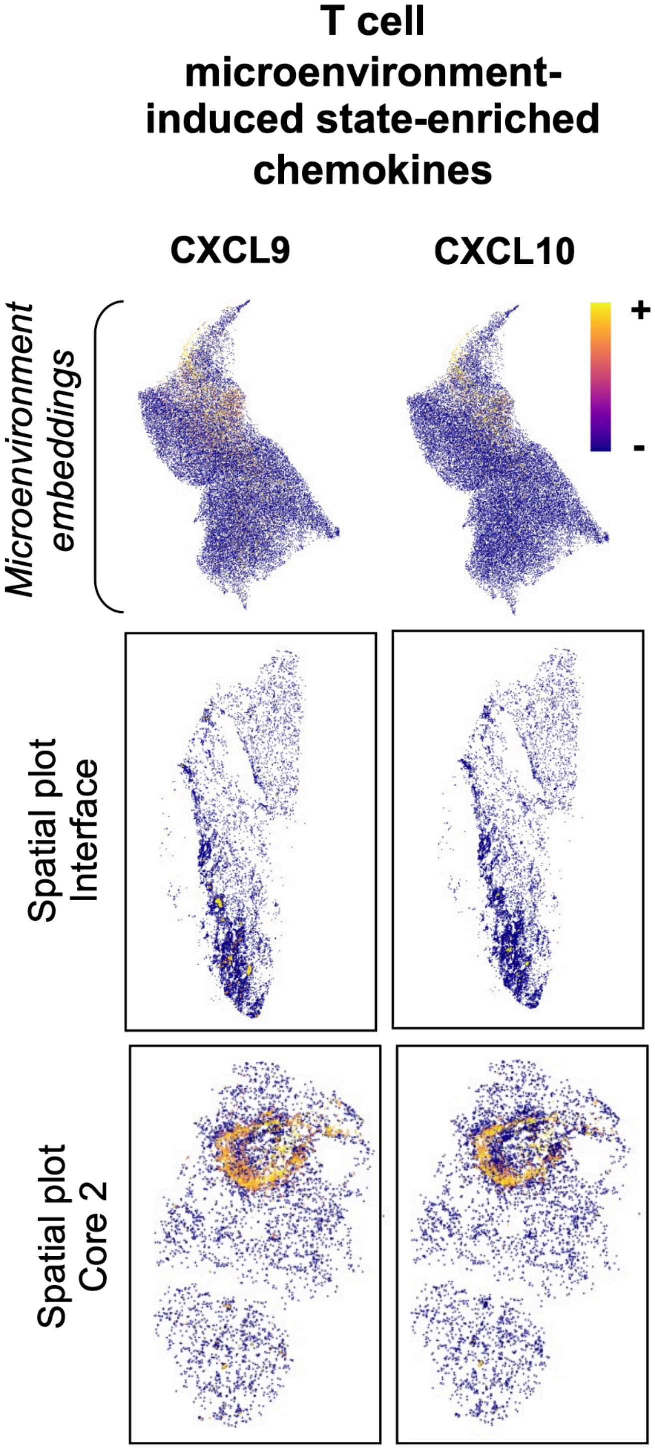

**Supplementary Fig. 9 | Microenvironment-induced T cell state-enriched chemokines in kidney cancer.** Analysis of expression of specific chemokines enriched in TLS T cell (*CXCL9, CXCL10*) across spatial plots for each region within the renal cell carcinoma (RCC) dataset.

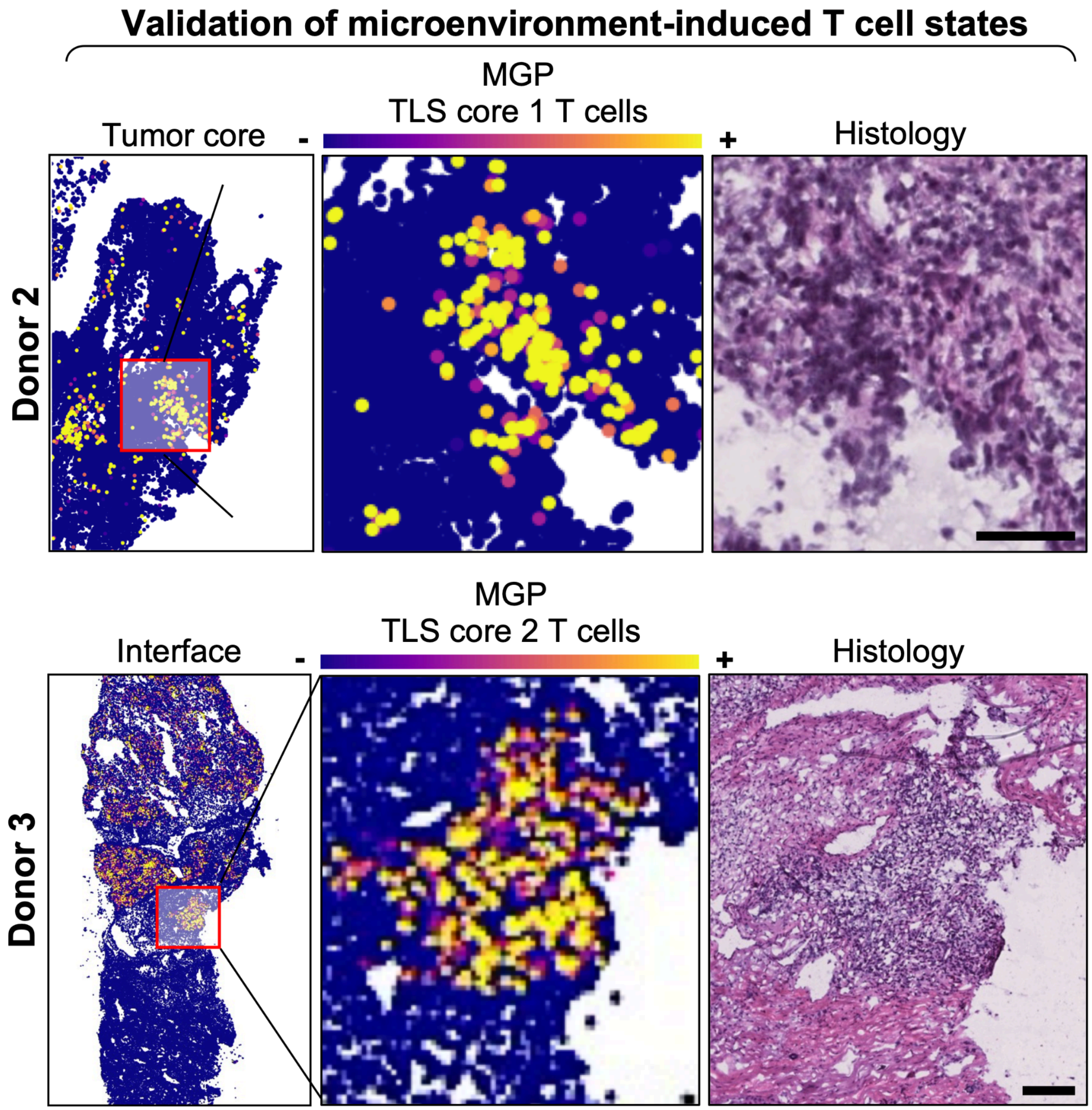

**Supplementary Fig. 10 | Validation of MintFlow microenvironment-induced cell states across multiple patients.** A microenvironment-induced gene program (MGP) was computed for each microenvironment-induced T cell state and its expression was analyzed across Xenium data from multiple donors not used for training MintFlow. The tertiary lymphoid structure (TLS) T cell core 1 gene program was recovered in Donor 2 (top left), with corresponding histology showing lymphocytic infiltration (top right). The equivalent has been done for expression of the TLS T cell core 2 gene program (bottom left), with corresponding TLS architecture observed in histology (bottom right). Scale bars in histology images = 200 μm.

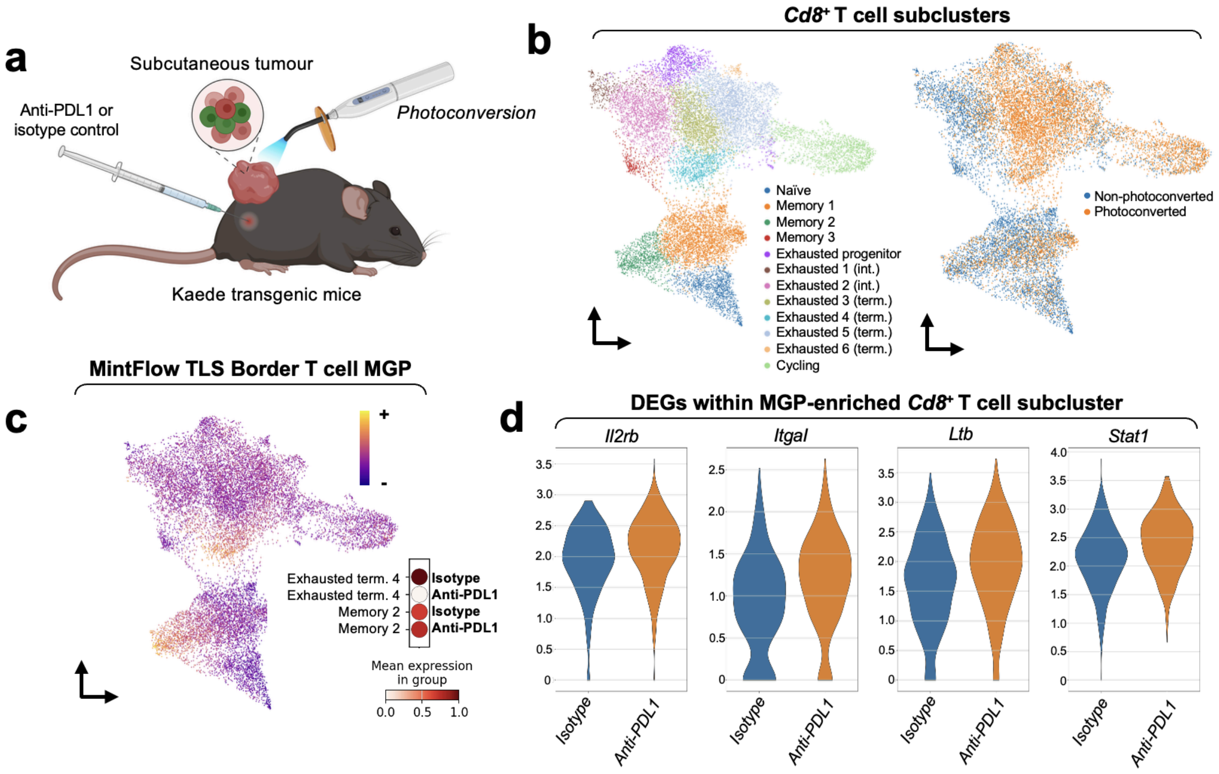

**Supplementary Fig. 11 | Validation of MintFlow-derived TLS Border T cell program in an in vivo mouse tumor model. a**, Schematic of in vivo experiment in Kaede mice: colorectal cancer cells were injected subcutaneously, followed by treatment with anti-PD-L1 antibody or isotype control. Photoconversion was used to distinguish tumor-resident (photoconverted) from recently infiltrating (non-photoconverted) immune cells. **b**, UMAP of CD8$^+$ T cells from tumors reveals distinct naïve, memory, cycling, and exhausted T cell subsets. **c**, Enrichment of the MintFlow-derived TLS Border T cell gene program across CD8$^+$ T cell subsets. The program was selectively expressed in memory and exhausted T cells, with exhausted T cells predominantly photoconverted, consistent with tumor residency. These exhausted cells had reduced expression of the MintFlow-derived TLS Border T cell gene program with anti-PD-L1 antibody treatment. **d**, Differential gene expression analysis of TLS Border-high exhausted CD8$^+$ T cells reveals upregulation of *Il2rb*, *Stat1*, and *Ltb* following anti-PD-L1 treatment, supporting checkpoint sensitivity of this spatially-defined, immunosuppressed T cell state.

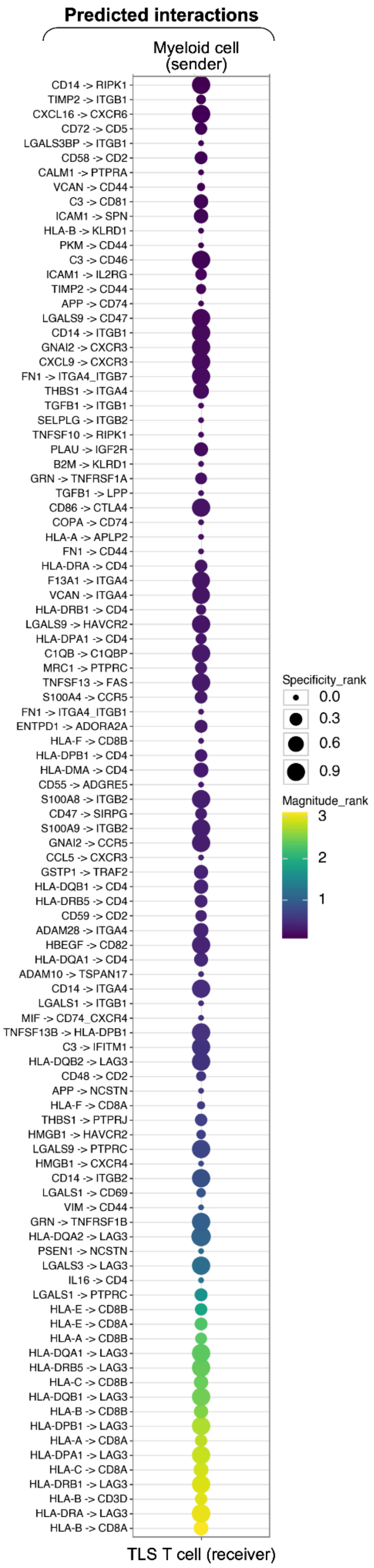
Predicted interactions
Myeloid cell (sender)
CD14 -> RIPK1
TIMP2 -> ITGB1
CXCL16 -> CXCR6
CD72 -> CD5
LGALS3BP -> ITGB1
CD58 -> CD2
CALM1 -> PTPRA
VCAN -> CD44
C3 -> CD81
ICAM1 -> SPN
HLA-B -> KLRD1
PKM -> CD44
C3 -> CD46
ICAM1 -> IL2RG
TIMP2 -> CD44
APP -> CD74
LGALS9 -> CD47
CD14 -> ITGB1
GNAI2 -> CXCR3
CXCL9 -> CXCR3
FN1 -> ITGA4_ITGB7
THBS1 -> ITGA4
TGFB1 -> ITGB1
SELPLG -> ITGB2
TNFSF10 -> RIPK1
PLAU -> IGF2R
B2M -> KLRD1
GRN -> TNFRSF1A
TGFB1 -> LPP
CD86 -> CTLA4
COPA -> CD74
HLA-A -> APLP2
FN1 -> CD44
HLA-DRA -> CD4
F13A1 -> ITGA4
VCAN -> ITGA4
HLA-DRB1 -> CD4
LGALS9 -> HAVCR2
HLA-DPA1 -> CD4
C1QB -> C1QBP
MRC1 -> PTPRC
TNFSF13 -> FAS
S100A4 -> CCR5
FN1 -> ITGA4_ITGB1
ENTPD1 -> ADORA2A
HLA-F -> CD8B
HLA-DPB1 -> CD4
HLA-DMA -> CD4
CD55 -> ADGRE5
S100A8 -> ITGB2
CD47 -> SIRPG
S100A9 -> ITGB2
GNAI2 -> CCR5
CCL5 -> CXCR3
GSTP1 -> TRAF2
HLA-DQB1 -> CD4
HLA-DRB5 -> CD4
CD59 -> CD2
ADAM28 -> ITGA4
HBEGF -> CD82
HLA-DQA1 -> CD4
ADAM10 -> TSPAN17
CD14 -> ITGA4
LGALS1 -> ITGB1
MIF -> CD74_CXCR4
TNFSF13B -> HLA-DPB1
C3 -> IFITM1
HLA-DQB2 -> LAG3
CD48 -> CD2
APP -> NCSTN
HLA-F -> CD8A
THBS1 -> PTPRJ
HMGB1 -> HAVCR2
LGALS9 -> PTPRC
HMGB1 -> CXCR4
CD14 -> ITGB2
LGALS1 -> CD69
VIM -> CD44
GRN -> TNFRSF1B
HLA-DQA2 -> LAG3
PSEN1 -> NCSTN
LGALS3 -> LAG3
IL16 -> CD4
LGALS1 -> PTPRC
HLA-E -> CD8B
HLA-E -> CD8A
HLA-A -> CD8B
HLA-DQA1 -> LAG3
HLA-DRB5 -> LAG3
HLA-C -> CD8B
HLA-DQB1 -> LAG3
HLA-B -> CD8B
HLA-DPB1 -> LAG3
HLA-A -> CD8A
HLA-DPA1 -> LAG3
HLA-C -> CD8A
HLA-DRB1 -> LAG3
HLA-B -> CD3D
HLA-DRA -> LAG3
HLA-B -> CD8A
TLS T cell (receiver)
Specificity_rank
0.0
0.3
0.6
0.9
Magnitude_rank
3
2
1

**Supplementary Fig. 12 | Analysis of MintFlow-informed cell-cell communication networks between TLS macrophages and T cells within single-cell RNA-seq data of kidney cancer.** Dot plot of predicted cell-cell interactions, generated with LIANA, from myeloid cells signaling to TLS T cells within the scRNA-seq data.

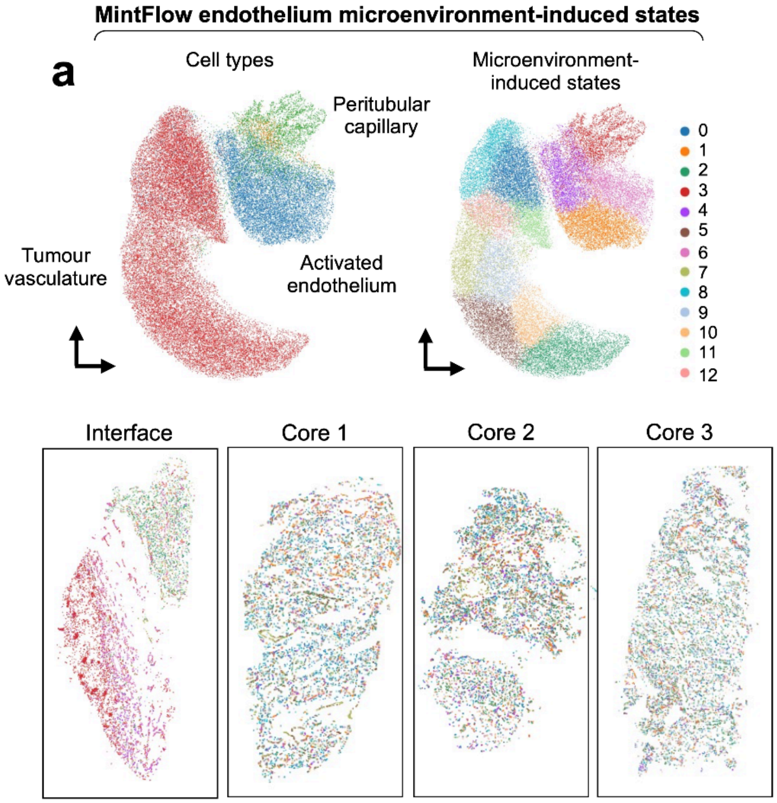
MintFlow endothelium microenvironment-induced states
a
Cell types
Peritubular capillary
Tumour vasculature
Activated endothelium
Microenvironment-induced states
0
1
2
3
4
5
6
7
8
9
10
11
12
Interface
Core 1
Core 2
Core 3

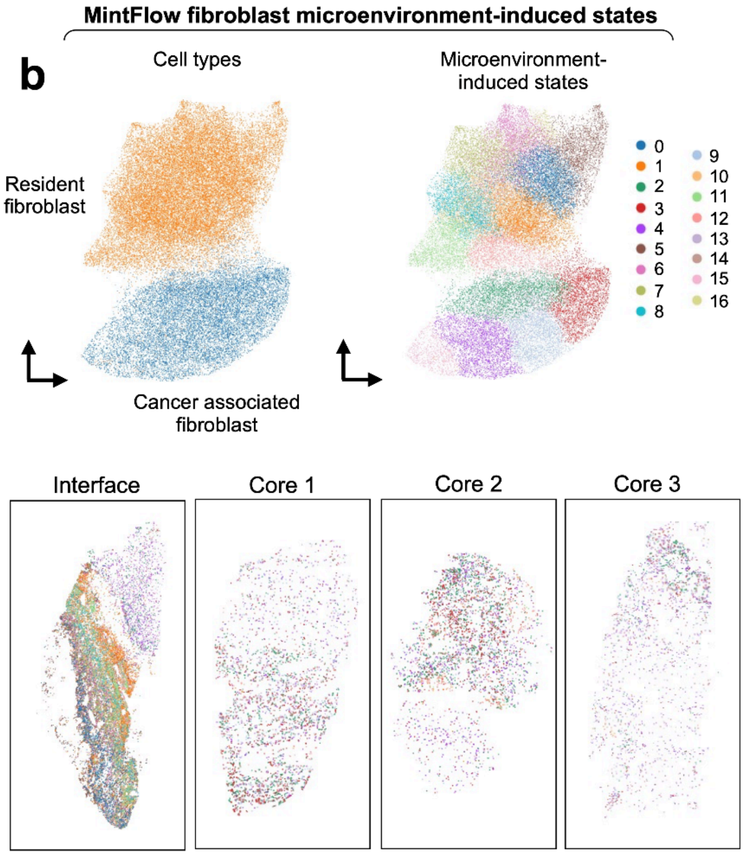
MintFlow fibroblast microenvironment-induced states
b
Cell types
Resident fibroblast
Cancer associated fibroblast
Microenvironment-induced states
Interface
Core 1
Core 2
Core 3

**Supplementary Fig. 13 | Spatial analysis of microenvironment-induced cell states for endothelium and fibroblasts in kidney cancer. a**, The left UMAP shows original endothelial cell types, plotted using microenvironment-induced embeddings, for endothelium within the RCC dataset. The right UMAP shows unsupervised clustering of microenvironment-induced cell states for endothelial cells. Below, the spatial plots show the endothelium microenvironment-induced states across each spatial region within the RCC tissue. **b**, The left UMAP shows original fibroblast types, plotted using microenvironment-induced embeddings, for fibroblast within the RCC dataset. The right UMAP shows unsupervised clustering of microenvironment-induced cell states for fibroblasts. Below, the spatial plots show the fibroblast microenvironment-induced states across each spatial region within the RCC tissue.

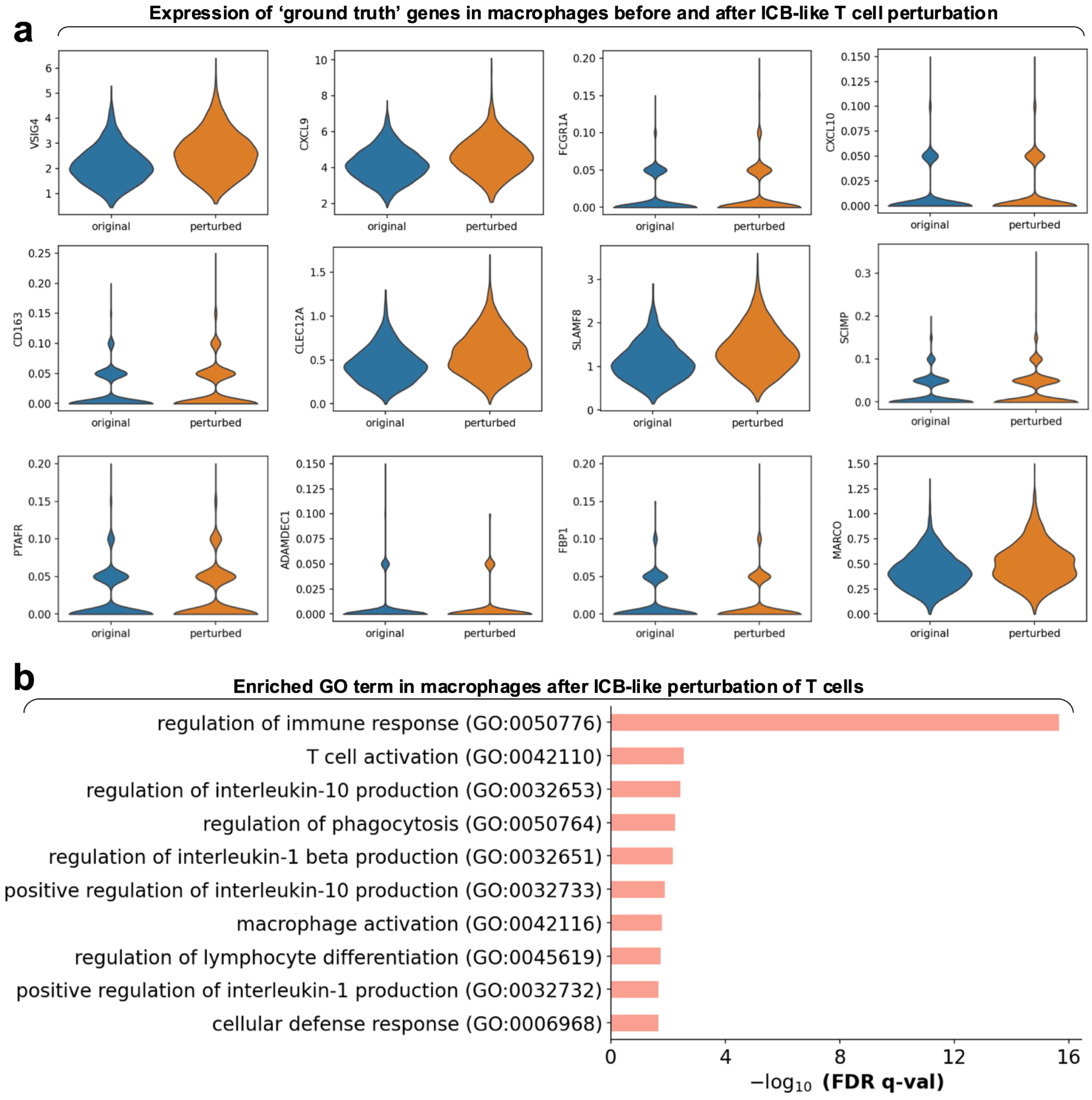

**Supplementary Fig. 14 | Assessing the biological validity of *in silico* T cell replacement within the TLS of ccRCC. a**, Violin plots showing the 12 individual ‘ground truth’ genes, derived from a previously published scRNA-seq study containing macrophages exposed to immune checkpoint blockade (ICB). The expression of these genes was compared in macrophages before or after replacement of all TLS T cells with T cells corresponding to a post-ICB signature derived from the same study. **b**, Gene set enrichment analysis (GSEA) for the top differentially expressed genes in perturbed macrophages. Top terms were listed by -log10 of their false discovery rate (FDR) q-value.

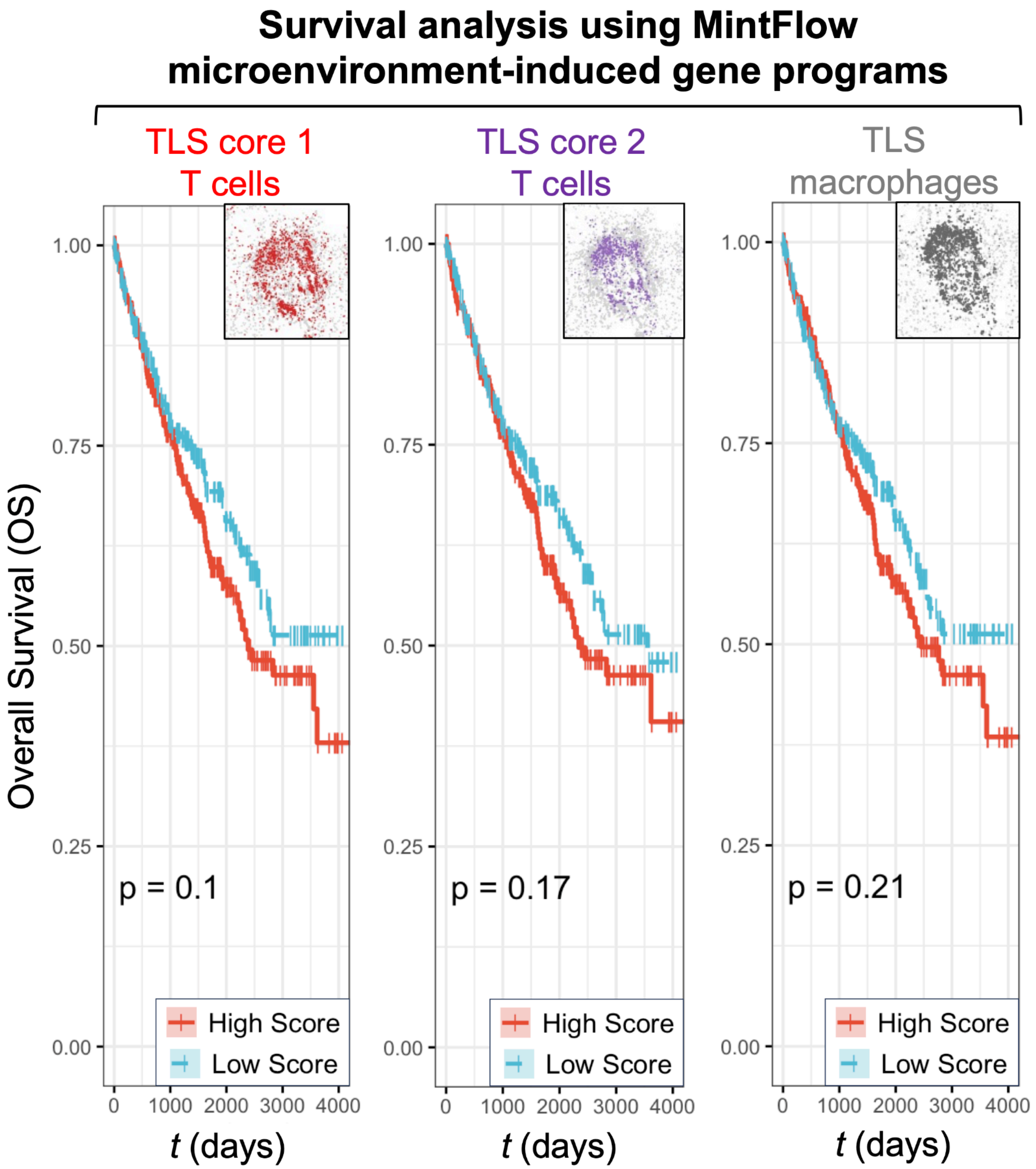

**Supplementary Fig. 15 | Survival analysis using microenvironment-induced T cell gene programs imputed by MintFlow.** Kaplan-Meier curves of patient survival across a bulk RNA-seq cohort of 606 patients derived from The Cancer Gene Atlas (TCGA). For each microenvironment-induced gene program (MGP), a 'high' score (red line) and a 'low' score (blue) line is assigned, and overall survival (OS) is plotted against time *t* in days. Neither TLS Core 1 ($p$ = 0.1), TLS Core 2 ($p$ = 0.17) nor TLS macrophages ($p$ = 0.21) were significantly associated with better or worse survival within this cohort.

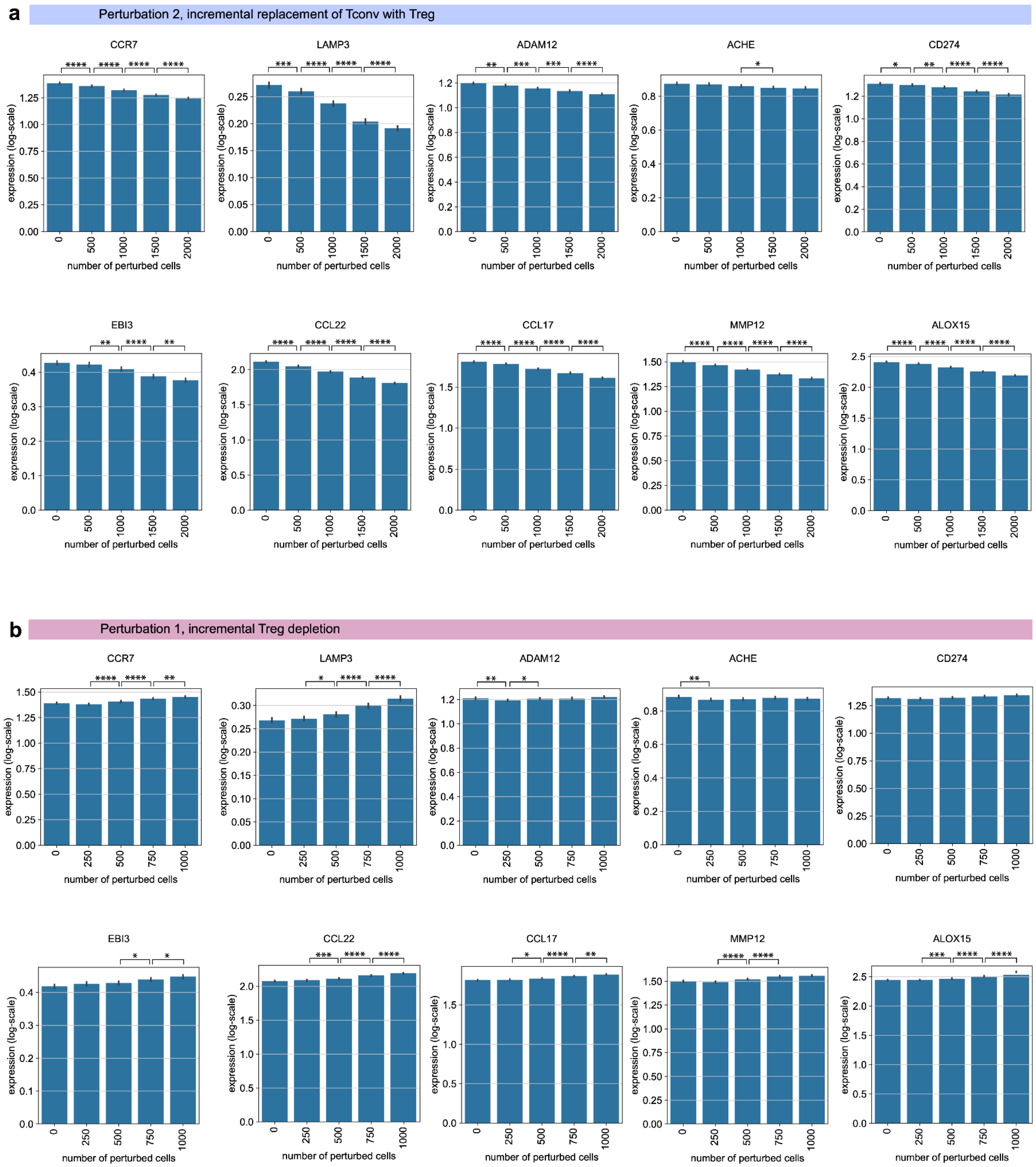

**Supplementary Fig. 16 | Incrementally applying MintFlow *in silico* perturbation has an incremental effect on generated gene expression vectors. a**, Perturbation 2 (Treg augmentation: replace Tconv cells with Tregs) as described in Fig. 3. **b**, Perturbation 1 (Treg depletion) as described in Fig. 3. Four, three, two, and one asterisks respectively indicate p-values less than or equal to 0.0001, 0.001, 0.01, and 0.05 in a t-test.

#### MintFlow modeling framework

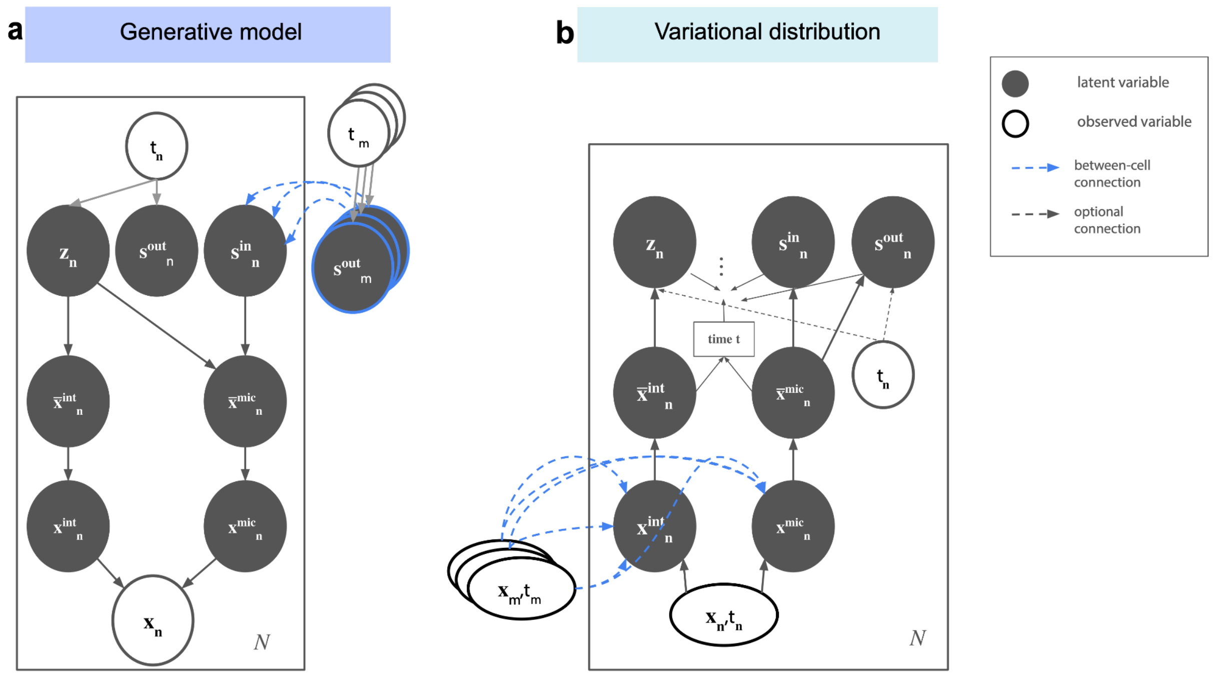

**Supplementary Fig. 17 | MintFlow modeling framework. a**, The proposed generative model. **b**, The variational distribution. Observed and latent variables are colored in white and black, respectively.

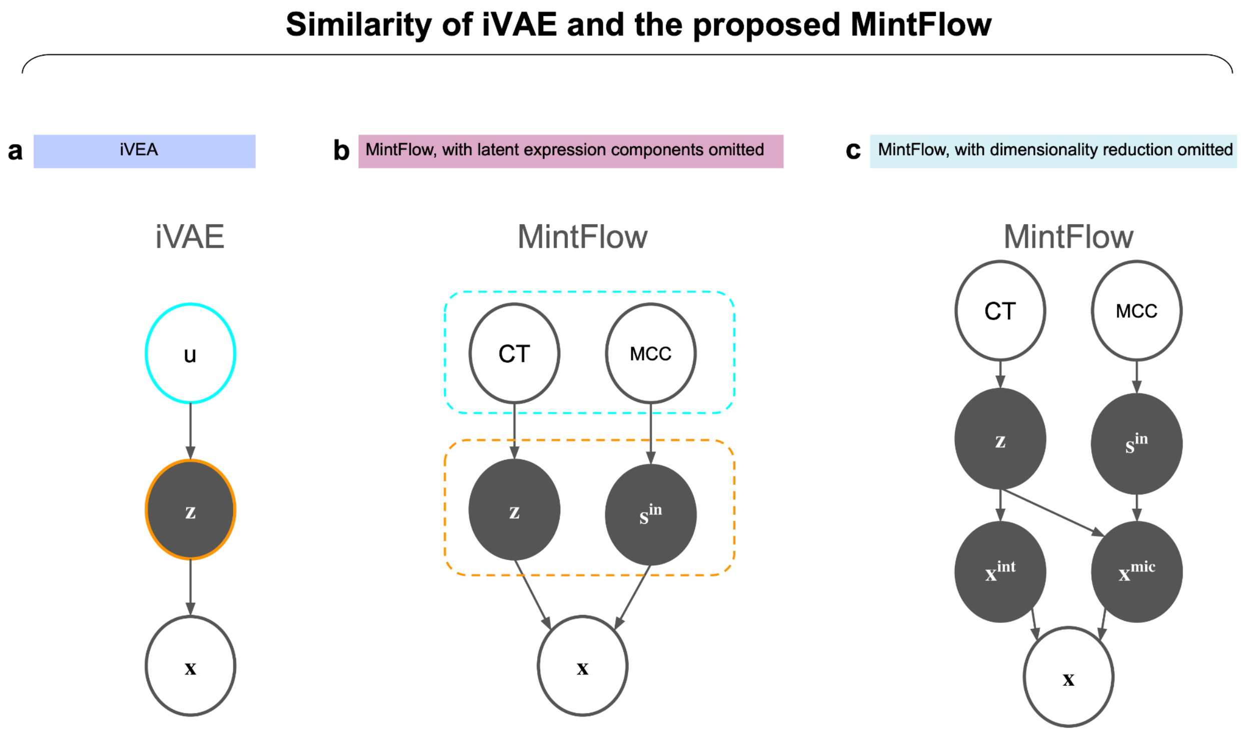

**Supplementary Fig. 18 | Similarity of iVAE and the proposed MintFlow**. Similarity of iVAE[1] (**a**) and our MintFlow method (**b**,**c**) in putting label-informed priors on latent embeddings. The cyan and orange outlines specify the additional side information to enable identifiability and latent vectors, respectively. MCC, microenvironment cell type composition; CT, cell type. **c**, A simplified version of MintFlow where no dimensionality reduction is performed, i.e. where the variables $\bar{x}_n^{int}$ and $\bar{x}_n^{mic}$ are omitted.

#### The general setting with inference objective connected to flow matching

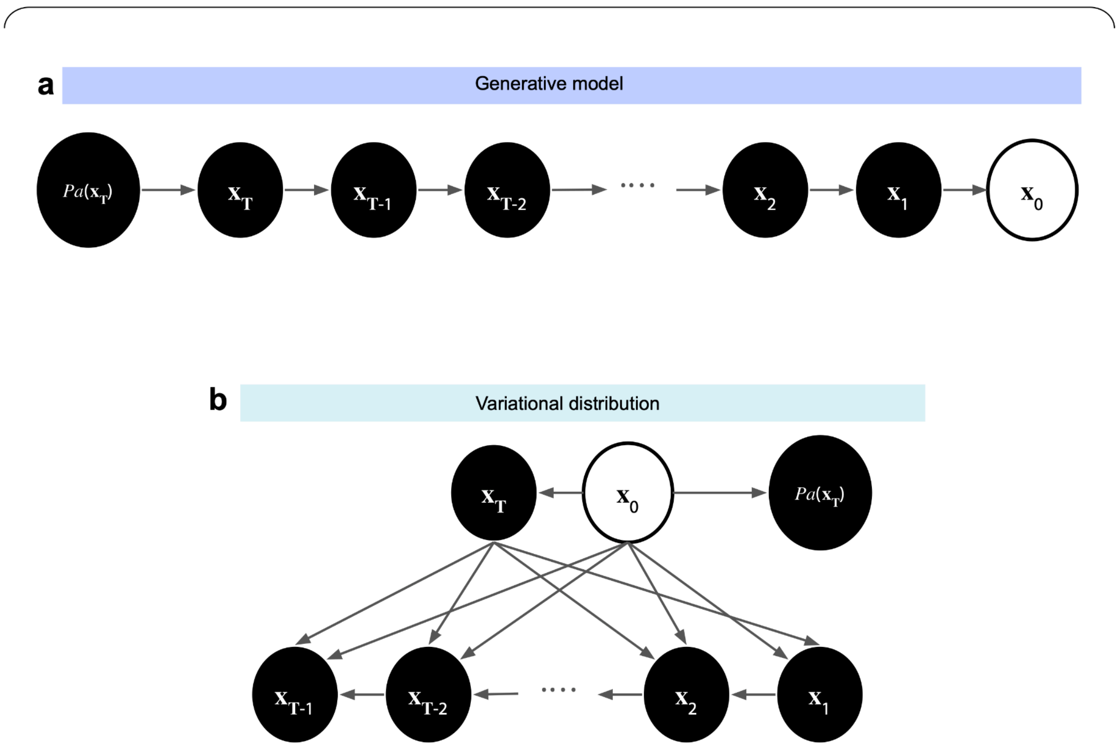

**Supplementary Fig. 19 | The general setting with inference objective connected to flow matching.** The general setting for which we show the connection of the usual ELBO and flow matching objective. Latent and observed variables are shown by dark and white ovals, respectively.

#### The default and customized samplers for data loaders

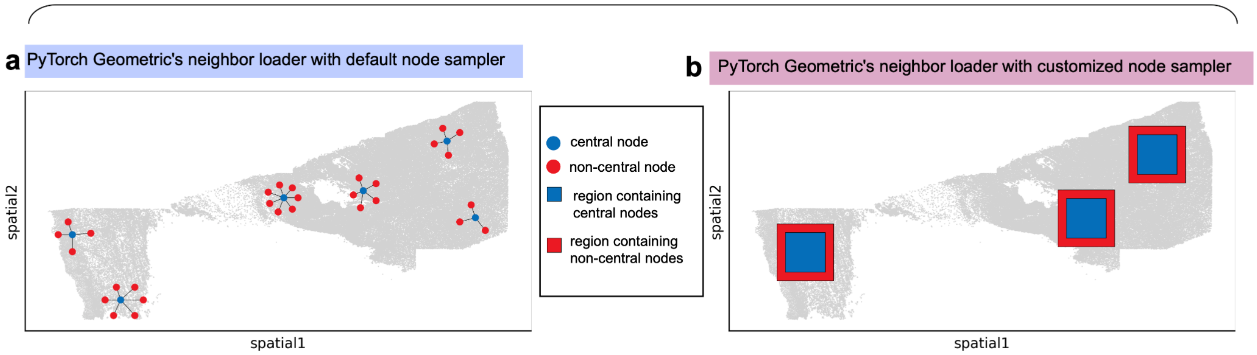

**Supplementary Fig. 20 | Neighbor loader design.** Subgraphs returned by PyTorch Geometric's neighbor loader with the default node sampler (**a**) and the customized sampler used by MintFlow (**b**). With the default sampler, the majority of subgraph nodes are non-central, while with the customized sampler it is vice versa.

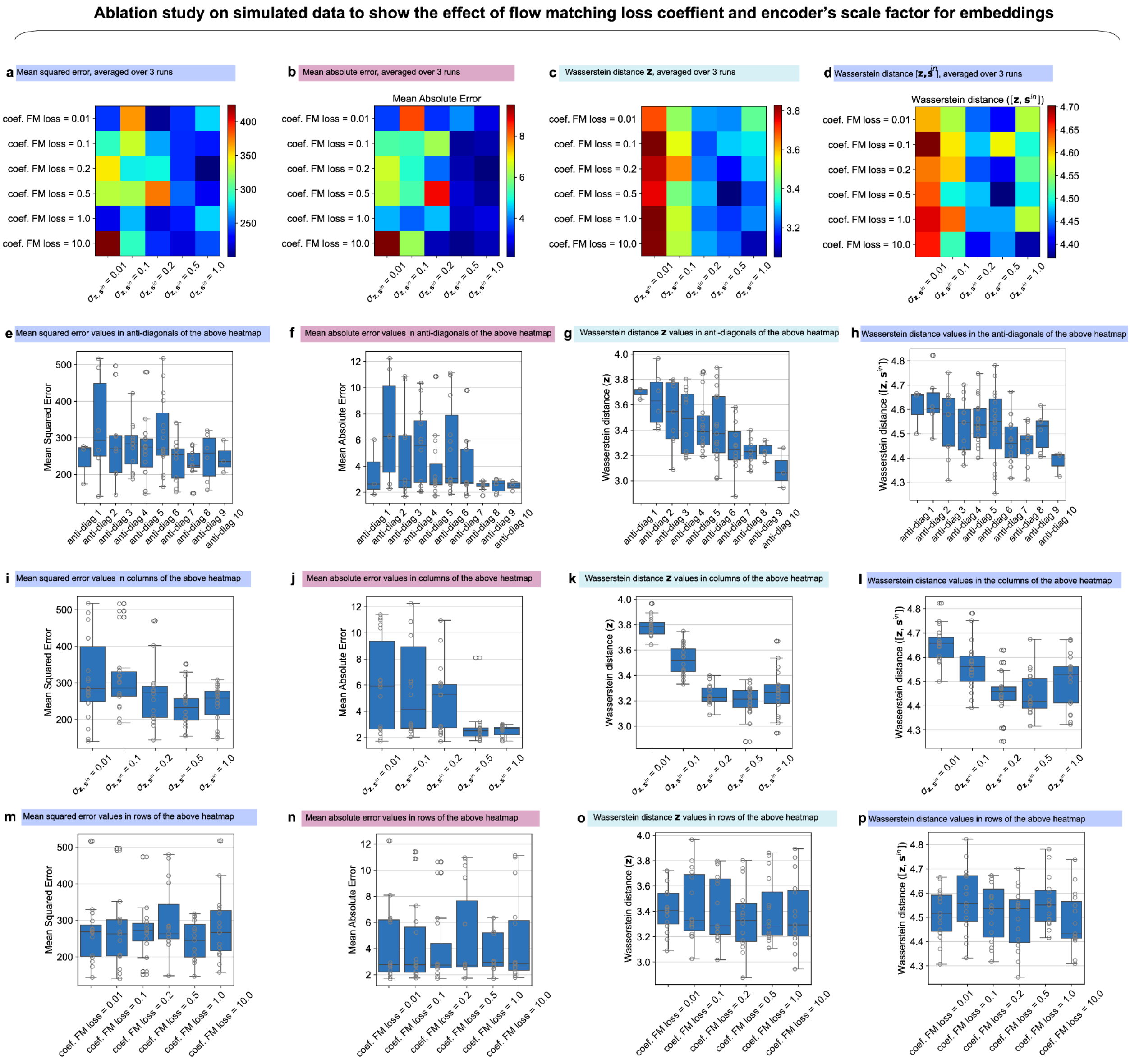

**Supplementary Fig. 21 | Loss ablation**. Ablation study to show the effect of flow matching loss on simulated data, as anticipated by Proposition 1. The importance coefficient of the flow matching objective and the scale factor for the encoders of $z$ and $s^{out}$ are varied, and for each setting of aforementioned values MintFlow is run 3 times. **a**, **b**, The closeness of MintFlow's predicted $x^{mic}$ to the ground-truth measured by MAE (mean absolute error) and MSE (mean squared error) averaged over the 3 runs. **c**, **d**, EMD (earth mover's distance) between $[z]\mapsto[\bar{x}^{int}]$ and $[z,s]\mapsto[\bar{x}^{int},\bar{x}^{mic}]$ averaged over the 3 runs. **e**, **f**, **g**, **h**, Box plots corresponding to the anti-diagonal traverse of the heatmaps in row one. Note that, unlike in the heatmaps, all 3 runs are included in the box plots. **i**, **j**, **k**, **l**, Box plots corresponding to the column by column traverse of the heatmaps. Note that, unlike in the heatmaps, all 3 runs are included in the boxplots. **m**, **n**, **o**, **p**, Box plots corresponding to the row by row traverse of the heatmaps. Note that, unlike in the heatmaps, all 3 runs are included in the box plots.

### Supplementary Notes

#### Supplementary Note 1: Related Work

In this section we briefly introduce some existing methods to identify microenvironment-induced cell states in spatial transcriptomics data. Then we compare their capabilities to those of the proposed MintFlow.

**MEFISTO**. MEFISTO[2] linearly decomposes the observed readout across different views (e.g. different tissue sections or different omics) to some factors and their corresponding weights. The factors are conditioned on the observed spatial or temporal coordinates non-linearly via Gaussian processes. The kernels of the Gaussian processes are capable of modeling view-view relations as well as dependence on spatial or temporal covariates. Factors whose corresponding weights are bigger than a threshold are considered to be significantly dependent on spatial and temporal covariates, and are used to extract the spatially- (i.e. microenvironment-) or temporally-induced component of the observed readout.

**SIMVI**. SIMVI[3] is an encoder/decoder based method that provides two separate embedding spaces to capture the intrinsic and spatial part of expression. The encoder for intrinsic embedding functions on each cell independently while the encoder for spatial embedding is a GNN (graph neural network) that incorporates neighbourhood information. The usual ELBO (evidence lower bound) is accompanied by a term that minimizes the mutual information between intrinsic and spatial embeddings, as well as an MMD (maximum mean discrepancy) regularizer to alleviate known collapse issues associated with VAE-based methods[4]. The obtained embedding spaces are analyzed to, e.g., identify cell populations and differential gene expression.

**DIALOGUE**. DIALOGUE[5] seeks to identify MCPs (MultiCellular Programs), defined as gene programs that function across different cell types in a concerted way to carry out a tissue-level biological task. DIALOGUE[5] achieves this in two stages. In the first stage some cell type-specific linear embeddings are trained to capture cell type-independent variation of readout data across different tissue samples or spatial microenvironments. To do so, cell type-specific linear encoders are trained such that the correlation of embeddings across different cell types are maximized. Intuitively, the first few linear embeddings are cell type-independent and are expected to capture the variation related to MCPs. In the second stage, DIALOGUE[5] identifies genes that mostly affect the learnt latent factors after taking into account the effect of other confounding factors such as age and sex. The identified MCPs are quantitatively validated by how well they are predictable from a cell's neighborhood expression as well as by spatial correlation.

**NCEM**. NCEM[6] is an encoder/decoder-based method whose encoder and decoder are conditioned on cell type labels, the abundance and/or interaction of cells of different types in a microenvironment, as well as batch labels. NCEM can take in and process this information in different ways. **Linear NCEM**: A design matrix containing the type and neighbors' cell types is fed to a linear decoder that reconstructs the read count matrix. **Non-linear NCEM**: neighborhood cell type information is fed to a graph neural network (GNN) and the resulting embedding is concatenated with cell type and batch label and is finally fed to an encoder. The decoder takes in the encoder's embedding and cell type and batch information, and reconstructs the gene expression vector of each cell. **CVAE NCEM**: A graph neural network (GNN) encodes cell type abundances in each cell's neighborhood which is then provided to both the encoder and the decoder along with cell type and batch labels. Besides this information, the encoder is provided with the observed gene expression vectors.

The method is evaluated by how well it can reconstruct or explain the read count variance on unseen data and in terms of the coefficient of determination.

At a high level, the goal of MintFlow overlaps with that of SIMVI[3], DIALOGUE[5], NCEM[6], and factor based methods like MEFISTO[2]. The key distinction of MintFlow with MEFISTO[2] is that the former is built around modeling cell cell interactions and inferring them, while the latter seeks to extract part of the observed expression which is highly dependent on and varies with the spatial location of cells. Notably, the dependence of the expression of a gene on cells' locations may not necessarily imply the gene is induced by microenvironment and can be due to, e.g., cells of a specific type exclusively overexpress the gene and form spatial patterns.

Both MintFlow and SIMVI[3] are encoder/decoder-based methods based on approximate Bayesian inference to disentangle microenvironment-induced part of observed expression from its intrinsic part. The identified microenvironment-induced component can be used for, e.g., cell population identification or patient stratification. One advantage of the proposed MintFlow is that it provides an explicit decomposition of the readout matrix to intrinsic and microenvironment-induced components, thereby providing explicit **interpretability** and removing the need for, e.g., the archetypal post-training analysis of the embeddings required by SIMVI[3]. Moreover, the scalability mechanism devised in MintFlow uniquely makes it scalable to datasets with several tissue sections and millions of cells.

Similar to MintFlow, NCEM[6] seeks to reconstruct the observed expression vectors from cell type labels in the index cell and its microenvironment as well as other covariates like batch labels. Unlike in MintFlow, in NCEM[6] there are no separate embeddings for intrinsic and microenvironment nor is the reconstruction loss confined to the microenvironment-induced part of the expression matrix. DIALOGUE's[5] approach to identify linear factors that capture multicellular programs, to some degree resembles MintFlow's $Objective_7$ and $Objective_8$ explained in the Methods section, with the difference that the former captures the variation which remains constant across different cell types while the latter captures the variation which remains the same for every cell type and across different microenvironments.

#### Supplementary Note 2: Proof of Proposition 1

In this section we restate Proposition 1 more formally and provide a proof for it.

**Proposition 1** *(more formal statement of Proposition 1) Let $\mathcal{Z} \times \mathcal{S}$ be measurable spaces with probability measure $dp(z,s)$. Suppose we have a known function $F$ on $\mathcal{Z} \times \mathcal{S}$. We aim to decompose $F$ into*

$$F(z,s) \;=\; H(z) \;+\; G(z,s), \tag{1}$$

*where $H$ and $G$ are the unknown functions to be found. Consider the following cost functional:*

$$\int_{\mathcal{Z}\times\mathcal{S}} c\big((z,s),(H(z),G(z,s))\big)\, dp(z,s) \;\;+\;\; \lambda \int_{\mathcal{Z}} c\big(z, H(z)\big)\, dp(z), \tag{2}$$

*where $\lambda > 0$ is a scalar constant, and $c$ is Euclidean distance (a cost function assumed to be* strictly convex*). There is exactly one global minimiser $\big(H^*, G^*\big)$ that minimises the cost functional of Eq. (2) subject to the constraint of Eq. (1).*

**Proof:**
Step 1: Rewriting $G$ in Terms of $H$. Since $F$ is known, from the decomposition of Eq. (1)

$$G(z,s) \;=\; F(z,s) \;-\; H(z).$$

Hence, once $H$ is given, there is no freedom left for $G$. Substituting $G(z,s) = F(z,s) - H(z)$ into Eq. (2), we separate the objective into two terms that depend *only* on $H$:

$$\underbrace{\int_{\mathcal{Z}\times\mathcal{S}} c\Big(\,(z,s),\; \big(H(z),\; F(z,s) - H(z)\big)\Big)\, dp(z,s)}_{\Psi(H)} \;+\; \underbrace{\lambda \int_{\mathcal{Z}} c\big(z,\; H(z)\big)\, dp(z)}_{\Phi(H)}.$$

We define the functionals

$$\Psi(H) \;=\; \int_{\mathcal{Z}\times\mathcal{S}} c\Big(\,(z,s),\; \big(H(z),\; F(z,s) - H(z)\big)\Big)\, dp(z,s), \qquad \Phi(H) \;=\; \lambda \int_{\mathcal{Z}} c\big(z, H(z)\big)\, dp(z). \tag{3}$$

Our problem becomes

$$\inf_{H} \Big\{\, \Psi(H) + \Phi(H) \Big\}.$$

Step 2: Strict Convexity of $\Psi$ and $\Phi$. We now show that $\Psi(H)$ and $\Phi(H)$ are both strictly convex in $H$.
Strict convexity of $\Psi$. Recall

$$\Psi(H) \;=\; \int_{\mathcal{Z}\times\mathcal{S}} c\Big(\,(z,s),\; \big(H(z),\; F(z,s) - H(z)\big)\Big)\, dp(z,s).$$

Fix two distinct functions $H_1$ and $H_2$, and define for $\alpha \in (0,1)$

$$H_\alpha(z) \;=\; \alpha\, H_1(z) + (1-\alpha)\, H_2(z).$$

Set

$$(h_1, g_1) = \big(H_1(z),\, F(z,s) - H_1(z)\big), \quad (h_2, g_2) = \big(H_2(z),\, F(z,s) - H_2(z)\big),$$

and

$$(h_\alpha, g_\alpha) = \big(H_\alpha(z),\, F(z,s) - H_\alpha(z)\big) = \alpha\,(h_1, g_1) + (1-\alpha)\,(h_2, g_2).$$

By the assumed strict convexity of $c((z,s),(\cdot,\cdot))$ in its second argument, we have

$$c\big((z,s),(h_\alpha,g_\alpha)\big) \;<\; \alpha\, c\big((z,s),(h_1,g_1)\big) \;+\; (1-\alpha)\, c\big((z,s),(h_2,g_2)\big),$$

for all $(z,s)$ where $(h_1,g_1) \neq (h_2,g_2)$. Integrating over $(z,s)$ via $dp(z,s)$, we obtain:

$$\Psi(H_\alpha) \;=\; \int c\big((z,s),(h_\alpha,g_\alpha)\big)\, dp(z,s) \;<\;$$
$$\alpha \int c\big((z,s),(h_1,g_1)\big)\, dp(z,s) \;+\; (1-\alpha) \int c\big((z,s),(h_2,g_2)\big)\, dp(z,s),$$

i.e.,

$$\Psi(H_\alpha) \;<\; \alpha\, \Psi(H_1) + (1-\alpha)\, \Psi(H_2),$$

as long as $\{z : H_1(z) \neq H_2(z)\}$ has positive measure. This proves $\Psi(H)$ is *strictly convex* in $H$.

Strict convexity of $\Phi$. We also have

$$\Phi(H) \;=\; \lambda \int_{\mathcal{Z}} c\big(z, H(z)\big)\, dp(z).$$

For each fixed $z$, the map $h \mapsto c(z,h)$ is strictly convex by assumption. Thus we can repeat the same argument:

$$H_\alpha(z) = \alpha\, H_1(z) + (1-\alpha)\, H_2(z) \quad \Longrightarrow \quad c\big(z, H_\alpha(z)\big) \;<\; \alpha\, c\big(z, H_1(z)\big) + (1-\alpha)\, c\big(z, H_2(z)\big),$$

for almost every $z$. Integrating over $dp(z)$ and multiplying by $\lambda$, we obtain:

$$\Phi(H_\alpha) \;<\; \alpha\, \Phi(H_1) + (1-\alpha)\, \Phi(H_2),$$

as long as $\{z : H_1(z) \neq H_2(z)\}$ has positive measure. Hence $\Phi(H)$ is strictly convex in $H$.

Step 3: Uniqueness of the Minimizer. Since $\Psi(H)$ and $\Phi(H)$ are both strictly convex, their sum $\Psi(H) + \Phi(H)$ is strictly convex as well. That is, for any distinct $H_1$ and $H_2$, as long as $\{z : H_1(z) \neq H_2(z)\}$ has positive measure,

$$\big(\Psi + \Phi\big)(H_\alpha) \;<\; \alpha\,(\Psi + \Phi)(H_1) \;+\; (1-\alpha)\,(\Psi + \Phi)(H_2),$$

for all $\alpha \in (0,1)$.

Step 4: Finally, uniqueness of $H^\star$ follows from strict convexity of $\Psi + \Phi$, and uniqueness of $G^\star$ follows from uniqueness of $H^\star$ and definition of $G$.

### Supplementary Note 3: Details of the Generative Process

In Methods we provided a big picture of the model and its assumptions. Here we provide the details of the generative process, which is as follows:

- $p(\boldsymbol{z}_n|\boldsymbol{t}_n) = \mathcal{N}\big(\boldsymbol{z}_n \ ; \ \boldsymbol{\mu}_{uz}^T\boldsymbol{t}_n \ , \ exp(\boldsymbol{\sigma_{uz}}(\boldsymbol{t}_n))\big)$.
- $p(\boldsymbol{s}_n^{out}|\boldsymbol{t}_n) = \mathcal{N}\big(\boldsymbol{s}_n^{out} \ ; \ \boldsymbol{\mu}_{us}^T\boldsymbol{t}_n \ , \ exp(\boldsymbol{\sigma_{us}}(\boldsymbol{t}_n))\big)$.
- $p\big(\boldsymbol{s}_n^{in}|\{\boldsymbol{s}_m^{out}|m \in Neigh(n)\}\big) = \mathcal{N}(\boldsymbol{s}_n^{in} \mid \frac{\sum_{m\in Neigh(n)} \boldsymbol{s}_m^{out}}{|Neigh(n)|}, \sigma_{aggr}^2\boldsymbol{I})$.
- The decoder module for $[\boldsymbol{z}, \boldsymbol{s}^{in}] \mapsto [\bar{\boldsymbol{x}}^{int}, \bar{\boldsymbol{x}}^{mic}]$ is implemented with a neural ODE network [7]. In other words:

  $$p\big(\bar{\boldsymbol{x}}_n^{int}, \bar{\boldsymbol{x}}_n^{mic} \mid \boldsymbol{z}_n, \boldsymbol{s}_n^{in}\big) = \mathcal{N}\big(. \mid NeuralODENet.forward(\boldsymbol{z}_n, \boldsymbol{s}_n^{in}), \sigma^2\boldsymbol{I}\big).$$

  The choice of a neural ODE here is motivated by Proposition 1 so that we can encourage the mappings to be close to straight paths via flow matching loss. One can simply think of the flow matching objectives as complementary terms to the usual ELBO objective, necessitated by Proposition 1. However, we show that ELBO can subsume the flow matching objective if the generative model and the variational family are designed in a specific way. In particular, we define the following conditional distributions

  $$p\big(\bar{\boldsymbol{x}}_n^{int}(t+dt), \bar{\boldsymbol{x}}_n^{mic}(t+dt) \mid \bar{\boldsymbol{x}}_n^{int}(t), \bar{\boldsymbol{x}}_n^{mic}(t)\big) = \mathcal{N}\Big(. \mid NeuralODENet.vectfield\big(t, \bar{\boldsymbol{x}}_n^{int}(t), \bar{\boldsymbol{x}}_n^{mic}(t)\big), \sigma^2\boldsymbol{I}\Big).$$

  In the above equation $t$ denotes time, $[\bar{\boldsymbol{x}}_n^{int}(0), \bar{\boldsymbol{x}}_n^{mic}(0)]$ equals $[\boldsymbol{z}, \boldsymbol{s}^{in}]$, and $[\bar{\boldsymbol{x}}_n^{int}(1), \bar{\boldsymbol{x}}_n^{mic}(1)]$ equals $[\bar{\boldsymbol{x}}_n^{int}, \bar{\boldsymbol{x}}_n^{mic}]$. Recall that $dim(\boldsymbol{z}) = dim(\boldsymbol{s}^{in}) = dim\big(\bar{\boldsymbol{x}}_n^{int}(t)\big) = dim\big(\bar{\boldsymbol{x}}_n^{mic}(t)\big)$, *i.e.*, at each point in time the vector field is over "$2 \times dim(\boldsymbol{z})$"-dimensional space. Since in the generative model of Supplementary Fig. 17a $\bar{\boldsymbol{x}}_n^{int}$ is not connected to $\boldsymbol{s}^{in}$, the first $dim(\boldsymbol{z})$ dimensions of the vector field are prohibited to depend on $\bar{\boldsymbol{x}}^{mic}(t)$ while the second $dim(\boldsymbol{z})$ dimensions of the vector field depend on both $\bar{\boldsymbol{x}}^{int}(t)$ and $\bar{\boldsymbol{x}}^{mic}(t)$.

- $p(\boldsymbol{x}_n^{int}|\bar{\boldsymbol{x}}_n^{int}) = \mathcal{Z}i\mathcal{N}eg\mathcal{B}in(\boldsymbol{x}_n^{int} \mid \theta_{dec.int}(\bar{\boldsymbol{x}}_n^{int}), \ \theta_{disper.int}, \ p_{0.int})$, where $\mathcal{Z}i\mathcal{N}eg\mathcal{B}in$ is the zero-inflated binomial distribution, $\theta_{dec.int}(.)$ is an MLP decoder that specifies the distribution's mean, $\theta_{disper.int}$ is inverse dispersion, and $p_{0.int}$ is the masking probability.

- Similarly, $p(\boldsymbol{x}_n^{mic}|\bar{\boldsymbol{x}}_n^{mic}) = \mathcal{Z}i\mathcal{N}eg\mathcal{B}in(\boldsymbol{x}_n^{mic} \mid \theta_{dec.mic}(\bar{\boldsymbol{x}}_n^{mic}), \ \theta_{disper.mic}, \ p_{0.mic})$, where $\mathcal{Z}i\mathcal{N}eg\mathcal{B}in$ is the zero-inflated binomial distribution, $\theta_{dec.mic}(.)$ is an MLP decoder that specifies the distribution's mean, $\theta_{disper.mic}$ is inverse dispersion, and $p_{0.mic}$ is the masking probability.

- Finally, $p(\boldsymbol{x}_n|\boldsymbol{x}_n^{int} + \boldsymbol{x}_n^{mic}) = \mathcal{N}(\boldsymbol{x}_n|\boldsymbol{x}_n^{int} + \boldsymbol{x}_n^{mic}, \sigma_{sum}^2\boldsymbol{I})$. Intuitively, this conditional distribution is responsible for encouraging the equality $\boldsymbol{x}_n = \boldsymbol{x}_n^{int} + \boldsymbol{x}_n^{mic}$. Nonetheless, in our implementation the first module of encoder can enforce the aforementioned equality by design, in which case $\sigma_{sum}^2$ is set to zero and the conditional distribution is omitted in practice.

### Supplementary Note 4: Details of the Variational Family

The variational distribution is shown in Supplementary Fig. 17b. As a notational convention we have been denoting the parameters of the generative model by $\theta(.)$, while in the following the variational parameters are denoted by $\varphi(.)$.

- The first stage of encoder is defined as the following conditional distribution:

$$q(\boldsymbol{x}^{mic}_{1:N}, \boldsymbol{x}^{int}_{1:N} \mid \boldsymbol{x}_{1:N}, CT_{1:N}, MCC_{1:N}) = \\ \mathcal{N}(. \mid \varphi_{x\to 2x}(\boldsymbol{x}_{1:N}, CT_{1:N}, MCC_{1:N}), \Sigma_{x\to 2x}(\boldsymbol{x}_{1:N}, CT_{1:N}, MCC_{1:N})),$$

  where $N$ is the number of cells in the subgraph, and $CT_{1:N}$ and $MCC_{1:N}$ denote cell type and microenvironment cell type composition for those $N$ cells, respectively.

- $q(\bar{\boldsymbol{x}}^{int}_n | \boldsymbol{x}^{int}_n) = \mathcal{N}(\bar{\boldsymbol{x}}^{int}_n \mid \varphi_{enc.int}(\boldsymbol{x}^{int}_n), \sigma^2_{enc}\boldsymbol{I})$, where $\varphi_{enc.int}(.)$ is an MLP module.
- Similarly, $q(\bar{\boldsymbol{x}}^{mic}_n | \boldsymbol{x}^{mic}_n) = \mathcal{N}(\bar{\boldsymbol{x}}^{mic}_n \mid \varphi_{enc.mic}(\boldsymbol{x}^{mic}_n), \sigma^2_{enc}\boldsymbol{I})$.
- The conditional distribution for $[\boldsymbol{z}, \boldsymbol{s}^{in}, \boldsymbol{s}^{out}]$ is defined as follows:

$$q(\boldsymbol{z}_{1:N}, \boldsymbol{s}^{in}_{1:N}, \boldsymbol{s}^{out}_{1:N} | \bar{\boldsymbol{x}}^{int}_{1:N}, \bar{\boldsymbol{x}}^{mic}_{1:N}) = \\ \mathcal{N}(. \mid \varphi_{flow0}(\bar{\boldsymbol{x}}^{int}_{1:N}, \bar{\boldsymbol{x}}^{mic}_{1:N}), \sigma_{flow0}(\bar{\boldsymbol{x}}^{int}_{1:N}, \bar{\boldsymbol{x}}^{mic}_{1:N})). \quad (4)$$

  The above conditional distribution has two important facets:

  - **Facet 1**: It is closely related to how bidirectional flow matching[8] defines a single mixture component of its target flow. More precisely, if the mapping $[\boldsymbol{z}, \boldsymbol{s}^{in}] \mapsto [\bar{\boldsymbol{x}}^{int}, \bar{\boldsymbol{x}}^{mic}]$ is trained with a flow matching objective a single mixture component of the target flow can be defined by extending a single sample $[\bar{\boldsymbol{x}}^{int}_n, \bar{\boldsymbol{x}}^{mic}_n]$ to $[\bar{\boldsymbol{x}}^{int}_n, \bar{\boldsymbol{x}}^{mic}_n, \boldsymbol{z}_n, \boldsymbol{s}^{in}_n]$, and this clearly resembles the conditional distribution of Eq. 4.
  - **Facet 2**: The conditional distribution of Eq. 4 is responsible for imposing the constraint $\boldsymbol{s}^{in}_n \approx \frac{1}{Neigh(n)} \sum_{m\in Neigh(n)} \boldsymbol{s}^{out}_m$ among the $\boldsymbol{s}^{in}$ and $\boldsymbol{s}^{out}$ vectors.

  In particular, in Eq. 4 if $\sigma_{flow0} \to \infty$ for $[\boldsymbol{z}, \boldsymbol{s}^{in}]$, then $[\boldsymbol{z}_{1:N}, \boldsymbol{s}^{in}_{1:N}]$ are generated from a Gaussian noise and this resembles the basic form of bidirectional flow matching and "Facet 1" dominates. Nonetheless, in this extreme case "Facet 2" becomes hard to achieve, because the generated $\boldsymbol{s}^{in}$ are totally random. Refer to ablation study of Supplementary Fig. 21 to see the effect of flow matching loss weight and $\sigma_{flow0}$.

- In sum, we explained above how the samples $[\bar{\boldsymbol{x}}^{int}_n, \bar{\boldsymbol{x}}^{mic}_n]$ and $[\boldsymbol{z}_n, \boldsymbol{s}^{in}_n]$ are generated, which - in the lens of flow matching - are samples from the target and base distributions, respectively. Having generated those samples, in the variational family we define the following conditional distribution

$$q(\bar{\boldsymbol{x}}^{int}_n(t), \bar{\boldsymbol{x}}^{int}_n(t) \mid \bar{\boldsymbol{x}}^{int}_n, \bar{\boldsymbol{x}}^{int}_n, \boldsymbol{z}_n, \boldsymbol{s}^{in}_n) = \\ \mathcal{N}\big(. \mid \varphi_{flowt}(\bar{\boldsymbol{x}}^{int}_n, \bar{\boldsymbol{x}}^{mic}_n, \boldsymbol{z}_n, \boldsymbol{s}^{in}_n, t), \sigma_{flowt}(t)^2\boldsymbol{I}\big). \quad (5)$$

  Motivated by the derivations of Supplementary Note 5, we choose $\varphi_{flowt}(.)$ to be a linear interpolation between $[\boldsymbol{z}_n, \boldsymbol{s}^{in}_n]$ and $[\bar{\boldsymbol{x}}^{int}_n, \bar{\boldsymbol{x}}^{mic}_n]$ at $[\bar{\boldsymbol{x}}^{int}_n(t), \bar{\boldsymbol{x}}^{int}_n(t)]$ so as to make the usual

ELBO terms mimic flow matching objective for $0 \leq t \leq 1$. Refer to Supplementary Note 5 to see how the conditional distribution of Eq. 5 is linked to flow matching objective.

### Supplementary Note 5: Linking ELBO and Flow Matching Objective

In this section we show that if the generative model and variational family are designed in a specific way, the usual ELBO objective will encompass the bidirectional flow matching objective. This enables the usage of Bayesian network/inference to model causal relationships while having the benefits of flow-based generative models to achieve, *e.g.*, identifiability as motivated by Proposition 1, or better generative power. We note that previous research has also made connections between likelihood-based and flow-based generative models[9,10].

We first show this connection for the model of Supplementary Fig. 19a, and afterwards we argue that the upper half of the proposed MintFlow model in Supplementary Fig. 17a resembles the model of Supplementary Fig. 19a. In this section our variable naming is totally discordant with the rest of the paper, and is similar to that of denoising diffusion models (DDPM)[11] : In the case of DDPM[11] $\boldsymbol{x}_0$ is a sample from the target distribution, to which a Guassian noise is added until it becomes pure noise $\boldsymbol{x}_T$. In our case the variable $\boldsymbol{x}_T$ can have some parents denoted by $Pa(\boldsymbol{x}_T)$. The variable $\boldsymbol{x}_0$ can be latent and can have some children, but now for simplicity let's assume samples from $\boldsymbol{x}_0$ are observed.

Here we derive the evidence lower-bound for the model/variational distribution of Supplementary Fig. 19. To arrive at terms similar to flow matching, $p(\boldsymbol{x}_{t-1}|\boldsymbol{x}_t)$ terms should be paired with $q(\boldsymbol{x}_{t-1}|\boldsymbol{x}_t, ...)$ terms, but in Supplementary Fig. 19b in the variational distribution we have $q(\boldsymbol{x}_t|\boldsymbol{x}_{t-1}, ...)$ terms. The below lemma enables replacing the $q(\boldsymbol{x}_t|\boldsymbol{x}_{t-1}, ...)$ terms by $q(\boldsymbol{x}_{t-1}|\boldsymbol{x}_t, ...)$ terms. Of note, a similar procedure has been done in the formulation of denoising diffusion model[11].

**Lemma 1** *For $T-1 \geq t > 1$ we have that*

$$q(\boldsymbol{x}_t|\boldsymbol{x}_{t-1}, \boldsymbol{x}_0, \boldsymbol{x}_T) = q(\boldsymbol{x}_{t-1}|\boldsymbol{x}_t, \boldsymbol{x}_0, \boldsymbol{x}_T) \times \frac{q(\boldsymbol{x}_t|\boldsymbol{x}_0, \boldsymbol{x}_T)}{q(\boldsymbol{x}_{t-1}|\boldsymbol{x}_0, \boldsymbol{x}_T)} \tag{6}$$

*The proof follows from basic probability and the definition of conditional probability.*

From Lemma 1 it follows that

$$\begin{aligned}\prod_{t>1}^{T-1} q(\boldsymbol{x}_t|\boldsymbol{x}_{t-1}, \boldsymbol{x}_0, \boldsymbol{x}_T) &= [\prod_{t>1}^{T-1} q(\boldsymbol{x}_{t-1}|\boldsymbol{x}_t, \boldsymbol{x}_0, \boldsymbol{x}_T)] \times [\prod_{t>1}^{T-1} \frac{q(\boldsymbol{x}_t|\boldsymbol{x}_0, \boldsymbol{x}_T)}{q(\boldsymbol{x}_{t-1}|\boldsymbol{x}_0, \boldsymbol{x}_T)}] \\ &= [\prod_{t>1}^{T-1} q(\boldsymbol{x}_{t-1}|\boldsymbol{x}_t, \boldsymbol{x}_0, \boldsymbol{x}_T)] \times \frac{q(\boldsymbol{x}_{T-1}|\boldsymbol{x}_0, \boldsymbol{x}_T)}{q(\boldsymbol{x}_1|\boldsymbol{x}_0, \boldsymbol{x}_T)}.\end{aligned} \tag{7}$$

Now we derive the ELBO as follows

$$\begin{aligned}\mathcal{L} =&\mathbb{E}_q\big[\log p(\boldsymbol{x}_T|Pa(\boldsymbol{x}_T)) + \log p(\boldsymbol{x}_0|\boldsymbol{x}_1)\big] + \mathbb{E}_q\big[\sum_{t>1}^{T}\log p(\boldsymbol{x}_{t-1}|\boldsymbol{x}_t)\big]\\ &-\mathbb{E}_q\big[\sum_{t>1}^{T-1}\log q(\boldsymbol{x}_t|\boldsymbol{x}_{t-1},\boldsymbol{x}_0,\boldsymbol{x}_T)\big]\\ &-\mathbb{E}_q\big[\log q(\boldsymbol{x}_T, Pa(\boldsymbol{x}_T)|\boldsymbol{x}_0)\big] - \mathbb{E}_q\big[\log q\big(\boldsymbol{x}_1|\boldsymbol{x}_0,\boldsymbol{x}_T\big)\big].\end{aligned} \tag{8}$$

Plugging the r.h.s of Eq. 7 in the second line of Eq. 8 we get

$$\begin{aligned}\mathcal{L} =&\mathbb{E}_q\big[\log p(\boldsymbol{x}_T|Pa(\boldsymbol{x}_T)) + \log p(\boldsymbol{x}_0|\boldsymbol{x}_1)\big] + \mathbb{E}_q\big[\sum_{t>1}^{T}\log p(\boldsymbol{x}_{t-1}|\boldsymbol{x}_t)\big]\\ &-\mathbb{E}_q\big[\sum_{t>1}^{T-1}\log q(\boldsymbol{x}_{t-1}|\boldsymbol{x}_t,\boldsymbol{x}_0,\boldsymbol{x}_T)\big] - \mathbb{E}\big[\log q(\boldsymbol{x}_{T-1}|\boldsymbol{x}_0,\boldsymbol{x}_T) - \log q(\boldsymbol{x}_1|\boldsymbol{x}_0,\boldsymbol{x}_T)\big]\\ &-\mathbb{E}_q\big[\log q(\boldsymbol{x}_T, Pa(\boldsymbol{x}_T)|\boldsymbol{x}_0)\big] - \mathbb{E}_q\big[\log q(\boldsymbol{x}_1|\boldsymbol{x}_0,\boldsymbol{x}_T)\big].\end{aligned} \tag{9}$$

Rearranging the terms in Eq. 9, we get

$$\mathcal{L} = \mathbb{E}_q\big[\sum_{t>1}^{T-1}\log p(\boldsymbol{x}_{t-1}|\boldsymbol{x}_t) - \log q(\boldsymbol{x}_{t-1}|\boldsymbol{x}_t,\boldsymbol{x}_0,\boldsymbol{x}_T)\big] \tag{10a}$$

$$+\mathbb{E}_q\big[\log p(\boldsymbol{x}_T|Pa(\boldsymbol{x}_T)) - \log q(\boldsymbol{x}_T, Pa(\boldsymbol{x}_T)|\boldsymbol{x}_0)\big] \tag{10b}$$

$$+\mathbb{E}_q\big[\log p(\boldsymbol{x}_{T-1}|\boldsymbol{x}_T) - \log q(\boldsymbol{x}_{T-1}|\boldsymbol{x}_0,\boldsymbol{x}_T)\big] \tag{10c}$$

$$+\mathbb{E}_q\big[\log p(\boldsymbol{x}_0|\boldsymbol{x}_1)\big]. \tag{10d}$$

At the following we dissect each term of Eq. 10.

The term 10a

The term Eq. 10a turns out to be closely related to the noise prediction objective in denoising diffusion model (*c.f.* Eq. 8 of [11]) and the flow matching objective (*c.f.* Eq. 10 of [8]). Let's assume

$$p(\boldsymbol{x}_{t-1}|\boldsymbol{x}_t) = \mathcal{N}\big(\boldsymbol{x}_{t-1} \mid \boldsymbol{\mu}_{p,t}(\boldsymbol{x}_t), \sigma_p^2\boldsymbol{I}\big), \tag{11a}$$

$$q(\boldsymbol{x}_{t-1}|\boldsymbol{x}_t,\boldsymbol{x}_0,\boldsymbol{x}_T) = \mathcal{N}\big(\boldsymbol{x}_{t-1} \mid \boldsymbol{\mu}_{q,t}(\boldsymbol{x}_t,\boldsymbol{x}_0,\boldsymbol{x}_T), \sigma_q(t)^2\boldsymbol{I}\big). \tag{11b}$$

Each summand of the term 10a is then simplified as

$$\mathbb{E}_q\big[\log p(\boldsymbol{x}_{t-1}|\boldsymbol{x}_t) - \log q(\boldsymbol{x}_{t-1}|\boldsymbol{x}_t,\boldsymbol{x}_0,\boldsymbol{x}_T)\big] \tag{12a}$$

$$= \mathbb{E}_q\Big[KL\big(q(\boldsymbol{x}_{t-1}|\boldsymbol{x}_t,\boldsymbol{x}_0,\boldsymbol{x}_T) \;||\; p(\boldsymbol{x}_{t-1}|\boldsymbol{x}_t)\big)\Big] \tag{12b}$$

$$= \frac{1}{\sigma_q(t)^2}\mathbb{E}_q\big[||\boldsymbol{\mu}_{p,t}(\boldsymbol{x}_t) - \boldsymbol{\mu}_{q,t}(\boldsymbol{x}_t,\boldsymbol{x}_0,\boldsymbol{x}_T)||^2\big]. \tag{12c}$$

The term 12c is similar to the objectives of both diffusion denoising models and flow-based models. More precisely, to arrive at diffusion denoising objective $\boldsymbol{\mu}_{q,t}(.)$ needs to simply ignore

$\boldsymbol{x}_T$ and compute a weighted sum of $\boldsymbol{x}_t$ and $\boldsymbol{x}_0$ (*c.f.* Eq. 7 of [11]). In the limit, the time interval $[0, 1]$ is split to $T \to \infty$ time steps, $\boldsymbol{\mu}_{p,t}(.)$ is the time-dependant residual block of a neural ODE network [7], and $\boldsymbol{\mu}_{q,t}(.)$ is the vector field of the target flow conditioned on $\boldsymbol{x}_0$ and $\boldsymbol{x}_T$. The function $\boldsymbol{\mu}_{q,t}(.)$ is usually a linear interpolator at $\boldsymbol{x}_t$ given $\boldsymbol{x}_0$ and $\boldsymbol{x}_T$. Here $\boldsymbol{x}_0$ and $\boldsymbol{x}_T$ correspond to the conditions in the target flow in conditional flow matching, and different choices of $q(\boldsymbol{x}_T|\boldsymbol{x}_0)$ determines how $\boldsymbol{x}_T$ is generated given $\boldsymbol{x}_0$. In other words, $q(\boldsymbol{x}_T|\boldsymbol{x}_0)$ determines how given the data point $\boldsymbol{x}_0$ a single mixture component of the target flow is created in the corresponding flow matching objective.

The term 10b

The term 10b is crucial in connecting the flow matching block to the rest of the generative model. Because this term implies that the generated samples from $q(\boldsymbol{x}_T, Pa(\boldsymbol{x}_T)|\boldsymbol{x}_0)$ have to comply to $p(\boldsymbol{x}_T|Pa(\boldsymbol{x}_T)$ in the original generative model.

The terms 10c and 10d

Notice that the term 10c is the equivalent of the term 10a for the boundary condition $t = T$. Moreover, the term 10d is the last step to generate a sample from the target distribution. This step can be treated differently since the generated sample takes, *e.g.*, discrete or count values (*c.f.* Eq. 13 of [11]). All in all, the terms 10c and 10d are the equivalent of 10b at the boundaries $t = 0$ and $t = T$.

Relation of the general model of Supplementary Fig. 19 to the proposed MintFlow model

Having discussed the relation between ELBO and flow matching objective for the model of Supplementary Fig. 19, we now move on to explain why the upper-half of the proposed MintFlow model in Supplementary Fig. 17a is similar to the model of Supplementary Fig. 19a, thus the derivations of this section apply to the proposed MintFlow model. More specifically, in the proposed generative model, $\boldsymbol{x}_T$ of Supplementary Fig. 19a corresponds to the vector $[\boldsymbol{z}, \boldsymbol{s}^{in}]$ in Supplementary Fig. 17a. Moreover, $Pa(\boldsymbol{x}_T)$ in Supplementary Fig. 19 corresponds to the vector $[\boldsymbol{t}, \boldsymbol{s}^{out}]$ in Supplementary Fig. 17a because in the latter figure, $\boldsymbol{z}$ and $\boldsymbol{s}^{in}$ are conditioned on $\boldsymbol{t}$ and $\boldsymbol{s}^{out}$, respectively.

Here we reiterate the importance of the term 10b in the proposed model. In the proposed generative model the term $p(\boldsymbol{s}_n^{in}|Pa(\boldsymbol{s}_n^{in}))$ is designed to ensure $\boldsymbol{s}_n^{in} \approx \frac{1}{Niegh(n)} \sum_{m \in Neigh(n)} \boldsymbol{s}_m^{out}$. In our objective, the term 10b asserts that the generated samples from $q(.)$ comply with the intended relation between $\boldsymbol{s}^{in}$ and the $\boldsymbol{s}^{out}$ vectors of the neighbouring cells.

### Supplementary Note 6: Detailed Datasets Description

**Simulated data**. We used the data simulation code from NicheCompass[12] to generate a tissue with eight regions comprising 10,000 cells and 2,000 simulated genes. Each region has varying MCCs (Supplementary Fig. 1). In the experiments presented in Supplementary Fig. 3, we excluded regions with MCCs consisting of only one cell type as this is an ill-posed setup where the MCC and CT labels are the same and thus the separation of intrinsic and microenvironment-induced effects could be harder (Methods, $Objective_1$ and $Objective_2$).

**Atopic dermatitis**. Atopic dermatitis data was newly generated using the 10x Genomics Xenium in situ 5k-plex platform. We used non-lesional and lesional human adult skin tissue from adult individuals with atopic dermatitis. All research ethics committees and regulatory approvals were in place for the collection and storage of research samples at St John's Institute of Dermatology, Guy's Hospital, London (REC reference number: EC00/128). Fresh frozen OCT-embedded skin samples were sectioned at 10 μm thickness directly onto the 10X Genomics xenium slide kept at -20°C. We then ran the slides through the 10x Genomics Xenium prime in situ gene expression protocol and imaged using the 10X Genomics Xenium Analyzer. This allowed us to probe 5000 genes at subcellular spatial resolution in human skin sections. We included protein staining to facilitate cell segmentation. Following imaging, the same slides were then used to generate H&E staining and imaged at ×20 magnification on a Hamamatsu Nanozoomer s60.

10x Genomics Xenium data was filtered to exclude cells with <10 genes per cell. For initial cell annotation, integration of all cells was performed using scVI using raw counts as previously described[13]. We constructed a kNN graph (k=20) using the scVI embedding. We performed community detection (Leiden algorithm) with resolution 0.1 and then performed hierarchical clustering on broad populations. We assigned cell annotations based on DEGs, spatial location of populations, and our previous skin marker genes[13]. To provide further confidence in our annotations, we also performed automated annotation using 1) a CellTypist model trained on scRNA-seq data[14] and 2) integrative label transfer using scanVI with scRNA-seq data[14]. For normalization of gene expression counts, we used the shifted log transformation (target 10,000).

For gene set enrichment analysis for the *Melanoma* MGPs and the *T_DC* MGP, we used GSEAPY[15] with the relevant MGPs as input and the GO_Biological_Process_2023 gene set. We used the top 500 differentially expressed genes per fibroblast subtype. A cut-off statistical significance of 0.01 was used.

For gene module scoring of the melanoma *Stromal* MGP across diseases, we scored the *T_DC* MGP in scRNA-seq data across skin diseases previously assembled in [13] using scanpy.tl.score_genes. We used the same function for scoring the *T_DC* MGP in the cross-tissue atlas we previously assembled [13].

For external validation of the *T_DC* MGP, we used scanpy.tl.score_genes to calculate scores for the MGP in scRNA-seq data for individual skin samples from donors with atopic dermatitis in paired lesional and non-lesional samples.

Drug2cell scores were generated using drugcell.score. We then used the scanpy rank_genes_groups function on the generated adata.uns['drug2cell'] object with the Wilcoxon method.

We used CellPhoneDB v5 (method 2) for cell-cell communication analysis. We used our previously published scRNA-seq data[14] combined with public datasets from skin for more granular immune cell annotations. We restricted interactions to chemokines from the *T_DC* MGP. We visualized the results using ktplotspy.

***Clear cell renal carcinoma.*** ccRCC data was newly generated from a single tumor nephrectomy, including three macroscopically distinct regions of the tumour core, as well as the tumour–normal interface where the cancer abuts non-neoplastic kidney tissue, using the 10x Genomics Xenium *in situ* 5k-plex platform. The clinical data of this patient (identifier: PD43284) is provided in a previous study[16], and samples were collected under the ethics of the DIAMOND study; Evaluation of biomarkers in urological disease (NHS National Research Ethics Service ref. 03/018). Fresh frozen OCT-embedded skin samples were sectioned at 10 μm thickness directly onto the 10X Genomics xenium slide at -20°C before running the 10x Genomics Xenium prime in situ gene expression protocol and imaging using the 10X Genomics Xenium Analyzer. Protein staining was used to facilitate cell segmentation, and the same slides were then used to generate H&E staining and imaged at ×20 magnification on a Hamamatsu Nanozoomer s60. Log normalization of gene expression counts was performed and scVI was used for integration of batches[13], before unsupervised clustering using the Leiden algorithm. Cell types were assigned at three hierarchical levels based on DEGs and the spatial location of each cluster. The annotated data were used to train MintFlow according to training parameters specified in config files (Supplementary Table 1). Regions of interest across Xenium-profiled tissue sections were identified using the microenvironment score, using normalized and log-transformed count matrices derived from both $X$ and $X^{mic}$ generated by the MintFlow pipeline. For downstream analyses using MintFlow outputs, the data were subsetted into T cells, macrophages, endothelial cells and fibroblasts. $X^{mic}$ was derived for each dataset, before normalization and log-transformation, PCA, neighborhood graph construction and UMAP embedding. Thereafter, Leiden clustering was performed, with sc.tl.score_genes used to derive DEGs, corresponding to a MGP for each clustering and thus resolving microenvironment-induced states for each cell type. To identify T cell microenvironment-induced states within previously published scRNA-seq data[16], we used scanpy.tl.score_genes on the annotated and filtered count matrix from the scRNA-seq dataset, utilising MGPs from each microenvironment-induced cell state for T cells identified using MintFlow. T cells from each patient were also scored according to the MintFlow-derived MGPs, and patients were compared using a heatmap, grouped by Leibovich clinical risk score provided in the original dataset[16]. Ligand-receptor interaction analysis was performed using the LIANA+ package[17] in "rank aggregate" mode, partitioning T cells based on ‘TLS’ or ‘non-TLS’ based on scanpy.tl.score_genes. For *in silico* perturbation, a polygon was selected around the TLS region from the relevant tumour core, before removal of TLS macrophages. MGPs were recomputed for each T cell microenvironment-induced state after perturbation, and compared to unmodified outputs from MintFlow. For survival analysis, we derived TCGA data from clinically annotated bulk RNA-seq data of ccRCC, downloading the data from the Xena platform as previously described[18]. Survival analysis was then performed in R, using the *survival* and *survminer* packages, using the median expression of MGP genes derived from MintFlow to stratify patients into ‘high score’ or ‘low score’. Kaplan-Meier plots were then generated using the *ggsurvplot* function, plotting overall survival (OS) against time (days) to event.

### Supplementary Tables

| Experiment | number of tissue sections | number of neighbors in spatial neighborhood graph | window width for the customized sampler for PyTorch Geometric's neighbor loader | number of graph hops | dimension of embeddings | lower-bound on the encoder variance for non-central nodes (Methods, Scalability) | coefficient of flow matching objectives i.e. objective 5 and objective 6 |
|---|---|---|---|---|---|---|---|
| | **Hyperparameter** | | | | | | |
| Fig. 1g (simulated) | 1 | 5 | 500 | 1 | 20 | 0.1 | 0.0 |
| Fig. 1h (left, eczema 1 section) | 1 | 5 | 800 | 1 | 10 | 0.1 | 10 |
| Fig. 1h (middle, melanoma) | 1 | 10 | 400 | 1 | 10 | 0.1 | 10 |
| Fig. 3 and 1h (right, eczema 10 sections) | 10 | 5 | Varying, in {600, 800, 900, 120} | 1 | 100 | 0.1 | 0.0 |
| Fig. 4 (melanoma) | 1 | 5 | 400 | 1 | 100 | 0.1 | 0.0 |
| Fig. 5,6 (kidney) | 4 | 5 | 850 | 1 | 100 | 0.1 | 0.0 |
| Ablation study on simulated data (Supplementary Fig. 18) | 1 | 5 | 500 | 1 | 100 | 0.1 | Varying in {0.01, 0.1, 0.2, 0.5, 1.0, 10.0} |

**Supplementary Table 1 (part 1) |** MintFlow's parameter settings in our experiments.

| Experiment | flow matching, scale of target flow noise | ODE solver number of time steps | coefficient of Objective 1, i.e. CT being predictable from intrinsic embedding (Methods) | coefficient of Objective 2, i.e. MCC being predictable from microenvironment embedding (Methods) | coefficient of Objective 3, i.e. CT being predictable from encoded intrinsic component of expression (Methods) | coefficient of Objective 4, i.e. MCC being predictable from encoded microenv-induced component of expression (Methods) | coefficient of Objective 7, i.e. MCC not being predictable from intrinsic embedding (Methods) |
|---|---|---|---|---|---|---|---|
| | **Hyperparameter** | | | | | | |
| Fig. 1g (simulated) | 0.01 | 10 | 0.1 | 0.1 | 0 | 0 | 1.0 |
| Fig. 1h (left, eczema 1 section) | 0.01 | 10 | 1.0 | 1.0 | 1.0 | 1.0 | 1.0 |
| Fig. 1h (middle, melanoma) | 0.01 | 10 | 1.0 | 1.0 | 1.0 | 1.0 | 1.0 |
| Fig. 3 and 1h (right, eczema 10 sections) | 0.01 | 10 | 1.0 | 1.0 | 1.0 | 1.0 | 1.0 |
| Fig. 4 (melanoma) | 0.01 | 10 | 0.1 | 0.1 | 0.0 | 0.0 | 1.0 |
| Fig. 5,6 (kidney) | 0.01 | 10 | 0.1 | 0.1 | 1.0 | 1.0 | 1.0 |
| Ablation study on simulated data (Supplementary Fig. 18) | 0.01 | 10 | 0.1 | 0.1 | 0.0 | 0.0 | 1.0 |

**Supplementary Table 1 (part 2) |** MintFlow's parameter settings in our experiments.

| Experiment | coefficient of Objective 8, i.e. MCC not being predictable from encoded intrinsic component of expression (Methods) | coefficient of batch mixing objective on encoded intrinsic component of expression (Methods) | coefficient of batch mixing objective on encoded microenv-induced component of expression (Methods) | coefficient of objective 9, i.e. the intrinsic vector of cells with the same cell type label being close | coefficient of objective 10, i.e. the embedded intrinsic component of expression of cells with the same cell type label being close | coefficient of objective 11, i.e. the intrinsic component of expression of cells with the same cell type label being close | target-sum in scanpy.pp.normalize_total |
|---|---|---|---|---|---|---|---|
| | **Hyperparameter** | | | | | | |
| Fig. 1g (simulated) | 1.0 | 0 | 0 | 100.0 | 100.0 | 100.0 | 10000 |
| Fig. 1h (left, eczema 1 section) | 1.0 | 0 | 0 | 100.0 | 100.0 | 100.0 | 10000 |
| Fig. 1h (middle, melanoma) | 1.0 | 0 | 0 | 100.0 | 100.0 | 100.0 | 10000 |
| Fig. 3 and 1h (right, eczema 10 sections) | 1.0 | 1.0 | 0.01 | 100.0 | 100.0 | 100.0 | 10000 |
| Fig. 4 (melanoma) | 1.0 | 0 | 0 | 100.0 | 100.0 | 100.0 | 10000 |
| Fig. 5,6 (kidney) | 1.0 | 0.01 | 0.01 | 100.0 | 100.0 | 100.0 | 10000 |
| Ablation study on simulated data (Supplementary Fig. 18) | 1.0 | 0 | 0 | 100.0 | 100.0 | 100.0 | 10000 |

**Supplementary Table 1 (part 3) |** MintFlow's parameter settings in our experiments.

| Experiment | Hyperparameter: parameters of the MCC-conditioned prior on microenvironment embeddings kept fixed during training | type of optimizer | learning rate of optimizer | number of epochs | at the beginning of each epoch, number of updates to the Wasserstein distance estimators | size of each mini-batch when updating the Wasserstein distance estimators at the beginning of each epoch | during training, the number of accumulated gradient steps (i.e. backward passes) before each update to model parameters |
|---|---|---|---|---|---|---|---|
| Fig. 1g (simulated) | True | Adam | 0.001 | 100 | 500 | 250 | 5 |
| Fig. 1h (left, eczema 1 section) | True | Adam | 0.001 | 100 | 10 | 50 | 2 |
| Fig. 1h (middle, melanoma) | True | Adam | 0.001 | 100 | 10 | 50 | 2 |
| Fig. 3 and 1h (right, eczema 10 sections) | True | Adam | 0.001 | 20 | 2000 | 50 | 2 |
| Fig. 4 (melanoma) | True | Adam | 0.001 | 50 | 500 | 250 | 2 |
| Fig. 5,6 (kidney) | True | Adam | 0.001 | 50 | 2000 | 50 | 2 |
| Ablation study on simulated data (Supplementary Fig. 18) | True | Adam | 0.001 | 30 | 2000 | 50 | 5 |

**Supplementary Table 1 (part 4) |** MintFlow's parameter settings in our experiments.

| Experiment | during training, the number of updates to Wasserstein distance estimators, after each update to model parameters | initial annealing coefficient for the reconstruction loss of intrinsic and microenvironment components of expression (Methods) | fraction of initial epochs where the annealing coefficient of reconstruction loss is fixed to the initial value (i.e. prev. column) | fraction of epochs where the annealing coefficient for reconstruction loss is linearly increased to its maximum value (Methods) | final annealing coefficient for the reconstruction loss of intrinsic and microenvironment components of expression (Methods) | fraction of final epochs where the annealing coefficient of reconstruction loss is fixed to the final value (i.e. prev. column) | ODE solver |
|---|---|---|---|---|---|---|---|
| Fig. 1g (simulated) | 50 | 1.0 | 100.0 | 0.0 | 1.0 | 0.0 | dopri5 |
| Fig. 1h (left, eczema 1 section) | 3 | 0.00001 | 0.5 | 0.2 | 0.001 | 0.3 | dopri5 |
| Fig. 1h (middle, melanoma) | 3 | 0.00001 | 0.5 | 0.2 | 0.001 | 0.3 | dopri5 |
| Fig. 3 and 1h (right, eczema 10 sections) | 20 | 0.0001 | 0.5 | 0.2 | 0.001 | 0.3 | dopri5 |
| Fig. 4 (melanoma) | 10 | 0.00001 | 0.5 | 0.2 | 0.001 | 0.3 | dopri5 |
| Fig. 5,6 (kidney) | 20 | 0.00001 | 0.5 | 0.2 | 0.1 | 0.3 | dopri5 |
| Ablation study on simulated data (Supplementary Fig. 18) | 10 | 0.00001 | 0.5 | 0.2 | 0.1 | 0.3 | dopri5 |

(Column group heading: **Hyperparameter**)

**Supplementary Table 1 (part 5) |** MintFlow's parameter settings in our experiments.

#### References

1. Khemakhem, I., Kingma, D. P., Monti, R. P. & Hyvärinen, A. Variational Autoencoders and Nonlinear ICA: A Unifying Framework. *arXiv [stat.ML]* (2019).
2. Velten, B. *et al.* Identifying temporal and spatial patterns of variation from multimodal data using MEFISTO. *Nat. Methods* **19**, 179–186 (2022).
3. Dong, M., Su, D. G., Kluger, H., Fan, R. & Kluger, Y. SIMVI disentangles intrinsic and spatial-induced cellular states in spatial omics data. *Nat. Commun.* **16**, 2990 (2025).
4. Zhao, S., Song, J. & Ermon, S. InfoVAE: Balancing Learning and Inference in Variational Autoencoders. *Proc. Conf. AAAI Artif. Intell.* **33**, 5885–5892 (2019).
5. Jerby-Arnon, L. & Regev, A. DIALOGUE maps multicellular programs in tissue from single-cell or spatial transcriptomics data. *Nat. Biotechnol.* (2022) doi:10.1038/s41587-022-01288-0.
6. Fischer, D. S., Schaar, A. C. & Theis, F. J. Modeling intercellular communication in tissues using spatial graphs of cells. *Nat. Biotechnol.* (2022) doi:10.1038/s41587-022-01467-z.
7. Chen, R. T. Q., Rubanova, Y., Bettencourt, J. & Duvenaud, D. Neural ordinary differential equations. *arXiv [cs.LG]* (2018).
8. Tong, A. *et al.* Improving and generalizing flow-based generative models with minibatch optimal transport. *arXiv [cs.LG]* (2023).
9. Kingma, D. P., Salimans, T., Poole, B. & Ho, J. Variational Diffusion Models. *arXiv [cs.LG]* (2021).
10. Vincent, P. A connection between score matching and denoising autoencoders. *Neural Comput.* **23**, 1661–1674 (2011).
11. Ho, J., Jain, A. & Abbeel, P. Denoising Diffusion Probabilistic Models. *arXiv [cs.LG]* (2020).
12. Birk, S. *et al.* Quantitative characterization of cell niches in spatially resolved omics data. *Nat. Genet.* (2025) doi:10.1038/s41588-025-02120-6.
13. Steele, L. *et al.* A single cell and spatial genomics atlas of human skin fibroblasts in health and disease. *bioRxiv* 2024.12.23.629194 (2024) doi:10.1101/2024.12.23.629194.
14. Reynolds, G. *et al.* Developmental cell programs are co-opted in inflammatory skin disease. *Science* **371**, eaba6500 (2021).

15. Fang, Z., Liu, X. & Peltz, G. GSEApy: a comprehensive package for performing gene set enrichment analysis in Python. *Bioinformatics* **39**, btac757 (2023).
16. Li, R. *et al.* Mapping single-cell transcriptomes in the intra-tumoral and associated territories of kidney cancer. *Cancer Cell* **40**, 1583–1599.e10 (2022).
17. Dimitrov, D. *et al.* LIANA+ provides an all-in-one framework for cell-cell communication inference. *Nat. Cell Biol.* **26**, 1613–1622 (2024).
18. Jammihal, T. *et al.* Immunogenomic determinants of exceptional response to immune checkpoint inhibition in renal cell carcinoma. *Nat. Cancer* **6**, 372–384 (2025).
